# Tissue-Specific Alteration of Metabolic Pathways Influences Glycemic Regulation

**DOI:** 10.1101/790618

**Authors:** Natasha H. J. Ng, Sara M. Willems, Juan Fernandez, Rebecca S. Fine, Eleanor Wheeler, Jennifer Wessel, Hidetoshi Kitajima, Gaelle Marenne, Jana K. Rundle, Xueling Sim, Hanieh Yeghootkar, Nicola L. Beer, Anne Raimondo, Andrei I. Tarasov, Soren K. Thomsen, Martijn van de Bunt, Shuai Wang, Sai Chen, Yuning Chen, Yii-Der Ida Chen, Hugoline G. de Haan, Niels Grarup, Ruifang Li-Gao, Tibor V. Varga, Jennifer L Asimit, Shuang Feng, Rona J. Strawbridge, Erica L. Kleinbrink, Tarunveer S. Ahluwalia, Ping An, Emil V. Appel, Dan E Arking, Juha Auvinen, Lawrence F. Bielak, Nathan A. Bihlmeyer, Jette Bork-Jensen, Jennifer A. Brody, Archie Campbell, Audrey Y Chu, Gail Davies, Ayse Demirkan, James S. Floyd, Franco Giulianini, Xiuqing Guo, Stefan Gustafsson, Benoit Hastoy, Anne U. Jackson, Johanna Jakobsdottir, Marjo-Riitta Jarvelin, Richard A. Jensen, Stavroula Kanoni, Sirkka Keinanen-Kiukaanniemi, Jin Li, Man Li, Kurt Lohman, Yingchang Lu, Jian’an Luan, Alisa K. Manning, Jonathan Marten, Carola Marzi, Karina Meidtner, Dennis O. Mook-Kanamori, Taulant Muka, Giorgio Pistis, Bram Prins, Kenneth M. Rice, Neil Robertson, Serena Sanna, Yuan Shi, Albert Vernon Smith, Jennifer A. Smith, Lorraine Southam, Heather M. Stringham, Salman M. Tajuddin, Vinicius Tragante, Sander W. van der Laan, Helen R. Warren, Jie Yao, Andrianos M. Yiorkas, Weihua Zhang, Wei Zhao, Emma Ahlqvist, Mariaelisa Graff, Heather M. Highland, Anne E Justice, Ken Sin Lo, Eirini Marouli, Carolina Medina-Gomez, Saima Afaq, Wesam A Alhejily, Najaf Amin, Folkert W. Asselbergs, Lori L. Bonnycastle, Michiel L. Bots, Ivan Brandslund, Ji Chen, Cramer Christensen, John Danesh, Renée de Mutsert, Abbas Dehghan, Tapani Ebeling, Paul Elliott, EPIC-InterAct Consortium, Aliki-Eleni Farmaki, Jessica D. Faul, Paul W. Franks, Steve Franks, Andreas Fritsche, Anette P. Gjesing, Mark O. Goodarzi, Vilmundur Gudnason, Göran Hallmans, Tamara B. Harris, Karl-Heinz Herzig, Marie-France Hivert, Jan-Håkan Jansson, Min A Jhun, Torben Jørgensen, Marit E. Jørgensen, Pekka Jousilahti, Eero Kajantie, Maria Karaleftheri, Sharon L.R. Kardia, Leena Kinnunen, Heikki A. Koistinen, Pirjo Komulainen, Peter Kovacs, Johanna Kuusisto, Markku Laakso, Leslie A. Lange, Lenore J. Launer, Jung-Jin Lee, Aaron Leong, Jaana Lindström, Jocelyn E. Manning Fox, Satu Männistö, Nisa M Maruthur, Leena Moilanen, Antonella Mulas, Mike A. Nalls, Matthew Neville, James S. Pankow, Alison Pattie, Eva R.B. Petersen, Hannu Puolijoki, Asif Rasheed, Paul Redmond, Frida Renström, Michael Roden, Danish Saleheen, Juha Saltevo, Kai Savonen, Sylvain Sebert, Tea Skaaby, Kerrin S Small, Alena Stančáková, Jakob Stokholm, Konstantin Strauch, E-Shyong Tai, Kent D. Taylor, Betina H. Thuesen, Anke Tönjes, Emmanouil Tsafantakis, Tiinamaija Tuomi, Jaakko Tuomilehto, Understanding Society Scientific Group, Matti Uusitupa, Marja Vääräsmäki, Ilonca Vaartjes, Magdalena Zoledziewska, Goncalo Abecasis, Beverley Balkau, Hans Bisgaard, Alexandra I. Blakemore, Matthias Blüher, Heiner Boeing, Eric Boerwinkle, Klaus Bønnelykke, Erwin P. Bottinger, Mark J. Caulfield, John C Chambers, Daniel I Chasman, Ching-Yu Cheng, Anne Clark, Francis S. Collins, Josef Coresh, Francesco Cucca, Gert J. de Borst, Ian J. Deary, George Dedoussis, Panos Deloukas, Hester M. den Ruijter, Josée Dupuis, Michele K. Evans, Ele Ferrannini, Oscar H Franco, Harald Grallert, Leif Groop, Torben Hansen, Andrew T. Hattersley, Caroline Hayward, Joel N. Hirschhorn, Arfan Ikram, Erik Ingelsson, Fredrik Karpe, Kay-Tee Kaw, Wieland Kiess, Jaspal S Kooner, Antje Körner, Timo Lakka, Claudia Langenberg, Lars Lind, Cecilia M Lindgren, Allan Linneberg, Leonard Lipovich, Ching-Ti Liu, Jun Liu, Yongmei Liu, Ruth J.F. Loos, Patrick E. MacDonald, Karen L. Mohlke, Andrew D Morris, Patricia B. Munroe, Alison Murray, Sandosh Padmanabhan, Colin N A Palmer, Gerard Pasterkamp, Oluf Pedersen, Patricia A. Peyser, Ozren Polasek, David Porteous, Michael A. Province, Bruce M Psaty, Rainer Rauramaa, Paul M Ridker, Olov Rolandsson, Patrik Rorsman, Frits R. Rosendaal, Igor Rudan, Veikko Salomaa, Matthias B. Schulze, Robert Sladek, Blair H Smith, Timothy D Spector, John M. Starr, Michael Stumvoll, Cornelia M van Duijn, Mark Walker, Nick J. Wareham, David R. Weir, James G. Wilson, Tien Yin Wong, Eleftheria Zeggini, Alan B. Zonderman, Jerome I. Rotter, Andrew P. Morris, Michael Boehnke, Jose Florez, Mark I McCarthy, James B Meigs, Anubha Mahajan, Robert A. Scott, Anna L Gloyn, Inês Barroso

## Abstract

Metabolic dysregulation in multiple tissues alters glucose homeostasis and influences risk for type 2 diabetes (T2D). To identify pathways and tissues influencing T2D-relevant glycemic traits (fasting glucose [FG], fasting insulin [FI], two-hour glucose [2hGlu] and glycated hemoglobin [HbA1c]), we investigated associations of exome-array variants in up to 144,060 individuals without diabetes of multiple ancestries. Single-variant analyses identified novel associations at 21 coding variants in 18 novel loci, whilst gene-based tests revealed signals at two genes, *TF* (HbA1c) and *G6PC* (FG, FI). Pathway and tissue enrichment analyses of trait-associated transcripts confirmed the importance of liver and kidney for FI and pancreatic islets for FG regulation, implicated adipose tissue in FI and the gut in 2hGlu, and suggested a role for the non-endocrine pancreas in glucose homeostasis. Functional studies demonstrated that a novel FG/FI association at the liver-enriched *G6PC* transcript was driven by multiple rare loss-of-function variants. The FG/HbA1c-associated, islet-specific *G6PC2* transcript also contained multiple rare functional variants, including two alleles within the same codon with divergent effects on glucose levels. Our findings highlight the value of integrating genomic and functional data to maximize biological inference.

**Highlights:**

- 23 novel coding variant associations (single-point and gene-based) for glycemic traits
- 51 effector transcripts highlighted different pathway/tissue signatures for each trait
- The exocrine pancreas and gut influence fasting and 2h glucose, respectively
- Multiple variants in liver-enriched G6PC and islet-specific G6PC2 influence glycemia

## Introduction

It has long been recognized that rare and penetrant disease-causing mutations can pinpoint key proteins and pathways involved in human metabolism (Froguel et al., 1992; Gloyn et al., 2004; Montague et al., 1997). Type 2 diabetes (T2D) results from an inability of the pancreatic islet beta cells to produce and secrete sufficient insulin, compounded by the failure of metabolic tissues to respond to insulin and store glucose appropriately. Blood glucose levels are regulated by the co-ordination of homeostatic pathways operating across multiple tissues that control metabolism, therefore a clearer understanding of their relative roles is critical in guiding efforts to modulate them pharmacologically to treat T2D and pre-diabetes. In recent years, technological advances have made it possible to assay genetic variation genome-wide and at scale. These provide tremendous opportunities to understand metabolic differences within the physiological range through the study of quantitative fasting and post-challenge glycemic measures (Mahajan et al., 2015; Scott et al., 2012; Wessel et al., 2015; Wheeler et al., 2017a). These measures can influence the risk of developing pathophysiological conditions such as T2D and cardiovascular disease. However, as in all genome-wide association studies (GWAS), it has proven challenging to translate the associated genetic signals into biological pathways, as the vast majority of association signals lie within non-coding regions, and connecting them to their respective effector genes is less straightforward. There are to date over 97 loci reported to be associated with glycemic traits, across different genetic approaches (Wheeler et al., 2017b). One approach to facilitate identification of likely causal variants and transcripts is to focus on coding variation, whose effects on protein sequence can be predicted and functionally tested, facilitating identification of likely causal genes and the ensuing biological insights. This strategy has been successfully used to establish not only the effector genes but also the direction of effect of T2D risk alleles on protein function such as in the case of SLC30A8 (Flannick et al., 2014) and PAM (Steinthorsdottir et al., 2014; Thomsen et al., 2018).

Here, we describe the largest exome-array study to date across four commonly-used glycemic traits (fasting glucose [FG], fasting insulin [FI], glycated hemoglobin [HbA1c], and two-hour glucose [2hGlu]) in up to 144,060 non-diabetic individuals from multiple ancestries, to discover variants and loci influencing these traits within the physiological range. We sought to identify causal variants and putative effector transcripts in known and novel loci, and subsequently highlight pathways and tissues that are enriched for these glycemic trait associations. We further complemented our analyses with functional validation of selected effector transcripts, focusing on novel FG/FI locus *G6PC* and FG/HbA1c locus *G6PC2*, to establish functional links between the associated rare coding variants in those loci and glucose regulation through different metabolic tissues. Together, our findings provide valuable insight into the biology underlying glycemic traits, and build on the knowledge required for validating candidate genes for therapeutic targeting in diabetes.

## Results

### Identification of coding variant and gene-based glycemic trait associations

We focused on coding variants on the exome chip as these could point more directly to their potential effector transcripts (i.e. likely causal gene[s]). Single-variant and gene-based association analyses with FG, FI, HbA1c, and 2hGlu levels were performed on exome-array coding variants in up to 144,060 individuals without diabetes of European (85%), African-American (6%), South Asian (5%), East Asian (2%), and Hispanic (2%) ancestry from up to 64 cohorts (**Table S1, Methods**).

We performed single-variant analyses in each individual cohort using a linear mixed model and combined results by fixed-effect meta-analyses within and across ancestries. As body mass index (BMI) is a major risk factor for T2D and is correlated with glycemic traits, all analyses were adjusted for BMI (**Methods**) to identify loci influencing glycemia independently from their effects on overall adiposity. We used distance-based clumping and considered signals to be novel, if they were located more than 500 kb from a variant with an established association with any of the glycemic traits or T2D at the time of the study (**Methods**).

Based on the above definition, we found 21 coding variants (in 18 genes) which were not previously associated with any other glycemic trait or T2D risk, that are now associated at exome-wide significance (defined as *P*<2.2 × 10^−7^) (Mahajan et al., 2018b; Sveinbjornsson et al., 2016) with their respective glycemic trait(s) (**Table 1, Methods**). Among these novel loci were a missense (p.E1365D) and splice region variant in *OBSL1* associated with FI, another missense variant (p.L300P) in *RAPGEF3* associated with FI, a missense variant (p.S439N) in *SPTB* associated with HbA1c, and missense variants p.R187Q in *ANKH* and p.R456Q in *STEAP2* that are associated with FG (**Table 1**). *OBSL1* encodes a cytoskeletal protein related to obscurin, mutations in which have been shown to lead to an autosomal recessive primordial growth disorder (OMIM: 612921). Loss of OBSL1 leads to downregulation of CUL7, a protein known to interact with IRS-1, downstream of the insulin receptor signaling pathway (Hanson et al., 2009). *RAPGEF3* encodes a cAMP-regulated guanine nucleotide exchange factor and is part of a cAMP-responsive signalling complex. The gene has been shown to be involved in cAMP-dependent adipogenesis (Jia et al., 2012), and investigation of associations in other related traits showed that the same *RAPGEF3* variant is also associated with BMI, waist-hip ratio and height (all *P*<1 × 10^−4^; **Tables 1 and S2**). This suggests that its role in adiposity and obesity is likely to link it to FI regulation. Since the directions of effect of the variant are opposite for FI and BMI, the observed association could however be due to collider bias and should thus be interpreted with caution. *SPTB* encodes the protein spectrin beta, which is a major constituent of the cytoskeletal network underlying the erythrocyte plasma membrane. Mutations in this gene underlie a range of hematological disorders such as hemolytic anemias (OMIM: 617948, 616649). Given that red blood cell disorders can interfere with HbA1c levels (Wheeler et al., 2017a), this missense variant identifies *SPTB* as the likely effector transcript at this locus. *ANKH* encodes a transmembrane protein likely acting as a transporter. Recently, the FG-lowering allele reported here was shown to associate with decreased T2D risk in Europeans (OR=0.78 [0.69-0.87], *P*_EUR_=2.0 × 10^−7^), and had a >97% posterior probability of being causal, suggesting that this gene is the effector transcript at this locus (Mahajan et al., 2018b). In the final example, *STEAP2* encodes a six transmembrane protein localized both intracellularly and on the plasma membrane, and is suggested to have roles in the regulation of iron transport (Sikkeland et al., 2016). A closely-related member of the *STEAP* family, *STEAP4*, has been reported to mediate cellular response to inflammatory stress through its role as a metalloreductase mediating iron and copper homeostasis (Scarl et al., 2017). Though little is known about *STEAP2* function, in the recent T2D analysis, the FG-associated variant in *STEAP2* was also found to be nominally associated with T2D risk (*P*<1 × 10^−4^) (Mahajan et al., 2018b) (**Tables 1 and S2**). In addition to the novel loci, 53 other significant coding variant associations (in 40 genes) were detected that were within 500 kb of an established glycemic GWAS locus. These were of interest as they could point to a causal gene (**Tables 1 and S3**).

**Table 1.** Single-point coding variant associations meeting the significant threshold for coding variants of *P*<2.2 x 10^−7^. This table includes all novel coding variants meeting this threshold, irrespective of whether they fall in completely new loci or in previously-established loci, provided that the association at the established locus was not shown to be due to a non-coding variant (Table S3) or another coding variant at the same locus. Novel loci are highlighted in bold. HbA1c: glycated haemoglobin; FG: fasting glucose; FI: fasting insulin; 2hGlu: 2h glucose; Alleles E/O: effect allele/other allele; Freq. Effect Allele: frequency of effect allele; Effect (SE): effect size (standard error); *P*: p-value; N: number of samples in the analysis; Novel/previous glycemic trait association: Novel corresponds to a new association result; Locus name of previous association – name used for previously-reported locus. ^1^Significant in the European-only analysis in our study. ^2^Genome-wide significant association with T2D since date of analysis (Mahajan et al., 2018b). ^3^Association with T2D at *P*<1×10^−4^ since date of analysis (Mahajan et al., 2018b). ^4^T2D locus identified in Japanese (Hara et al.,2014) and Mexican (Williams et al., 2014) populations only. The date of our exomes analysis is May 2015. Related to Table S3.

| Trait | SNP | Gene | Protein Consequence | Alleles E/O | Freq. Effect Allele | Effect (SE) | P | N | Previous glycemic trait association (if any) | Locus name of previous association |
| --- | --- | --- | --- | --- | --- | --- | --- | --- | --- | --- |
| **FG** | **rs1886686** | **WDR78** | **p.G12A** | **G/C** | **0.739** | **0.014 (0.002)** | **2.24×10<sup>-11</sup>** | **123558** | **Novel** |  |
| HbA1c | rs267738 | CERS2 | p.E106A | G/T | 0.186 | -0.01 (0.002) | 6.96×10<sup>-10</sup> | 144043 | HbA1c | CERS2 |
| HbA1c | rs863362 | OR10X1 | p.W66X | T/C | 0.465 | 0.011 (0.001) | 6.76×10<sup>-15</sup> | 114945 | HbA1c | SPTA1 |
| HbA1c | rs857725 | SPTA1 | p.K1693Q | G/T | 0.262 | 0.022 (0.001) | 1.56×10<sup>-50</sup> | 143956 | HbA1c | SPTA1 |
| HbA1c | rs11887523 | MFSD2B | p.A60T | A/G | 0.007 | -0.072 (0.01) | 1.44×10<sup>-12</sup> | 122060 | HbA1c | ATAD2B |
| FG | rs1260326<sup>2</sup> | GCKR | p.L446P | C/T | 0.631 | 0.029 (0.002) | 6.36×10<sup>-48</sup> | 129588 | FG, FI, 2hGlu | GCKR |
| FI | rs1260326<sup>2</sup> | GCKR | p.L446P | C/T | 0.626 | 0.024 (0.002) | 5.55×10<sup>-32</sup> | 104076 | FG, FI, 2hGlu | GCKR |
| 2hGlu | rs1260326<sup>2</sup> | GCKR | p.L446P | C/T | 0.618 | -0.069 (0.009) | 4.48×10<sup>-15</sup> | 57813 | FG, FI, 2hGlu | GCKR |
| FG | rs35720761<sup>2</sup> | THADA | p.C845Y | T/C | 0.108 | -0.018 (0.003) | 4.35×10<sup>-9</sup> | 129622 | T2D | THADA |
| HbA1c | rs35720761<sup>2</sup> | THADA | p.C845Y | C/T | 0.113 | 0.014 (0.002) | 2.58×10<sup>-12</sup> | 144001 | T2D | THADA |
| FG | rs7578597 | THADA | p.T897A | C/T | 0.106 | -0.019 (0.003) | 1.99×10<sup>-8</sup> | 113162 | T2D | THADA |
| FI | rs7607980<sup>2</sup> | COBLL1 | p.N901D | C/T | 0.128 | -0.032 (0.003) | 1.30×10<sup>-24</sup> | 97817 | FI | COBLL1 |
| FG | rs2232323 | G6PC2 | p.Y207S | C/A | 0.006 | -0.129 (0.012) | 1.05×10<sup>-28</sup> | 123981 | FG, HbA1c | G6PC2 |
| HbA1c | rs2232323 | G6PC2 | p.Y207S | C/A | 0.007 | -0.053 (0.007) | 3.25×10<sup>-13</sup> | 144038 | FG, HbA1c | G6PC2 |
| FG | rs146779637 | G6PC2 | p.R283X | T/C | 0.002 | -0.138 (0.02) | 1.78×10<sup>-12</sup> | 127278 | FG, HbA1c | G6PC2 |
| HbA1c | rs146779637 | G6PC2 | p.R283X | T/C | 0.002 | -0.074 (0.012) | 4.58×10<sup>-10</sup> | 141728 | FG, HbA1c | G6PC2 |
| **FI** | **rs1983210** | **OBSL1** | **p.E1365D** | **G/C** | **0.729** | **0.016 (0.003)** | **8.48×10<sup>-10</sup>** | **79767** | **Novel** |  |
| **FI** | **rs3183099** | **OBSL1** | **splice region variant** | **A/G** | **0.226** | **-0.013 (0.002)** | **4.70×10<sup>-8</sup>** | **100713** | **Novel** |  |
| FI | rs1801282<sup>2</sup> | PPARG | p.P12A | G/C | 0.117 | -0.031 (0.003) | 3.50×10<sup>-23</sup> | 98631 | FI | PPARG |
| HbA1c | rs35726701 | RNF123 | p.K596E | G/A | 0.019 | 0.025 (0.005) | 4.19×10<sup>-8</sup> | 131203 | HbA1c | USP4 |
| FG | rs5400 | SLC2A2 | p.T110I | A/G | 0.161 | -0.022 (0.003) | 2.14×10<sup>-17</sup> | 129591 | FG, HbA1c | SLC2A2 |
| HbA1c | rs5400 | SLC2A2 | p.T110I | A/G | 0.153 | -0.013 (0.002) | 2.27×10<sup>-13</sup> | 144012 | FG, HbA1c | SLC2A2 |
| **HbA1c<sup>1</sup>** | **rs2237051** | **EGF** | **p.M708I** | **A/G** | **0.374** | **-0.007 (0.001)** | **2.11×10<sup>-7</sup>** | **121204** | **Novel** |  |
| HbA1c | rs7683365 | GYPB | p.T48M | A/G | 0.312 | 0.012 (0.002) | 1.61×10<sup>-8</sup> | 45191 | HbA1c | FREM3 |
| **FG** | **rs146886108<sup>2</sup>** | **ANKH** | **p.R187Q** | **T/C** | **0.004** | **-0.088 (0.014)** | **5.67×10<sup>-10</sup>** | **129647** | **Novel** |  |
| **HbA1c** | **rs31244** | **SV2C** | **p.D543N** | **A/G** | **0.083** | **0.012 (0.002)** | **6.05×10<sup>-8</sup>** | **144000** | **Novel** |  |
| FG | rs6235 | PCSK1 | p.S690T | G/C | 0.264 | -0.022 (0.002) | 9.22×10<sup>-24</sup> | 123560 | FG | PCSK1 |
| 2hGlu | rs2549782 | ERAP2 | p.K392N | T/G | 0.519 | -0.055 (0.009) | 6.81×10<sup>-10</sup> | 57836 | 2hGlu | ERAP2 |
| HbA1c | rs35742417<sup>3</sup> | RREB1 | p.S1499Y | A/C | 0.173 | -0.01 (0.002) | 3.76×10<sup>-9</sup> | 143967 | FG, T2D | RREB1 |
| FG | rs35742417<sup>3</sup> | RREB1 | p.S1499Y | A/C | 0.183 | -0.019 (0.002) | 1.27×10<sup>-16</sup> | 129577 | FG, T2D | RREB1 |
| HbA1c | rs1799945 | HFE | p.H63D | G/C | 0.129 | -0.023 (0.002) | 1.20×10<sup>-30</sup> | 128354 | HbA1c | HFE, HIST1H4A |
| HbA1c | rs1800562 | HFE | p.C279Y | A/G | 0.051 | -0.042 (0.003) | 3.30×10<sup>-47</sup> | 138093 | HbA1c | HFE, HIST1H4A |
| FG | rs10305492 | GLP1R | p.A316T | A/G | 0.014 | -0.08 (0.008) | 2.37×10<sup>-25</sup> | 129601 | FG | GLP1R |
| **HbA1c** | **rs35332062** | **MLXIPL** | **p.A358V** | **A/G** | **0.117** | **0.011 (0.002)** | **6.18×10<sup>-9</sup>** | **144042** | **Novel** |  |
| **HbA1c** | **rs3812316** | **MLXIPL** | **p.Q241H** | **G/C** | **0.112** | **0.012 (0.002)** | **2.15×10<sup>-8</sup>** | **108605** | **Novel** |  |
| **FG** | **rs194524<sup>3</sup>** | **STEAP2** | **p.R456Q** | **A/G** | **0.523** | **0.01 (0.002)** | **7.65×10<sup>-8</sup>** | **129629** | **Novel** |  |
| HbA1c | rs34664882 | ANK1 | p.A1503V | A/G | 0.026 | -0.049 (0.004) | 2.43×10<sup>-39</sup> | 144034 | HbA1c | ANK1 |
| FG | rs13266634<sup>2</sup> | SLC30A8 | p.R276W | T/C | 0.305 | -0.029 (0.002) | 1.63×10<sup>-46</sup> | 129614 | FG, HbA1c, T2D | SLC30A8 |
| HbA1c | rs13266634<sup>2</sup> | SLC30A8 | p.R276W | T/C | 0.300 | -0.015 (0.001) | 8.50×10<sup>-28</sup> | 143982 | FG, HbA1c, T2D | SLC30A8 |
| **HbA1c** | **rs11557154** | **DCAF12** | **p.R113Q** | **T/C** | **0.138** | **-0.009 (0.002)** | **1.70×10<sup>-7</sup>** | **144045** | **Novel** |  |
| FG | rs17853166 | IKBKAP | p.S251G | C/T | 0.026 | -0.037 (0.006) | 4.82×10<sup>-11</sup> | 129640 | FG | IKBKAP |
| HbA1c | rs60980157<sup>2</sup> | GPSM1 | p.S391L | T/C | 0.246 | -0.013 (0.002) | 6.71×10<sup>-17</sup> | 118824 | FG, T2D | GPSM1 |
| FG | rs60980157<sup>2</sup> | *GPSM1* | p.S391L | T/C | 0.254 | -0.014 (0.002) | 2.35×10<sup>-9</sup> | 110915 | FG, T2D | *GPSM1* |
| HbA1c | rs906220 | *HK1* | p.H7R | G/A | 0.916 | 0.025 (0.003) | 2.16×10<sup>-21</sup> | 94970 | HbA1c | *HK1* |
| **FG** | **rs701865** | ***PDE6C*** | **p.S270T** | **A/T** | **0.366** | **-0.01 (0.002)** | **1.14×10<sup>-7</sup>** | **118580** | **Novel** |  |
| **HbA1c** | **rs61732434** | ***OR51V1*** | **p.S161N** | **T/C** | **0.008** | **-0.052 (0.009)** | **1.75×10<sup>-8</sup>** | **127507** | **Novel** |  |
| **HbA1c** | **rs415895** | ***SWAP70*** | **p.Q447E** | **G/C** | **0.641** | **-0.013 (0.001)** | **1.15×10<sup>-21</sup>** | **138028** | **Novel** |  |
| **HbA1c** | **rs117706710** | ***AMPD3*** | **p.V311L** | **T/G** | **0.009** | **0.037 (0.006)** | **2.32×10<sup>-10</sup>** | **144048** | **Novel** |  |
| FG | rs2167079 | *ACP2* | p.R29Q | T/C | 0.340 | 0.016 (0.002) | 7.99×10<sup>-15</sup> | 129580 | FG | *MADD* |
| HbA1c | rs35233100 | *MADD* | p.R766X | T/C | 0.055 | -0.015 (0.003) | 1.13×10<sup>-8</sup> | 144034 | FG | *MADD* |
| FG | rs35233100 | *MADD* | p.R766X | T/C | 0.054 | -0.029 (0.004) | 1.46×10<sup>-12</sup> | 126231 | FG | *MADD* |
| FG | rs56200889<sup>2</sup> | *ARAP1* | p.Q802E | C/G | 0.270 | -0.016 (0.002) | 1.79×10<sup>-14</sup> | 122674 | FG | *ARAP1* |
| **HbA1c** | **rs643788** | ***DPAGT1*** | **p.I393V** | **C/T** | **0.425** | **-0.006 (0.001)** | **1.77×10<sup>-7</sup>** | **144009** | **Novel** |  |
| **FI<sup>1</sup>** | **rs145878042** | ***RAPGEF3*** | **p.L300P** | **G/A** | **0.011** | **-0.054 (0.01)** | **1.15×10<sup>-7</sup>** | **91485** | **Novel** |  |
| HbA1c | rs2732481 | *ZNF641* | p.Q363P | G/T | 0.315 | -0.009 (0.001) | 2.07×10<sup>-11</sup> | 142280 | HbA1c | *SENP1* |
| HbA1c | rs3184504 | *SH2B3* | p.W262R | C/T | 0.567 | 0.007 (0.001) | 5.98×10<sup>-8</sup> | 138551 | HbA1c | *ATXN2* |
| 2hGlu | rs1169288<sup>2</sup> | *HNF1A* | p.I75L | C/A | 0.345 | 0.06 (0.011) | 7.90×10<sup>-9</sup> | 44278 | T2D | *HNF1A* |
| HbA1c | COSM147717 | *ATP11A* | p.M317V | G/A | 0.748 | 0.009 (0.001) | 3.77×10<sup>-12</sup> | 144022 | HbA1c | *ATP11A,TUB*<br>*GCP3* |
| **HbA1c** | **rs229587** | ***SPTB*** | **p.S439N** | **T/C** | **0.357** | **0.007 (0.001)** | **2.60×10<sup>-8</sup>** | **134780** | **Novel** |  |
| HbA1c | rs35097172 | *SLC25A47* | splice region<br>variant, 5'<br>UTR variant | T/C | 0.216 | -0.008 (0.002) | 5.67×10<sup>-8</sup> | 144028 | FG | *SLC25A47* |
| 2hGlu | rs3784634 | *VPS13C* | p.R974K | T/C | 0.540 | -0.069 (0.011) | 6.40×10<sup>-10</sup> | 37217 | 2hGlu | *VPS13C/*<br>*C2CD4A/*<br>*C2CD4B* |
| **HbA1c<sup>1</sup>** | **rs3747481** | ***PRR14*** | **p.P359L** | **T/C** | **0.261** | **0.009 (0.002)** | **3.30×10<sup>-8</sup>** | **103338** | **Novel** |  |
| HbA1c | rs201226914 | *PIEZO1* | p.L939M | T/G | 0.002 | -0.159 (0.015) | 4.42×10<sup>-26</sup> | 144024 | HbA1c | *CDT1,CYBA* |
| 2hGlu | rs72839768<sup>4</sup> | *DVL2* | p.T529I | A/G | 0.020 | 0.197 (0.03) | 4.10×10<sup>-11</sup> | 57866 | T2D | *SLC16A13* |
| HbA1c | rs2748427 | *TMC6* | p.W125R | G/A | 0.233 | 0.027 (0.002) | 8.56×10<sup>-70</sup> | 132326 | HbA1c | *TMC6* |
| HbA1c | rs7225887 | *B3GNTL1* | p.A163T | T/C | 0.211 | -0.015 (0.002) | 5.73×10<sup>-22</sup> | 125749 | HbA1c | *FN3KRP,*<br>*FN3K* |
| **HbA1c** | **rs35413309** | ***RGS9BP*** | **p.A223V** | **T/C** | **0.030** | **-0.02 (0.004)** | **1.42×10<sup>-8</sup>** | **141598** | **Novel** |  |
| 2hGlu | rs1800437<sup>2</sup> | *GIPR* | p.E318Q | C/G | 0.217 | 0.103 (0.011) | 2.59×10<sup>-23</sup> | 56252 | 2hGlu | *GIPR* |
| FG | rs17265513<sup>3</sup> | *ZHX3* | p.N310S | C/T | 0.188 | 0.016 (0.002) | 2.59×10<sup>-10</sup> | 126253 | FG | *ZHX3* |
| HbA1c | rs855791 | *TMPRSS6* | V727A | G/A | 0.577 | -0.019 (0.001) | 9.46×10<sup>-51</sup> | 143907 | HbA1c | *TMPRSS6* |
| **FG** | **rs15943** | ***MAP3K15*** | **p.Q1083E** | **C/G** | **0.005** | **-0.084 (0.014)** | **2.83×10<sup>-9</sup>** | **67004** | **Novel** |  |
| **FG** | **rs56381411** | ***MAP3K15*** | **p.G670S** | **T/C** | **0.005** | **-0.085 (0.013)** | **1.51×10<sup>-11</sup>** | **62319** | **Novel** |  |
| HbA1c | rs2229241 | *RENBP* | splice<br>acceptor<br>variant | C/T | 0.012 | -0.123 (0.007) | 1.14×10<sup>-62</sup> | 95622 | HbA1c | *G6PD* |
| HbA1c | rs1050828 | *G6PD* | p.V68M | T/C | 0.007 | -0.334 (0.008) | 7.41×10<sup>-322</sup> | 112209 | HbA1c | *G6PD* |

To increase power to detect rare variant associations, we additionally performed gene-burden and sequence kernel association (SKAT) tests for gene-level analyses (**Methods**). We identified six genes with significant evidence of association (*P*<2.5 × 10^−6^), of which two – *G6PC* (for FG and FI) and *TF* (for HbA1c) – represented novel associations (**Tables 2 and S4**).

**Table 2.** Gene-based results from broad (NSbroad mask) and strict (NSstrict mask) analyses. Genes in bold are newly discovered from this effort. N var: total number of variants in that gene-based analysis; *P*_burden_: p-value from burden test which assumes all variants have the same direction of effect; *P*_SKAT_: p-value from SKAT test which allows for different directions of effect between variants. The lowest p-value is highlighted in bold. Related to Table S4.

| Trait | Gene | NSbroad mask |  |  | NSstrict mask |  |  |
| --- | --- | --- | --- | --- | --- | --- | --- |
| | | N var | $P_{burden}$ | $P_{SKAT}$ | N var | $P_{burden}$ | $P_{SKAT}$ |
| FG | **G6PC** | 9 | **1.41x10<sup>-6</sup>** | 1.32x10<sup>-5</sup> | 3 | 1.41x10<sup>-3</sup> | 7.43x10<sup>-4</sup> |
| FI | **G6PC** | 8 | **1.62x10<sup>-6</sup>** | 8.58x10<sup>-6</sup> | 3 | 1.85x10<sup>-3</sup> | 7.80x10<sup>-3</sup> |
| HbA1c | **TF** | 10 | **2.15x10<sup>-6</sup>** | 5.98x10<sup>-3</sup> | 3 | 5.48x10<sup>-2</sup> | 5.48x10<sup>-2</sup> |
| FG | MAP3K15 | 18 | **1.86x10<sup>-25</sup>** | 1.07x10<sup>-18</sup> | 7 | 1.34x10<sup>-14</sup> | 4.01x10<sup>-11</sup> |
| HbA1c | MAP3K15 | 18 | **1.27x10<sup>-7</sup>** | 1.53x10<sup>-04</sup> | 7 | 2.65x10<sup>-4</sup> | 9.46x10<sup>-3</sup> |
| FG | G6PC2 | 18 | 4.09x10<sup>-67</sup> | 5.38x10<sup>-58</sup> | 7 | **7.8x10<sup>-69</sup>** | 3.83x10<sup>-56</sup> |
| HbA1c | G6PC2 | 18 | 6.18x10<sup>-30</sup> | 4.65x10<sup>-27</sup> | 7 | **1.04x10<sup>-31</sup>** | 1.92x10<sup>-26</sup> |
| FG | SLC30A8 | 13 | 5.69x10<sup>-4</sup> | **6.42x10<sup>-11</sup>** | 7 | 6.55x10<sup>-11</sup> | 3.74x10<sup>-10</sup> |
| HbA1c | SLC30A8 | 12 | 7.20x10<sup>-8</sup> | 2.18x10<sup>-5</sup> | 6 | **5.66x10<sup>-8</sup>** | 3.22x10<sup>-6</sup> |
| FG | VPS13C | 52 | 9.66x10<sup>-6</sup> | **3.73x10<sup>-7</sup>** | 26 | 1.27x10<sup>-5</sup> | 1.44x10<sup>-5</sup> |

### Identification of effector transcripts

To establish whether the associated coding variants (both novel and those at established loci) were likely to be causal, and/or likely to pinpoint an effector transcript, we first integrated these results with published data with higher density GWAS coverage (Manning et al., 2012; Wheeler et al., 2017a). This is important because coding variants can sometimes erroneously point to the wrong effector transcript, as they can “piggy-back” on non-coding alleles that drive the association, and by virtue of having a predicted effect on protein sequence they may falsely implicate the gene in which they reside as the causal one. For example, the coding variant rs56200889 (p.Q802E) at *ARAP1* is strongly associated with FG (β=-0.016, *P*=1.8 × 10^−14^, **Table 1**), and when considered in isolation might have suggested *ARAP1* as the effector transcript. However, T2D fine-mapping efforts showed this association to be secondary to a much stronger non-coding signal (Mahajan et al., 2018b), and recent data integrating human islet and mouse knockout information has established neighbouring gene *STARD10* as the most likely gene mediating the GWAS signal at this locus (Carrat et al., 2017). Therefore, we conditioned the coding variants identified here on existing non-coding GWAS index variants at established loci from two previously published GWAS datasets (Manning et al., 2012; Wheeler et al., 2017a), and also performed the reciprocal analysis (**Table S3, Methods**). At novel loci, we also assessed whether the coding variant identified here was being driven by association of a sub-threshold (i.e. non genome-wide significant in smaller sample size) non-coding variant based on published GWAS results with higher density coverage (Manning et al., 2012; Wheeler et al., 2017a) (**Methods**). As reciprocal conditional analysis was not always possible, or was not informative, we also used additional published data, including fine-mapping results from comparable T2D efforts (Mahajan et al., 2018b), results for associations with blood cell traits (Astle et al., 2016; Soranzo et al., 2010) (**Table S5**), as well as a body of literature establishing the role of certain genes (mapping within our loci) in glucose metabolism, or red blood cell biology (for HbA1c) to inform effector transcript classification. We further considered significant gene-based associations driven by multiple coding variants within a single gene as strong evidence for the determination of effector transcripts (**Methods**).

Combining the above approaches, we curated the 74 coding variant associations (in 58 genes) displayed in Table 1, and where possible identified and classified effector transcripts into “gold”, “silver” and “bronze” categories, depending on the strength of evidence (**Table S6, Methods**). Loci with strong evidence from reciprocal conditional analysis or from published data that supported the relevance of the identified effector transcript to the glycemic trait were labelled “gold” (e.g. *GLP1R, SLC30A8, G6PD, PPARG, ANK1*); those where an effector transcript could not be defined by conditional analysis (either because it was inconclusive or due to lack of data) but where there was strong biological plausibility for a given gene at the locus were labelled “silver” (e.g. *MADD, MLXIPL, FN3K/FN3KRP, HK1, VPS13C*); those where we had some evidence but that was not as strong as “silver” were labelled “bronze” (e.g. *DCAF12, OBSL1, STEAP2, RAPGEF3*); the remaining were left with an undetermined effector transcript (**Figure 1, Table S6**). Effector transcript classification into the three categories was undertaken independently by four of the authors and the consensus was used as the final classification for effector transcripts. From 74 single variant and six gene-based signals, we identified 51 unique effector transcripts (24 gold, 11 silver, 16 bronze), with many of them shared across traits (**Figure 1**). One case in point pertains to *VPS13C*, which harboured a missense variant (p.R974K) associated with 2hGlu (labelled “bronze”) at exome-wide significance (β=-0.069, *P*=6.4 × 10^−10^; **Table 1**), and also exhibited a significant gene-based association with FG (labelled “silver”; *P*_SKAT_=3.7 × 10^−7^; **Table S4**). *VPS13C* belongs to the previously-established *VPS13C*/*C2CD4A*/*C2CD4B* glycemic trait and T2D risk locus, and recent follow-up studies have with varying levels of evidence suggested *C2CD4A*, encoding a calcium-dependent nuclear protein, as the causal gene for T2D through its potential role in the pancreatic islets (Kycia et al., 2018; Mehta et al., 2016; O’Hare et al., 2016). In our data, it is however not possible to rule out *VPS13C* as a potential effector transcript at this locus, warranting further functional studies for *VPS13C*, which encodes a protein reported to be necessary for proper mitochondrial function (Lesage et al., 2016).

**Figure 1.**
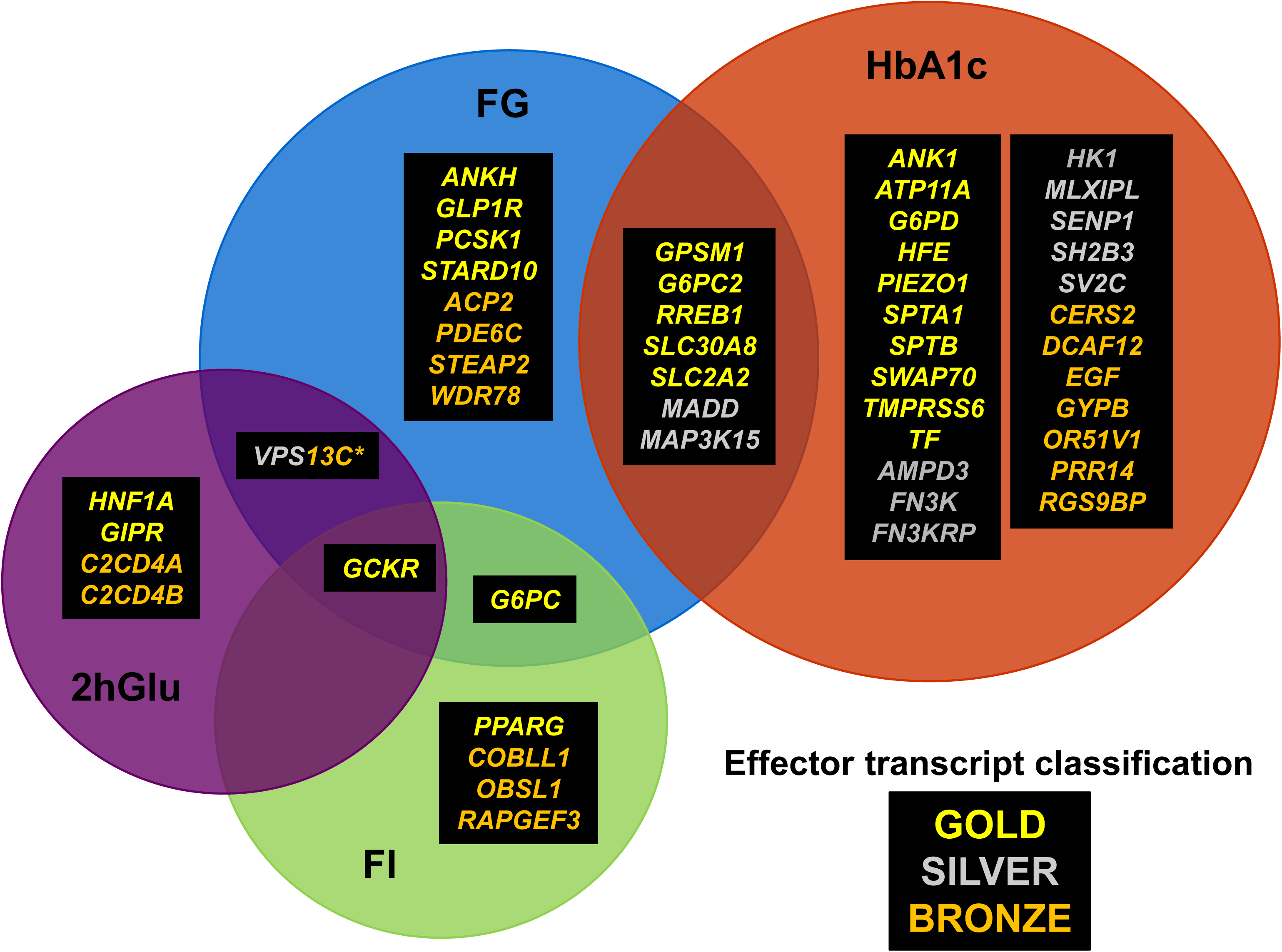
Effector transcript classification into “gold”, “silver” and “bronze” categories based on strength of genetic and biological evidence. A total of 51 effector transcripts from 74 single variant and six gene-based signals were identified, with many of them shared across traits. The classification was undertaken independently by four of the authors and the consensus was used as the final classification for effector transcripts (see **Methods**). *Asterisk indicates “silver” for FG, “bronze” for 2hGlu.

### Pathway analyses identifies relevant gene sets regulating glycemia

To identify pathways enriched for glycemic trait associations, and to subsequently determine the extent to which associations within the same trait implicate the same or similar pathways (as indicated by the functional connectivity of the network), we used GeneMANIA network analysis (Franz et al., 2018). GeneMANIA takes a query list of genes and finds functionally-similar genes based on large, publicly-available biological datasets. We analysed all loci harbouring non-synonymous variants that reached *P*<1 × 10^−5^ for any of the four glycemic traits in our study (totaling 121 associations). A high degree of connectivity was observed within the HbA1c network, with enrichment of processes related to blood cell biology such as porphyrin metabolism, erythrocyte homeostasis and iron transport (**Figures 2 and S1, Table S7**). In comparison, the network generated from FG-associated genes captured several processes known to contribute to glucose regulation and islet function, including insulin secretion, zinc transport and fatty acid metabolism (**Figure 2, Table S7**). The FG network further revealed linking nodes (that are not among the association signals) with known links to glucose homeostasis and diabetes, such as *GCK* (encoding the beta cell glucose sensor glucokinase), *GCG* (encoding the peptide hormone glucagon secreted by the alpha cells of the pancreas) and *GIP* (encoding the incretin hormone gastric inhibitory polypeptide). One gene within the FG cluster for lipid-related pathways is *CERS2*, which encodes ceramide synthase 2, an enzyme known to be associated with the sphingolipid biosynthetic process (**Figure 2, Table S7**). Although *CERS2* is only nominally associated with FG and is significantly associated with HbA1c, it does not cluster together with any HbA1c-enriched pathway, suggesting that *CERS2* is regulating FG and HbA1c indirectly through its role in lipid metabolism. Given that there were fewer genes associated with FI and 2hGlu, we were less powered to draw meaningful insights from the enriched pathways in those traits (**Figure S1, Table S7**).

**Figure 2.**
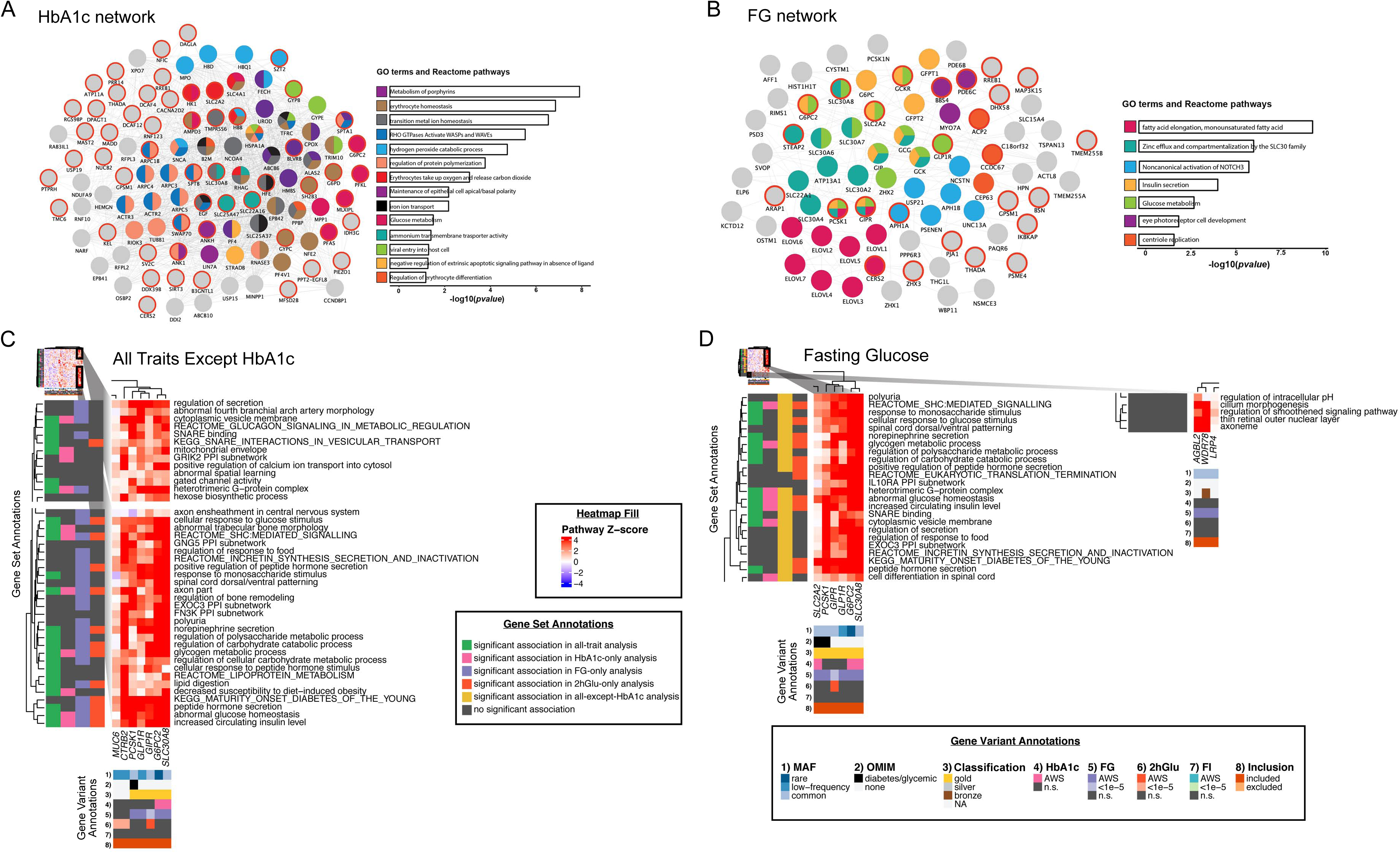
Network and pathway analyses identify relevant gene sets regulating glycemia using two different methods for variant associations with P<1 × 10-5. (A-B) The networks represent composite networks for (A) HbA1c and (B) FG, from the GeneMANIA analysis using genes with variant associations at P<1 × 10-5 for each trait as input. Nodes outlined in red correspond to genes from the input list. Other nodes correspond to related genes based on 50 default databases. Based on the network, GO terms and Reactome pathways that were significantly enriched are depicted. To summarize these results, the most significant term of all calculated terms within the same group is represented. Barplots with the Bonferroni-adjusted -log10(p-values) of the most significant terms within each group are are shown. Each group was assigned a specific color; if a gene is present in more than one term, it is displayed in more than one color. (C-D) Heatmaps showing EC-DEPICT results from analysis of (C) all traits except HbA1c and (D) FG. The columns represent the input genes for the analysis. In (C), these are genes with variant associations of P<1 × 10-5 for FG, FI, and/or 2hGlu, and in (D) these are genes with variant associations of P<1 × 10-5 for FG. Rows in the heatmap represent significant meta-gene sets (FDR <0.05). The color of each square indicates DEPICT’s z-score for membership of that gene in that gene set, where dark red means “very likely a member” and dark blue means “very unlikely a member.” The gene set annotations indicate whether that meta-gene set was significant at FDR <0.05 or not significant (n.s.) for each of the other EC-DEPICT analyses. For heatmap intensity and EC-DEPICT P-values, the meta-gene set values are taken from the most significantly enriched member gene set. The gene variant annotations are as follows: (1) the European minor allele frequency (MAF) of the input variant, where rare is MAF <1%, low-frequency is MAF 1-5%, and common is MAF >5%, (2) whether the gene has an Online Mendelian Inheritance in Man (OMIM) annotation as causal for a diabetes/glycemic-relevant syndrome or blood disorder, (3) the effector transcript classification for that variant: gold, silver, bronze, or NA (note that only array-wide significant variants were classified, so suggestively-significant variants are by default classified as “NA”), (4-7) whether each variant was significant (P<2 × 10-7), suggestively significant (P<1 × 10-5), or not significant in Europeans for each of the four traits, and (8) whether each variant was included in the analysis or excluded by filters (see Methods). AWS: array-wide significant. Related to Figures S1 to S3.

We also performed gene set enrichment analysis (GSEA) using EC-DEPICT (Marouli et al., 2017; Turcot et al., 2018) (**Methods**). The primary innovation of EC-DEPICT is the use of 14,462 gene sets extended based on large-scale co-expression data (Fehrmann et al., 2015; Pers et al., 2015). These gene sets take the form of z-scores, where higher z-scores indicate a stronger prediction that a given gene is a member of a gene set. To reduce some of the redundancy in the gene sets (many of which are strongly correlated with one another), we clustered them into 1,396 “meta-gene sets” using affinity propagation clustering (Frey and Dueck, 2007). These meta-gene sets are used to simplify visualizations and aid interpretation of results. Here, we combined and analyzed all variants that reached *P*<1 × 10^−5^ for any of the four glycemic traits (**Methods**). We found 234 significant gene sets in 86 meta-gene sets with false discovery rate (FDR) of <0.05 (**Table S8**). As expected, we observed a strong enrichment of insulin- and glucose-related gene sets, as well as exocytosis biology (in keeping with insulin vesicle release). In agreement with the GeneMANIA network analyses, we also noted a strong enrichment for blood-related pathways, which was primarily driven by HbA1c-associated variants. This was likely because HbA1c levels are influenced not only by glycation but also by blood cell turnover rate (Cohen et al., 2008; Wheeler et al., 2017a). To disentangle blood cell turnover from effects due to glycation, we repeated the analysis excluding variants that were significantly associated with HbA1c only and found 128 significant gene sets in 53 meta-gene sets (FDR <0.05) (**Table S8**). We also analyzed each of the four traits separately (**Table S8, Methods**).

To identify additional candidate genes, we then performed heat map visualization with unsupervised clustering of the membership predictions (z-scores) of trait-associated genes for each significant gene set (**Figures 2, S2 and S3**). This strategy has previously been effective for gene prioritization for downstream analyses (Marouli et al., 2017; Turcot et al., 2018), as it becomes visually apparent which genes are the strongest drivers of the significant gene sets and thus are natural targets for follow-up. This can be particularly helpful for prioritizing genes that are not well-characterized, as it leverages DEPICT’s prediction of gene function. For the analysis of all traits except HbA1c, one cluster showed particularly strong predicted membership for highly relevant gene sets, including “abnormal glucose homeostasis”, “peptide hormone secretion”, “Maturity Onset Diabetes of the Young”, and multiple pathways involved in the regulation of glycogen, incretin, and carbohydrate metabolism (**Figure 2C**). Strikingly, this cluster of six genes (*PCSK1, GLP1R, GIPR, G6PC2, SLC30A8* and *CTRB2*) contained five of the genes that had independently been assigned to “gold” status during effector transcript identification (**Table S6**). Therefore, the sixth gene, *CTRB2*, represents a novel gene for prioritization, since it showed strong similarity to other genes for which there was already substantial biological evidence. *CTRB2* encodes chymotrypsinogen B2, a digestive enzyme that is expressed in the exocrine pancreas, and subsequently secreted into the gut. The gene contains a borderline significant variant for 2hGlu (rs147238447; p.L6V; *P*=1.9 × 10^−6^). Another variant at this locus, rs7202877 (6.2kb downstream of *CTRB2*, r^2^=0.0006, D’=1 with rs147238447 in European populations), has previously been shown to be an eQTL for *CTRB1* and *CTRB2*, with the minor G allele (MAF=11%) associated with increased expression (t Hart et al., 2013). In the same study, the rs7202877-G allele was associated with increased glucagon-like peptide 1 (GLP-1)-stimulated insulin secretion (*P*=8.8 × 10^−7^, N=196). In our data, rs7202877-G was nominally associated with lower 2hGlu (*P*=6.3 × 10^−3^) and lower FG (*P*=2.8 × 10^−3^) levels. Multiple distinct signals in this region (previously referred to as the *BCAR1* locus) have also been associated with T2D risk, including rs7202877 (where the G allele is protective), rs72802342 (r^2^=0.65 with rs7202877 in European populations) and rs3115960, although the coding variant rs147238447 described here is not (Mahajan et al., 2018a; Mahajan et al., 2018b; Morris et al., 2012; Zhao et al., 2017). This can potentially be explained by limited power to identify a significant association given the low MAF (∼0.5%) of the coding variant. In contrast to its effect on T2D, the rs7202877-G allele has been associated with increased risk of type 1 diabetes (OR=1.28, *P*=3.1 × 10^−15^, N=21,293) (Barrett et al., 2009). Other variants at this locus are associated with risk of chronic pancreatitis (rs8055167, r^2^=0.0021 with rs147238447 and r^2^=0.12 with rs7202877 in European populations, in LD with an inversion that changes the expression ratio of *CTRB1* and *CTRB2* isoforms) (Rosendahl et al., 2017) and pancreatic cancer (rs7190458, r^2^=0.0002 with rs147238447 and r^2^=0.31 with rs7202877 in European populations) (Wolpin et al., 2014). The prioritization of *CTRB2* is intriguing as it supports an emerging hypothesis that the exocrine pancreas contributes to complex mechanisms influencing 2hGlu levels and diabetes risk (Esteghamat et al., 2019; Hart et al., 2018; Woodmansey et al., 2017). Given the earlier associations with GLP-1 stimulated insulin secretion, we investigated whether this effect could be mediated by incretin levels. However, we found no associations at rs147238447 for GLP-1 levels in the largest available dataset (fasting GLP-1, N=4170: MAF=0.00457, *P*=0.495; 2h GLP-1, N=3839: MAF=0.00464, *P*=0.076) (Almgren et al., 2017), though this might again be explained by limited power. Although additional validation of the rare coding variant rs147238447 (p.L6V) as a potential causal variant is required given the absence of clear associations with T2D risk and other glycemic traits, the results discussed above suggest a role of *CTRB2* in glycemic regulation.

We also noted a small but distinct cluster in the FG-only analysis indicating the role of the cilium/axoneme, pointing to novel biology relating to sensing and signaling in response to the extracellular environment (**Figure 2D**). Two genes were the main drivers of this association: *WDR78* and *AGBL2*. These represent potentially interesting candidates for follow-up, although we note that the *AGBL2* signal may be driven through effects of the nearby *MADD* gene, which harbors a FG-associated coding variant in our study and is labelled “silver” in our effector transcript classification (**Tables 1 and S6**). Overall, our network and pathway analyses highlighted several trait-associated genes that do not reach exome-wide significance in conventional single variant or gene-based tests, but show evidence of contribution to glycemic regulation.

### Tissue enrichment analysis reveals shared roles of key tissues in the regulation of glycemic traits

In addition to identifying key metabolic pathways involved in glucose regulation, we sought to establish the relative importance of particular tissues in the regulation of the different glycemic phenotypes. This time, we assessed the tissues that are most highly enriched for the expression of the 51 effector transcripts we have curated at the associated loci identified in this study, to highlight specific tissues that contribute critically to the regulation of each glycemic trait. Using publicly-available tissue expression data from GTEx (Battle et al., 2017) and human islets (van de Bunt et al., 2015), we noted clear differences in tissue enrichment patterns as well as tissues shared between traits (**Figure 3**). Comparisons between analyses of FG- and FI-associated effector transcripts underscored the relative roles of the liver in both traits (*P*<0.05), whereas pancreatic islets were enriched in associations for FG (*P*=9.99 × 10^−5^) but not FI (*P*=0.75). In contrast, adipose (*P*=0.01) and kidney tissues (*P*=0.01) were enriched in FI but not FG (*P*>0.05). These results not only highlight the established role of pancreatic islets in influencing FG levels, but also the under-appreciated role of insulin clearance in the kidney and likely the liver, in addition to insulin action in liver and adipose tissue, in influencing FI levels (Goodarzi et al., 2011). Consistent with the EC-DEPICT GSEA, there was also support for the role of the exocrine pancreas (which typically represents >95% of whole pancreas tissue) in addition to the endocrine pancreas (islets) in FG (*P*=9.99 × 10^−5^) and 2hGlu (*P*=2.99 × 10^−4^) associations. We also observed enrichment for genes expressed in stomach for 2hGlu (*P*=1.99 × 10^−4^) but not for FG (*P*=0.16). HbA1c analysis revealed enrichment in “metabolic” tissues reflecting insulin secretion (islets, *P*=1.59 × 10^−2^ and pancreas, *P*=0.01), insulin action (muscle, *P*=1.50 × 10^−2^), insulin clearance (liver, *P*=0.03), as well as strong enrichment for whole blood (*P*=3.99× 10^−3^). These indicate key factors relating to hemoglobin glycation and blood cell function in influencing overall HbA1c levels (**Figure 3**).

**Figure 3.**
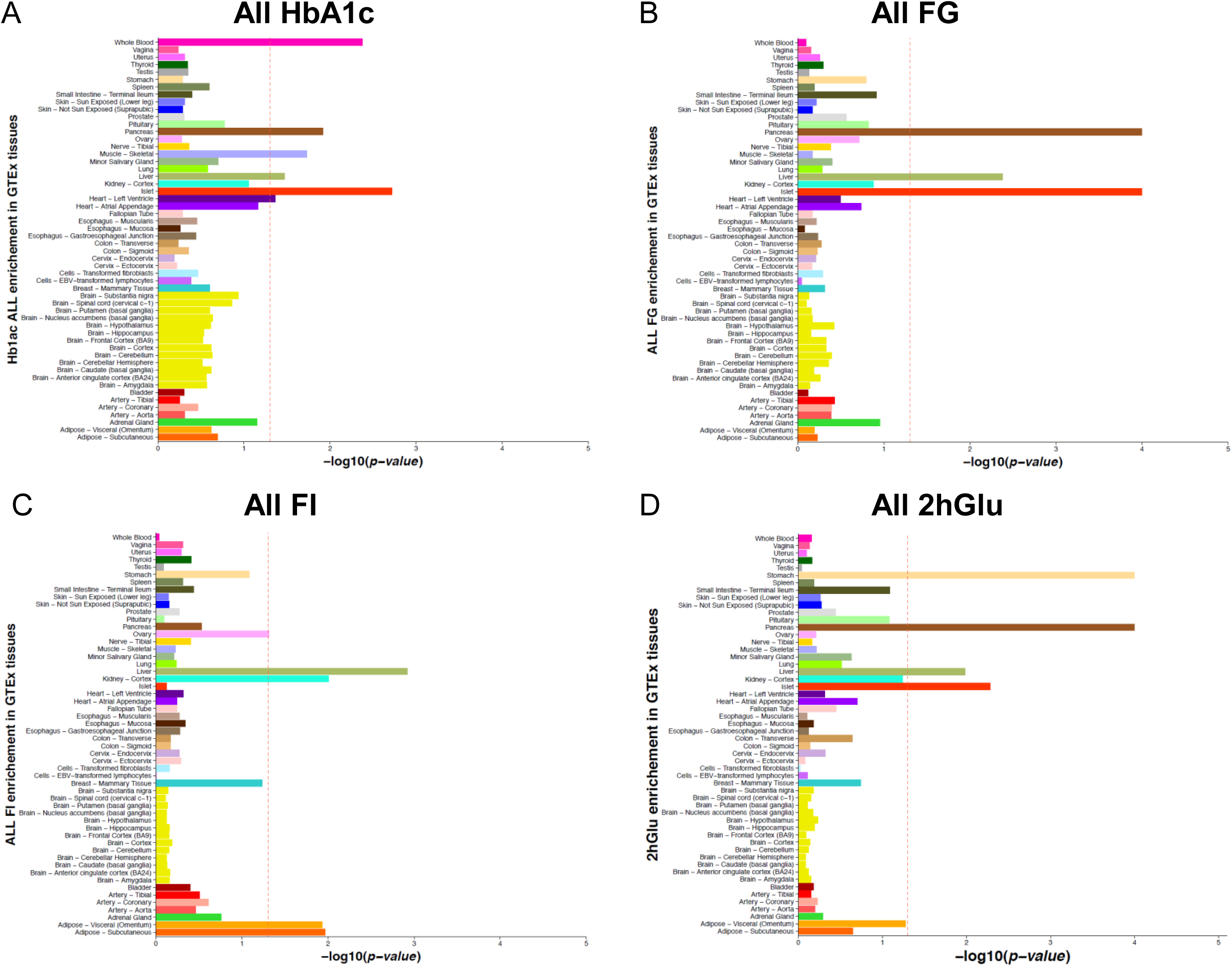
Tissue enrichment analysis reveals the key tissues involved in the regulation of glycemic traits. The figures display expression enrichment of genes from all of the golden, silver, and bronze gene set lists for (A) HbA1c, (B) FG, (C) FI and (D) 2hGlu in GTEx tissue samples plus islet data. Enrichment *P*-values were assessed empirically for each tissue using a permutation procedure (10,000 iterations), and the red vertical line shows the significance threshold (empirical *P*<0.05).

Our results from the pathway and tissue enrichment analyses demonstrate the role of specific tissues with known functions in blood glucose regulation in particular glycemic traits. These observations add further support to emerging reports of an underappreciated role for the exocrine pancreas in FG and 2hGlu regulation, the stomach-incretin axis in 2hGlu, and the importance of insulin clearance through the kidney and liver in FI.

### Novel glycemic trait associations in liver-enriched *G6PC* are driven by functional coding variants

To delve deeper into tissue-specific gene effects, we focused on two homologues, *G6PC* and *G6PC2*, with constrasting tissue expression profiles where we identified gene-based association signals for FG/FI and FG/HbA1c, respectively (**Tables 2 and S4**). Both genes encode gluconeogenic enzymes that catalyze the same biochemical pathway but are known to have distinct tissue expression profiles. *G6PC2* is largely expressed in pancreatic islets whereas *G6PC* is highly expressed in the liver, kidney, and small intestine (Foster et al., 1997; Mithieux, 1997). Our gene-based analyses highlighted *G6PC* through novel associations with FG and FI, driven primarily by rare missense variants p.A204S (rs201961848) and p.R83C (rs1801175), and protein-truncating variant (PTV) p.Q347X (rs80356487), none of which achieved exome-wide significance at single-variant level (**Table S4**). Homozygous inactivating alleles in *G6PC*, which include both p.R83C and p.Q347X, are known to give rise to glycogen storage disease type Ia (GSD1a), a rare autosomal recessive metabolic disorder (Chou and Mansfield, 2008; Lei et al., 1995), but this is the first time that rare coding variants in *G6PC* have been shown to influence FG and FI levels in normoglycemic individuals.

Given the well-known role of *G6PC* in hepatic glucose homeostasis, we were interested in elucidating the molecular impact of rare heterozygous *G6PC* coding variants highlighted in our exome-array analysis, in particular novel variant p.A204S, one of the statistical drivers of the gene-based *G6PC* signal (**Table S4**). In transient protein overexpression assays, p.R83C and p.A204S resulted in significantly reduced protein levels compared to wild type (WT) G6PC in both Huh7 (human hepatoma) and HEK293 (human embryonic kidney) cell lines (**Figure 4A-D**). The PTV p.Q347X, which in our *in vitro* system generated a smaller molecular weight protein, exhibited markedly lower protein expression levels in Huh7 cells but not HEK293 cells. However, in both cell types, the cellular localization pattern of p.Q347X appears to be largely diffuse and did not co-localize with the Golgi apparatus, which is important for post-translational modification of G6PC protein (**Figures 4E and S4A**). Further functional characterization of glucose-6-phosphatase (G6Pase) activity revealed that both p.R83C and p.Q347X variants lead to proteins lacking any detectable phosphatase activity (**Figure S4B-C**), consistent with previous observations of several GSD1a-causing coding variants (Shieh et al., 2002). As we observed that the p.R83C variant resulted in complete loss of glycosylation, we determined if glycosylation is essential for G6Pase activity by treating cells with tunicamycin to inhibit N-linked glycosylation. The ability of unglycosylated G6PC to catalyze glucose-6-phosphate (G6P) was found to be downregulated by up to 14%, although this difference was not statistically significant (**Figure 4F**). We therefore concluded that whilst glycosylation contributes to overall functional activity, it may not be a requisite for G6P hydrolysis. Finally, we were unable to accurately assess p.A204S-G6PC phosphatase activity as the level of expression in the microsomes was reduced by 41% relative to WT, supporting the hypothesis that p.A204S-G6PC exhibits partial loss-of-function (LOF) most likely due to loss of protein expression.

**Figure 4.**
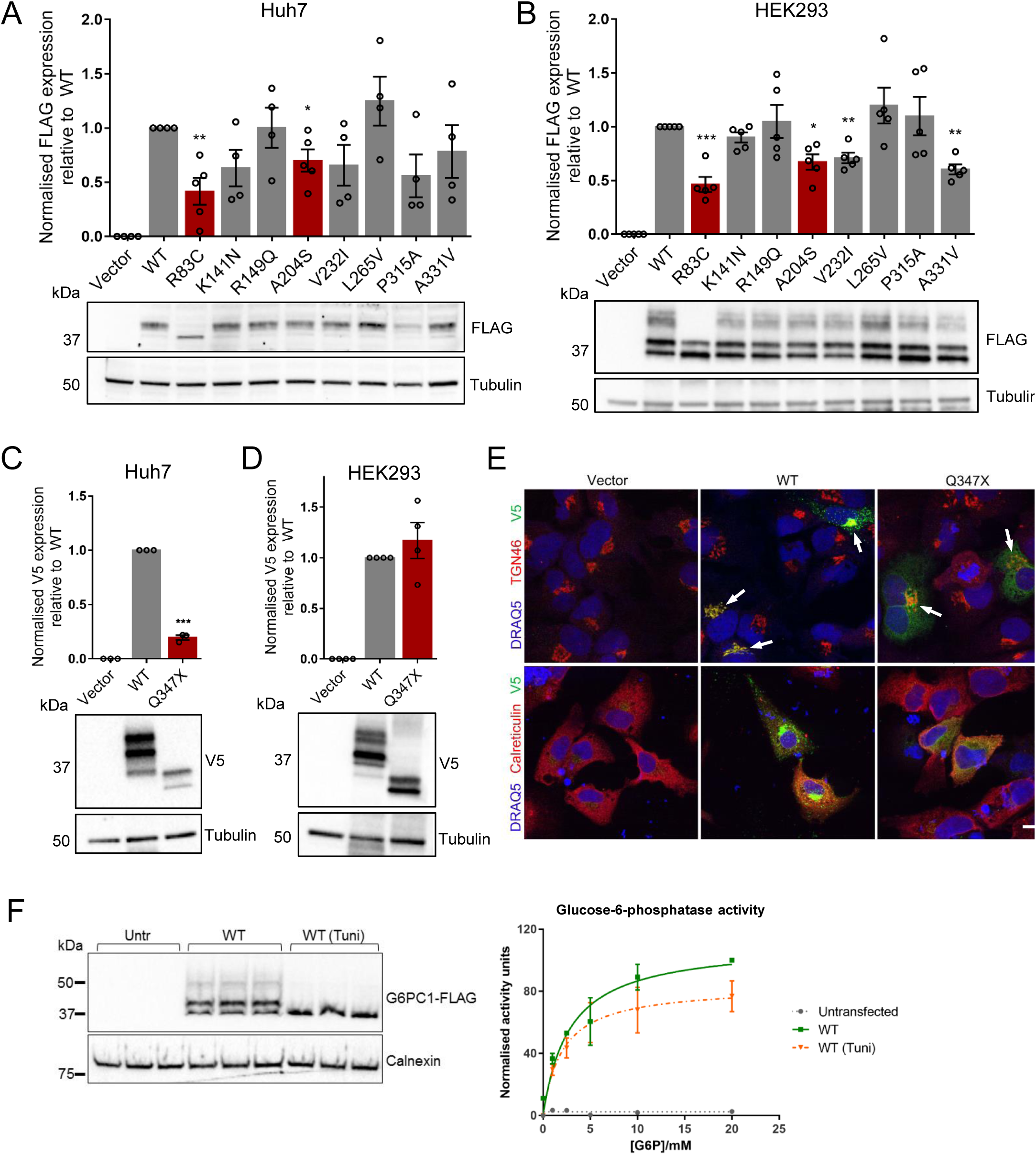
Functional characterisation of G6PC variant proteins. Related to Figure S4. (A) Protein expression levels of missense G6PC variants were determined in Huh7 cells (n=4-5) and (B) HEK293 cells (n=5) by western blot densitometric analysis of FLAG-tagged G6PC constructs relative to tubulin control, with representative blots shown. (C) Protein expression levels of PTV Q347X were determined in Huh7 cells (n=3) and (D) HEK293 cells (n=4) by western blot densitometric analysis of V5-tagged G6PC constructs relative to tubulin control, with representative blots shown. Bars in red indicate variants that are statistical drivers of the gene-based signal. (E) Cellular localisation of V5-tagged G6PC-Q347X was assessed in Huh7 cells and overlaid with markers for the ER (calreticulin) and the trans-golgi network (TGN46). White arrows point to positions of the Golgi apparatus. Scale bar indicates 10μm. (F) Glucose-6-phosphatase activity of unglycosylated WT G6PC protein obtained from tunicamycin-treated (Tuni) HEK293 microsomes (n=2), with representative western blot of microsomal protein shown. All data presented as mean ± SEM. * p=0.01-0.05; ** p=0.001-0.01; *** p<0.001.

Together, our functional studies support p.A204S, p.R83C, and p.Q347X as functional LOF variants due to loss of G6Pase protein expression and/or activity. This results in a reduced potential to hydrolyze G6P to glucose in gluconeogenic tissues (such as in the liver and kidney), thus directly reducing FG levels and consequently lowering circulating FI levels in the plasma. Our data suggest that rare inactivating mutations in *G6PC* (such as p.R83C and p.Q347X) that cause the autosomal recessive disorder GSD1a can also modulate fasting glycemic traits within a normoglycemic range in asymptomatic heterozygous variant carriers.

### *G6PC2* alleles influence protein function by multiple mechanisms

*G6PC2*, a gene homolog of *G6PC*, is an established effector transcript at a GWAS locus which contains multiple coding variants known to influence FG and HbA1c but not FI levels (Bouatia-Naji et al., 2008; Chen et al., 2008; Mahajan et al., 2015; Soranzo et al., 2010; Wessel et al., 2015). In this current study, gene-based association signals for both FG and HbA1c were observed at the *G6PC2* locus, primarily driven by multiple coding variants (p.H177Y, p.Y207S, p.R283X, and p.S324P) (**Table S4**). We aimed to extend the investigation of coding variation in this gene, which is likely to harbor a series of functional alleles, by characterizing the four *G6PC2* coding variants above and six others, across the allelic frequency spectrum (all with single-variant *P*<0.05 for FG or HbA1c in our analyses) (**Table S4; Figure S5A**). Protein overexpression studies in the rat insulinoma cell line INS-1 832/13 and HEK293 cells revealed that seven of the G6PC2 variants characterized (including PTV p.R283X) resulted in significantly reduced protein levels (**Figures 5A and S5B-C**). In INS-1 832/13 cells, this effect was largely due to partial or total loss of the glycosylated form of the protein. In HEK293 cells, the reduction in total protein levels could be rescued when the proteasomal pathway (but not the lysosomal pathway) was inhibited, consistent with an earlier study involving a smaller subset of variants (Mahajan et al., 2015), confirming proteasome-mediated protein turnover.

**Figure 5.**
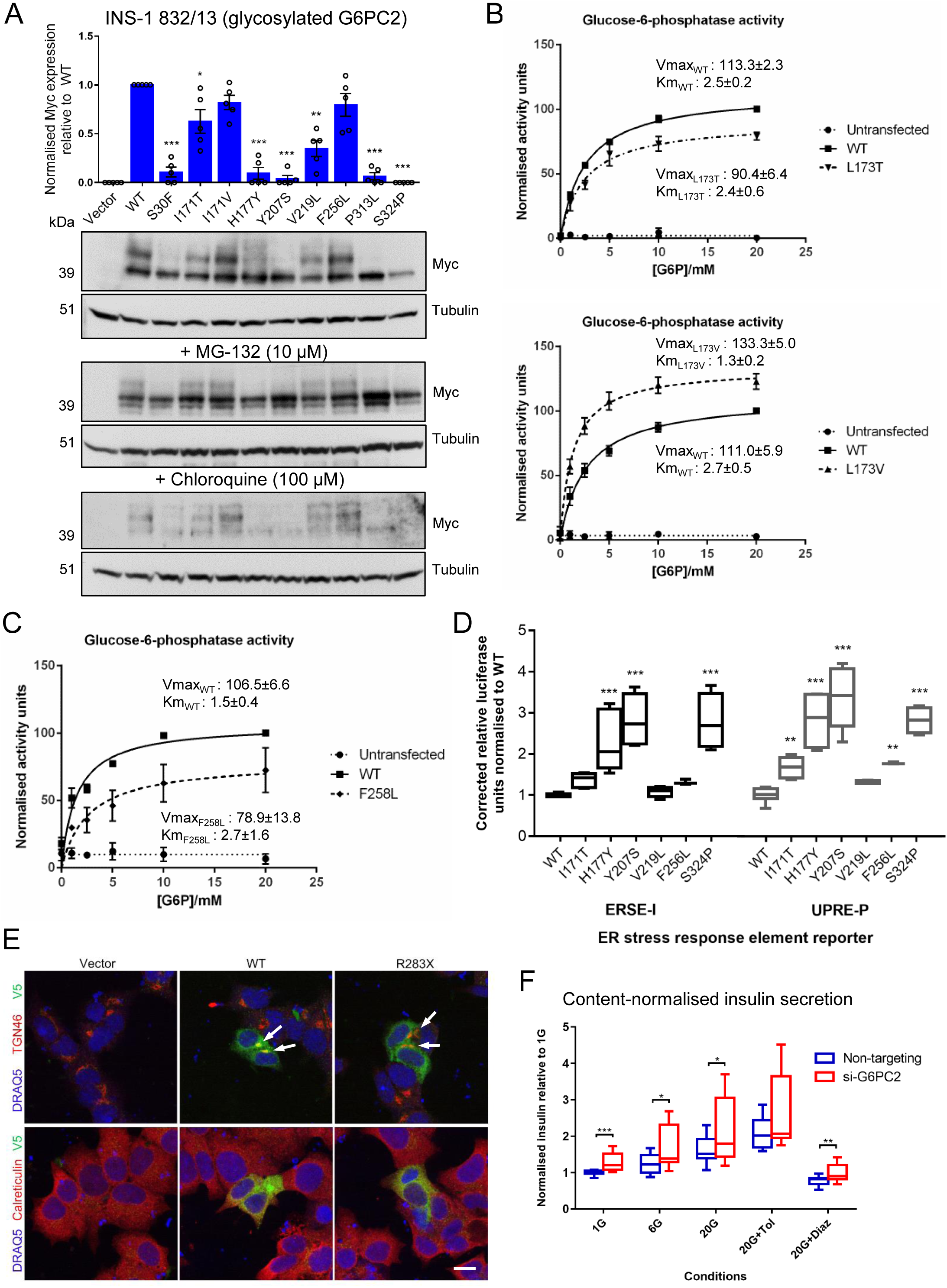
Functional characterisation of G6PC2 variant proteins and the role of G6PC2 in human beta cells. Related to Figure S5. (A) Expression levels of the glycosylated forms (upper bands only) of G6PC2 variant proteins were determined in INS-1 832/13 cells by western blot densitometric analysis of Myc-tagged G6PC2 constructs relative to tubulin control (n=5). Representative blots are shown for untreated cells together with cells treated with proteasomal inhibitor MG-132 or lysosomal inhibitor chloroquine. (B) Glucose-6-phosphatase activity of L173T and L173V variants in G6PC (proxy for I171T and I171V in G6PC2 respectively) in HEK293 against increasing glucose-6-phosphate concentrations (n=4), with mean Vmax ± SEM and Km ± SEM values shown for WT and each variant. (C) Glucose-6-phosphatase activity of F258L variant in G6PC (proxy for F256L in G6PC2) in HEK293 against increasing glucose-6-phosphate concentrations (n=3), with mean Vmax ± SEM and Km ± SEM values shown. Vmax and Km results were computed based on the Michaelis-Menten kinetic model. (D) Effect of G6PC2 WT and variant protein expression on luciferase activity driven by ER stress response elements in HEK293 cells. Relative luciferase units corrected for background activity were normalised to WT for each reporter, from n=6 across two independent experiments (except for F256L, n=3 in one experiment) using two-way ANOVA with Fisher’s LSD test comparing each variant to WT. (E) Cellular localisation of R283X in EndoC-βH1 overlaid with markers for the ER (calreticulin) and the trans-golgi network (TGN46). White arrows point to positions of the Golgi apparatus. Scale bar indicates 10μm. (F) Insulin secretion normalised to total content at basal and high glucose conditions (with and without drug treatments) following 96-120h *G6PC2* knockdown in EndoC-βH1. Unpaired two-tailed Students’ t tests were used to compare *G6PC2* knockdown to control for each condition, from n=16 across 4 independent experiments. Tol: tolbutamide; Diaz: diazoxide. All data presented as mean ± SEM. * p=0.01-0.05; ** p=0.001-0.01; *** p<0.001.

As three variants (p.I171T, p.I171V, and p.F256L) appeared to be stably expressed and processed like WT G6PC2 protein, we hypothesized that these alleles could be influencing glycemic levels through effects on protein activity. As there is a high level of conservation between the catalytic domains in G6PC and G6PC2, we adapted the G6Pase assay used earlier, to indirectly analyse the effect of the G6PC2 variants on G6Pase enzymatic activity. We assumed that the *G6PC2* alleles of interest, which mapped to the conserved regions, will give rise to the same consequence in the G6PC backbone due to the strong homology and preserved topology of both proteins. The adaptation was necessary as we were unable to detect G6PC2 activity using the same experimental conditions. First, we generated variants that mapped to equivalent sites within the G6PC protein (G6PC-p.L173T, p.L173V, and p.F258L correspond to G6PC2-p.I171T, p.I171V, and p.F256L, respectively), and then performed the enzymatic studies. Two alleles, p.L173T, p.L173V, affected the same codon and were each genetically associated with FG levels but with opposite directions of effect (**Table S4**). We found that G6PC-p.L173T exhibited ∼20% decreased activity compared to WT based on assessment of V_max_ (maximal rate of reaction), a measure of enzymatic activity (**Figure 5B**). In contrast, G6PC-p.L173V had enhanced activity through both increased V_max_ and lowered K_m_ (Michaelis constant, whereby a lower K_m_ indicates higher substrate affinity) (**Figure 5B**). Importantly, our *in vitro* observations mirrored the genetic effects on FG (β_I171T_=-0.084 mmol/l; β_I171V_=+0.131 mmol/l) and HbA1c levels (β_I171T_=-0.007%; β_I171V_=+0.093%) (**Table S4**). The G6PC-p.F258L variant also displayed impaired phosphatase activity due to reduced V_max_ and a tendency towards higher K_m_ relative to WT (**Figure 5C**), consistent with the observed glucose-lowering effects of G6PC2-p.F256L. To ensure that the observed effects of the rare variants on FG were not influenced by the common *G6PC2* variant rs560887, as was the case for a common variant V219L shown in an earlier study (Mahajan et al., 2015) which we confirm here, conditional analyses were performed conditioning on rs560887 (**Table S9**). Conditional results for p.I171T, p.I171V and p.F256L confirmed that the directions of effect for the variants remain unchanged, making it unlikely that the regulatory variant rs560887 is regulating these effects (**Table S9**). These results provided the first example of an activating allele in *G6PC2* (p.I171V) and highlighted the unique protein changes at a single codon that can give rise to a corresponding loss or gain of functional activity. These data therefore show that variations in *G6PC2* may influence FG levels through their impact on protein expression or activity.

To further characterize these variants, we set out to determine the effect of the G6PC2 LOF variants on ER integrity, given that G6PC2 is an ER-resident protein and that beta cells, which are highly-specialized secretory cells, are highly sensitive to ER stress. Specifically, we evaluated the expression of G6PC2 variant proteins on the canonical ER stress response (ERSE) and unfolded protein response (UPRE) pathways. The three G6PC2 variants which displayed relatively severe effects on protein stability (p.H177Y, p.Y207S, p.S324P) in our study were found to activate ERSE and UPRE reporter activities by ∼3-fold, in contrast to the variants p.I171T and p.F256L which exert their effects primarily on enzymatic function (**Figure 5D**). The common p.V219L variant, which reduces protein expression by approximately 50%, displayed an intermediary effect (**Figure 5D**). These results suggest that G6PC2 variant proteins, especially those that result in severe LOF due to protein instability, may also influence beta cell ER homeostasis.

In previous studies, the G6PC2-p.R283X variant has shown inconsistencies in terms of their associations with FG levels (Mahajan et al., 2015; Wessel et al., 2015). With a larger dataset we have now confirmed that this variant influences both FG and HbA1c levels (**Tables 1 and S3**). As the nonsense p.283X allele is located in the last exon of the gene and may evade NMD, we queried RNA sequencing data from human islets and observed an allelic balance in heterozygous carriers, indicating that variant transcripts are indeed likely to escape NMD and be translated (**Figure S6A**). Based on our pipeline of *in vitro* assays, we confirmed G6PC2-p.R283X loss-of-function due to reduced protein expression, failure to localize to the Golgi network, and a high likelihood of complete loss of phosphatase activity (**Figures 5A and S5D**).

In contrast to the mechanisms in play for the coding variants in *G6PC2*, the non-coding GWAS index variant at the *G6PC2* locus (rs560887) is suggested to influence expression of *G6PC2* splice variants based on previous minigene analyses in HeLa cells (Baerenwald et al., 2013; Bouatia-Naji et al., 2010). To establish whether this variant indeed influences *G6PC2* regulation in human islets, we determined its effect on *G6PC2* isoform expression. We found that in human islets, the presence of the rs560887-G allele is associated with increased expression of the full-length *G6PC2* isoform as compared with the shorter isoform lacking exon 4 (**Figure S6B**). This observation supports the hypothesis that rs560887 may alter splicing and is consistent with the association between rs560887-G and elevated FG and HbA1c levels due to increased *G6PC2* function. As the phenotypic consequence of rare coding variants can be influenced by regulatory variants on the same haplotype, we therefore performed conditional analyses to explore the relationship between rs560887 and the rare coding variants. We showed that the direction of effects of all the rare alleles in our study remained the same after conditioning on rs560887, though it is notable that the variants p.Y207S and p.R283X showed some reduction in strength of association after conditioning (**Table S9**).

### Functional assessment of *G6PC2* variants improves gene-based association analysis

We next evaluated the utility of our functional data to enhance gene-based association analyses. We showed that the gene-based signals were strengthened when the tests were informed by *in vitro* functional validation of the variants (as determined in this study) as opposed to the predictive *in silico* annotations based on the NSbroad and NSstrict masks (**Table S9, Methods**). In fact, in line with expectation, flipping the alleles in the gain-of-function variant p.I171V (which we now know acts in the opposite direction compared to other rare variants in the test), to align all alleles with the same direction of effect, augmented the strength of association for both FG (from *P*=4.34 × 10^−71^ to *P*=6.47 × 10^−78^)and HbA1c (*P*= 6.37 × 10^−30^ to *P*=6.37 × 10^−33^) in the gene burden test (**Table S9**). Improved methods of filtering variants will enhance the performance of gene-based tests and increase the likelihood of identifying true association signals, especially for those that are of borderline significance or that initially fall below the significance threshold.

### G6PC2 regulates basal insulin secretion in human beta cells

Although G6PC2 is known to be specifically enriched in pancreatic islet beta cells, its role in the regulation of human beta cell function has not been shown. Using gene knockdown studies in the human EndoC-βH1 beta cell line, we found that G6PC2-deficient cells exhibited significantly (but modestly) increased insulin secretion at low glucose (1 mM) and a trend towards increased insulin secretion at sub-maximal glucose (6 mM) levels (**Figures 5F and S5E**). When expressed as a fraction of insulin content (**Figure 5F**), insulin secretion was significantly increased across multiple glucose conditions, although this was primarily driven by reduced total insulin content in G6PC2-deficient cells by ∼15%. Overall, *G6PC2* knockdown increases glucose responsiveness at sub-threshold levels of glucose but not at maximal glucose concentration in EndoC-βH1 cells, suggesting enhancement of basal glucose sensitivity by promoting glycolytic flux at sub-stimulatory glucose concentrations, and warranting more in-depth characterization experiments.

## Discussion

We have identified novel coding variant associations with FG, FI, 2hGlu and HbA1c, across the allele frequency spectrum, and assigned these variants to their effector transcripts using available genetic and biological evidence. We further pinpointed novel loci and effector transcripts that have now been associated with T2D and other related metabolic traits since the time of our analysis. Our results revealed that 15 out of 58 glycemic trait-associated loci have evidence of association with T2D risk (**Table S2**) (Hara et al., 2014; Mahajan et al., 2018b; Williams et al., 2014). For instance, FG-associated loci *ANKH* and *STEAP2*, and HbA1c-associated *DCAF12* all associate with T2D risk (**Table S2**), providing opportunities to investigate the mechanisms through which associated variants influence both glycemic regulation within the physiological range as well as T2D pathophysiology. The FI-associated *RAPGEF3* locus is also associated with various obesity-related measures including BMI and WHR, potentially supporting our tissue enrichment analyses linking FI with adiposity.

We used this work to explore the pathways and metabolic tissues through which the associated genes influence variation in glycemic traits and highlighted those with key roles in glucose regulation and traits that act through multiple metabolic tissues, including islets, liver, fat, and in addition, exocrine pancreas, gut and kidney. Our GSEA enabled us to identify additional genes (e.g. *CTRB2*) within these tissues and pathways which were below the threshold for statistical significance in our initial discovery effort and that merit follow-up. We report an emerging role for the gut and exocrine pancreas for 2hGlu levels and potentially T2D risk through multiple analyses, consistent with current understanding that both incretins and digestive enzymes are important in controlling postprandial glucose levels (Esteghamat et al., 2019; Hart et al., 2018; Woodmansey et al., 2017). We also show that different traits are influenced by pathways operating in distinct tissues: FG is almost exclusively influenced by pathways in the endocrine and exocrine pancreas and liver, whilst FI is mediated by the insulin-sensitive tissues such as liver, kidney, and adipose tissue, indicating the importance of both insulin action and insulin clearance mechanisms. Genes expressed in muscle, also an insulin-sensitive tissue, were enriched in HbA1c-associated effector loci but not FI, though this could be due to differences in power between the two analyses. We see evidence of multiple metabolic tissues being important for HbA1c regulation, and note that the HbA1c-associated set of effector transcripts appear enriched for those that influence blood cell biology.

We have also shown for the first time that genetic variation in *G6PC*, a gene implicated in GSD1a, influences glycemic traits within the normal physiological range in heterozygote carriers. *In vitro* follow-up of the variants driving the gene-based association – p.A204S, p.R83C, and p.Q347X – confirmed that these were indeed causal LOF variants at this locus that contribute to modulation of FG and FI levels. We then reported novel rare coding variant associations for FG and HbA1c within a member of the same gene family, *G6PC2*, and expanded the allelic spectrum of reported variants to include variants affecting the same codon with both loss and gain of function alleles. Our comprehensive analysis of this locus demonstrates multiple molecular mechanisms by which variants influence protein function, including evidence from human islets that the common regulatory variant rs560887 influences *G6PC2* isoform expression, and that a rare PTV (p.R283X) evades NMD and results in a catalytically-null enzyme. Given the possiblility that the effects of any coding variants in exon 4 which are carried in *cis* with the rs560887-A allele could potentially be “diluted” due to the splicing effect, we checked whether the observed rare variant effects could be driven by rs560887 in LD by repeating the single-variant association tests with conditional analyses (**Table S9**). In our analysis, the directions of effect of the rare coding alleles do not appear to be influenced by the non-coding regulatory allele. We then used our *in vitro* data to refine existing methods for gene-based association analysis to demonstrate the value of functional data in improving their sensitivity. New developments in high-throughput functional annotation that can overcome the time-consuming nature of functional experiments will greatly facilitate such efforts (Liu et al., 2017; Tewhey et al., 2016; Ulirsch et al., 2016). Finally, to understand how loss of *G6PC2* influences FG levels, we silenced it in a human beta cell model and demonstrated increased insulin secretion at low glucose levels, in line with the genetic observations.

It has long been suspected that particular metabolic tissues are key to governing specific processes of glucose metabolism. Using human genetics, our study has explored this within an unbiased approach and has illustrated the impact of altered glycolysis in multiple metabolic tissues on various glycemic phenotypes. Uniquely, our parallel studies of G6PC and G6PC2 highlighted two homologous proteins that act through different tissues to influence glycemic traits. As G6PC is involved in hepatic glucose production it influences both FG and FI levels. Previous studies have also established a potential role for *G6PC* in influencing lipid and urate levels (Dewey et al., 2016; Sever et al., 2012). In contrast, due to its restricted expression in the islet beta cell, variants in *G6PC2* only influence FG and HbA1c due to a beta cell-driven effect. There are also notable differences in the molecular mechanisms underlying protein dysfunction: for *G6PC* variants the effect is primarily on enzymatic activity, whilst G6PC2 variants largely cause protein instability.

A limitation of the present study is that we were not able to fine-map association signals, being restricted to variants captured on the exome array, leaving many associated loci with unknown effector transcripts. Additional large-scale studies, with higher density GWAS arrays and imputation to dense reference panels, will be required for fine-mapping and further effector transcript identification.

In conclusion, we have combined human genetic discovery with pathway analysis and functional studies to uncover tissue-specific effects in common pathways that influence glycemic traits. Our findings will inform efforts to target these pathways therapeutically to modulate metabolic function.

## Methods

### LEAD CONTACT AND MATERIALS AVAILABILITY

Further information and requests for resources and reagents should be directed to and will be fulfilled by the Lead Contacts, Inês Barroso and Anna L Gloyn.

### EXPERIMENTAL MODEL AND SUBJECT DETAILS

#### Studies in humans

MAGIC (Meta-Analysis of Glucose and Insulin-related traits Consortium) was established to focus on the genetic analysis of glycemic traits in individuals without diabetes. In this MAGIC effort, non-diabetic individuals of European (85%), African-American (6%), South Asian (5%), East Asian (2%) and Hispanic (2%) ancestry from up to 64 cohorts participated. Sample sizes were up to 144,060 for HbA1c, 129,665 for FG, 104,140 for FI and 57,878 for 2hGlu. Participating cohorts and their characteristics are detailed in **Table S1**.

#### Studies in cellular models

HEK293 cells were cultured in Dulbecco’s Modified Eagle’s Medium (DMEM) (D6429, Sigma Aldrich), 10% (v/v) foetal bovine serum (FBS) (10500-064, Life Technologies), 100 U/ml penicillin and 100 µg/ml streptomycin (15140122, Life Technologies). Huh7 cells were cultured in DMEM (31885, Life Technologies), 10% (v/v) FBS, 100 U/ml penicillin and 100 µg/ml streptomycin. INS-1 832/13 cells were cultured in Roswell Park Memorial Institute-1640 (RPMI-1640) media (R0883, Sigma Aldrich), 10% (v/v) FBS, 100 U/ml penicillin and 100 µg/ml streptomycin, 2 mM L-glutamine (25030081, Life Technologies), 1 mM sodium pyruvate (S8636, Sigma Aldrich), 10 mM HEPES (H3537, Sigma Aldrich), 50 µM 2-mercaptoethanol (Life Technologies). EndoC-βH1 cells were cultured in DMEM (31885, Life Technologies), Bovine Serum Albumin (BSA) fraction V (10775835001, Roche), 100 U/ml penicillin and 100 µg/ml streptomycin, 2 mM L-glutamine, 50 µM 2-mercaptoethanol, 10 mM nicotinamide (Sigma Aldrich), 5.5 µg/ml transferrin (Sigma Aldrich) and 6.6 ng/ml, sodium selenite (Sigma Aldrich). All cell lines were tested negative for mycoplasma contamination using the MycoAlert Assay kit (Lonza). Cells were maintained at 37°C and 5% CO_2_.

## METHOD DETAILS

### Studies in humans

#### Phenotypes

Studied outcomes were FG (mmol/L), Ln-transformed FI (pmol/L), 2hGlu (mmol/L) and HbA1c (% of hemoglobin). Glycemic measurements are described in detail for each contributing cohort in **Table S1**. Individuals with diagnosed or treated diabetes, or those with diabetes on the basis of FG (≥ 7 mmol/L), 2hGlu ((≥ 11.1 mmol/L) and/or HbA1c (≥ 6.5%) were excluded from analyses.

#### Genotyping and QC

The Illumina HumanExome BeadChip is a genotyping array containing variants that have been observed in sequencing data of ∼12,000 individuals. Non-synonymous variants seen at least three times across at least two datasets were included on the exome chip. More lenient criteria were used for splice and nonsense variants. Besides the core content of protein-altering variants, the exome chip contains additional variants including common variants identified in GWAS, ancestry informative markers, mitochondrial variants, randomly selected synonymous variants, HLA tag variants and Y chromosome variants. In this study we analysed association with glycemic traits of 247,470 autosomal and X chromosome variants present on the exome chip. Genotype calling and quality control were performed following protocols developed by the UK Exome Chip or CHARGE consortium (Grove et al., 2013). The exact genotyping array, calling algorithm and QC procedure used by each cohort are depicted in **Table S1**.

#### Annotation and functional prediction of variants

Annotation of the exome chip variants was performed using the Ensembl Variant Effect Predictor v78. *In silico* functional prediction from SIFT, Polyphen HumDiv, Polyphen HumVar, LRT and MutationTaster was added using dbNSFP v2.9 (Liu et al., 2013; Yourshaw et al., 2015).

#### Statistical analyses

##### Single variant analyses

Individual cohorts ran linear mixed models using the raremetalworker (v 4.13.2) or rvtests (v20140723) software (**Table S1**). For each glycemic outcome, analyses were performed using an additive model for the raw and the inverse normal transformed trait. In the manuscript and in all tables and figures effect estimates and standard errors are for the raw trait, while the p-values are from the inverse normal transformed trait analyses. Analyses were adjusted for age, sex, BMI, study-specific number of PCs and other study-specific covariates (**Table S1**). Raremetal (v4.13.7 or higher) was used to combine results by fixed-effect meta-analyses. Variants with *P* < 10^−4^ for deviation from Hardy-Weinberg equilibrium or with call rate < 0.99 in individual cohorts were excluded from meta-analyses. In single variant analyses, the threshold for significance was *P* < 2.2×10^−7^ for coding variants (stop-gained, stop lost, frameshift, splice donor, splice acceptor, initiator codon, missense, in-frame indel and splice region variants). These *P*-value thresholds were based on a Bonferroni correction weighted by the enrichment for complex trait associations among the different functional annotation categories (Mahajan et al., 2018b; Sveinbjornsson et al., 2016). Significant association signals located more than 500 kb from any variant already known to be associated with the trait at the time of analysis (May 2015) were considered novel for the trait.

##### Gene-based analyses

In addition, raremetal was used to perform gene-based burden and sequence kernel association (SKAT) tests. For both burden and SKAT tests, two *in silico* masks for inclusion of variants in the test were used: NSstrict and NSbroad. The NSstrict mask includes PTVs (splice donor, splice acceptor, stop gained, frameshift, stop lost or initiator codon variant) OR variants that are missense and predicted to be damaging by five prediction algorithms (SIFT, Polyphen HumDiv, Polyphen HumVar, LRT, MutationTaster). The NSbroad mask additionally includes missense variants predicted to be damaging by at least one of the five prediction algorithms AND that have a MAF<1% in each ancestry group. These MAFs were derived from our single variant HbA1c meta-analyses results (N up to 144,060). For *G6PC2*, we also used masks filtering on functional variants that have been determined *in vitro* to influence protein expression or function. The *P*-value threshold for significance in gene-based analyses was 2.5×10^−6^ (Bonferroni correction for 20,000 genes).

##### Conditional analyses

Approximate conditional analyses were performed using Raremetal v 4.13.8. At known glycemic trait loci, if previously known GWAS index variants (or good proxies) were present on the exome chip, significant lead coding variants were conditioned on these known index variants and vice versa to identify distinct coding variant signals. At novel loci, to identify additional distinct associated variants, analyses were performed conditioning on the most significant variant at the locus. These analyses were repeated by including the next most significant and distinct associated variant until no exome- or genome-wide significantly-associated variants were left at the locus. For gene-based signals, to identify the variants driving the signal, analyses were performed conditioning on the variant with the most significant p-value that was included in the mask. These analyses were repeated including the next most significant variant until association at the gene was attenuated (*P* > 0.05). If there were both gene-level and known or novel single variant associations at the same locus (within 500 kb), we additionally conditioned on the associated single variant to assess whether the gene-based association was distinct from the single variant association.

#### Putative effector transcript identification

To identify putative effector transcripts, at known glycemic trait loci we considered the transcript a putative effector transcript if there was a distinct coding variant signal (still meeting the threshold for significance of *P* < 2.2×10^−7^ after conditioning on the non-coding GWAS index variant, for details on these conditional analyses methods refer to the *conditional analyses* methods section above). Coding variant associations at novel loci were followed up on in published GWAS results with higher density coverage (Manning et al., 2012; Wheeler et al., 2017a). If the coding variant was present in the GWAS results, approximate conditional analyses were performed using GCTA (Yang et al., 2012). If the GWAS index variant signal was abolished by conditioning on the coding variant, we considered this as evidence supporting the transcript as a putative effector transcript. If the both the GWAS index variant and the coding variant signals were attenuated, the results were considered uninformative and we considered the transcript in light of other data. We additionally utilized published data to classify effector transcripts, including (1) fine-mapping results from comparable T2D efforts (Mahajan et al., 2018b) and a body of literature establishing a role in glucose metabolism or red blood cell biology (for HbA1c) for certain genes that mapped within our loci. Significant gene-based associations driven by multiple coding variants within a single gene, in particular where an impact on protein expression or function could be demonstrated, were considered strong evidence for the determination of effector transcripts. Combining these approaches, we attempted to identify effector transcripts at each locus, and we classified their likelihood of being correct depending on the strength of the evidence. Those effector transcripts where there was strong evidence from reciprocal conditional analysis or support from published data for the relevant glycemic trait or phenotype were labelled “gold”; those where the effector transcript could not be defined by conditional analysis (either because it was inconclusive or due to lack of data) but where there was strong biological plausibility for a given gene at the locus were labelled “silver”; those where we had some tentative evidence but that was not strong enough to warrant a “silver” classification were labelled “bronze”, and the remainder were left with an unknown effector transcript. Effector transcript classification into “gold”, “silver” and “bronze” was undertaken independently by four of the authors and the highly concordant consensus score was given (**Table S6**).

#### GeneMANIA network analysis

For network analyses, we used GeneMANIA (v3.5.1), a network approach that searches many large, publicly-available biological datasets to find related genes. These include protein-protein, protein-DNA and genetic interactions, pathways, reactions, gene and protein expression data, protein domains and phenotypic screening profiles. Briefly, GeneMANIA uses a label propagation algorithm for predicting gene function given the composite functional association network (calculated from the databases selected). The weights needed for the label propagation method to work are selected at the beginning of the process. In our case, and according to the defaults, we weighted the network using linear regression, to make genes in the input list interact as much as possible with each other. We analyzed all non-synonymous variants for each locus with a cut-off of association *P*<1×10^−5^ with any trait (input genes). We performed four network analyses: (1) HbA1c-associated variants only, (2) FI-associated variants only, (3) FG-associated variants only, and (4) 2hGlu-associated variants only.

We selected the 50 default databases to create the composite network, and we allowed the method to find at most 50 genes that are related to our query input list. The resultant networks were investigated to find enriched Gene Ontology (GO) terms and Reactome Pathways. Gene Set Enrichment (GSE) of networks and sub-networks were assessed with ClueGO (Bindea et al., 2009) using GO terms and Reactome gene sets (Croft et al., 2014). The enrichment results were grouped using a Cohen’s Kappa score of 0.4, and terms were considered significant with a Bonferroni-adjusted p-value<0.05, provided that there was an overlap of at least three network genes in the relevant GO gene set when calculating GO enrichment. For the pathway selection (Reactome), we set a threshold that the network genes should represent at least 4% of the pathway. These values were applied given the recommended defaults when running ClueGO (Bindea et al., 2009). Cohen’s Kappa statistic was used to measure the gene-set similarity of GO terms and Reactome pathways and allowed us to group enriched terms into functional groups to improve visualization of enriched pathways. We used all genes with GO annotations and at least one interaction in our network database as the background set.

#### Gene set enrichment analysis (GSEA)

For GSEA, we used EC-DEPICT, an extension of the GWAS GSEA method DEPICT (Pers et al., 2015). EC-DEPICT has been described elsewhere (Marouli et al., 2017; Turcot et al., 2018). Briefly, the key feature of EC-DEPICT is the use of “reconstituted” gene sets, which are gene sets collected from many different databases (e.g. canonical pathways, protein-protein interaction networks, and mouse phenotypes) that have been extended based on large-scale microarray co-expression data (Fehrmann et al., 2015; Pers et al., 2015).

Six groups of variants were analyzed: (1) HbA1c-associated variants only, (2) FI-associated variants only, (3) FG-associated variants only, (4) 2hGlu-associated variants only, (5) all trait-associated variants, and (4) all trait-associated variants except for HbA1c (see **Methods**). For each trait, we clumped the European summary statistics (+/- 500 kb on either side). Then, the most significant nonsynonymous variant for each locus was included in the analysis, with a cut-off of *P*<10^−5^. Annotations from the CHARGE consortium were used to assign variants to genes (see **URL**). After GSEA, highly correlated gene sets were grouped by affinity propagation clustering of all 14,462 gene sets (Frey and Dueck, 2007) into “meta-gene sets” using SciKitLearn.clustering.AffinityPropagation version 0.17 (Abraham et al., 2014). For all visualizations, the gene set within a meta-gene set with the best enrichment *P*-value was used; heat maps were created with the ComplexHeatmap package in R (Gu et al., 2016).

##### URL

CHARGE Consortium ExomeChip annotation file (v6): http://www.chargeconsortium.com/main/exomechip/

###### Method and choice of data for permutations

We performed the EC-DEPICT analysis as described elsewhere (Marouli et al., 2017; Turcot et al., 2018). All analyses are based on a group of 14,462 “reconstituted” gene sets, which contains a z-score for probability of gene set membership for each gene (for details, see (Fehrmann et al., 2015; Pers et al., 2015)).

Briefly, the basic EC-DEPICT method is as follows. We first obtain a list of significant input variants (the most significant nonsynonymous variant per locus) and then map variants to genes based on annotations from the CHARGE consortium (see **URL**). For each gene set, we obtain the gene set membership z-scores for all trait-associated input genes and sum them to generate a test statistic. We then take 2,000 permuted ExomeChip association studies (described in more detail below) and calculate the average permuted test statistic for that gene set, as well as the permuted standard deviation. For each permutation, the number of top genes we take as “input genes” is matched to the actual observed number of input genes. We then calculate (observed test statistic – average permuted test statistic)/(permuted standard deviation) to generate a z-score, which is converted to a p-value via the normal distribution. False discovery rates were calculated by comparing the observed p-values to a permuted *P*-value distribution generated with an additional set of 50 permuted association studies.

The permuted ExomeChip association studies are conducted by (1) generating 2,200 sets of normally distributed phenotypes and (2) using these randomly generated phenotypes to conduct 2,200 association studies with real ExomeChip data. Using these permutations to adjust the observed test statistics corrects for any inherent structure in the data (e.g. that pathways made up of longer genes may be more likely to come up as significant by chance).

For these analyses, we first generated permutations based on ExomeChip data we had used previously for this purpose: 11,899 samples drawn from three cohorts (Malmö Diet and Cancer [MDC], All New Diabetics in Scania [ANDIS], and Scania Diabetes Registry [SDR]). For simplicity, we refer to these cohorts as the “Swedish permutations.”

As part of our GSEA pipeline, we remove input trait-associated variants that are not present in the permuted data to ensure that all variants are appropriately modeled. When using the Swedish permutations, this generally results in removing a substantial fraction of the variants, especially of the very rarest variants (due to the smaller sample size of the Swedish data relative to the data being analyzed). We have previously observed that this filtering can actually improve the GSEA signal, possibly due to more heterogeneous biology or a higher false-positive rate in these very rare variants (Turcot et al., 2018). However, in this case, we observed that in performing this filtering, we excluded variants in several known monogenic disease genes, such as *HNF1A* and *SLC2A2*. Therefore, we wished to repeat the analysis with a set of permutations which would allow us to retain these variants. We thus repeated the analysis with a second set of permutations consisting of 152,249 samples from the UK Biobank (referred to as the “UKBB permutations”). The larger sample size in the UKBB permutations means more variants are present and can therefore be included in the analysis.

###### Concordance of results from two different sets of permuted distributions across phenotypes

For completeness, we report the results from the use of both sets of permutations. We note that the results are strongly concordant. The larger number of significant gene sets reported based on the UK Biobank permutations is generally a combination of 1) overall improved power (i.e. more variants are included) and 2) the inclusion of variants in key driver genes absent in the Swedish permutations, encompassing both the monogenic genes mentioned above (e.g. *SLC2A2*) and additional genes with clearly relevant biology (e.g. *CTRB2, SLC30A8*). The results from both sets of permutations are summarized below. For all analyses, “significance” refers to a false discovery rate of <0.05.

###### All-trait analysis

After filtering, 78 input genes were included for the analysis with the UKBB permutations and 60 for the analysis with the Swedish permutations. (Note that the difference in the number of input genes is due to the presence of a larger number of input variants in the UKBB permutations – see above). We found 234 significant gene sets in 86 meta-gene sets based on the UKBB permutations (**Figure S2**) and 133 gene sets in 51 meta-gene sets based on the Swedish permutations (**Figure S3**). The correlation between the UKBB and Swedish analyses was r = 0.902, *P*< 10^−300^.

###### All-traits-except-HbA1c analysis

After filtering, 45 input genes were included for the analysis with the UKBB permutations and 33 for the analysis with the Swedish permutations. We found 128 significant gene sets in 53 meta-gene sets based on the UKBB permutations (**Figure S2**) and 45 significant gene sets in 18 meta-gene sets based on the Swedish permutations (**Figure S3**). The correlation between the UKBB and Swedish analyses was r = 0.882, *P*< 10^−300^.

###### HbA1c-only analysis

After filtering, 41 input genes were included for the analysis with the UKBB permutations and 33 for the analysis with the Swedish permutations. We found 191 significant gene sets in 73 meta-gene sets based on the UKBB permutations (**Figure S2**) and 120 gene sets in 41 meta-gene sets based on the Swedish permutations. (**Figure S3**). The correlation between the UKBB and Swedish analyses was r = 0.936, *P*< 10^−300^.

###### FG-only analysis

After filtering, 26 input genes were included for the analysis with the UKBB permutations and 22 for the analysis with the Swedish permutations. We found 106 significant gene sets in 39 meta-gene sets based on the UKBB permutations (**Figure S2**) and 48 significant gene sets in 15 meta-gene sets based on the Swedish permutations (**Figure S3**). The correlation between the UKBB and Swedish analyses was r = 0.939, *P*< 10^−300^.

###### 2hGlu-only analysis

After filtering, 12 input genes were included for the analysis with the UKBB permutations and 7 for the analysis based on the Swedish permutations. We found 56 significant gene sets in 17 meta-gene sets based on the UKBB permutations (**Figure S2**), with no significant gene sets based on the Swedish permutations. The correlation between the UKBB and Swedish analyses was r = 0.787, *P*< 10^−300^.

###### FI-only analysis

After filtering, 11 input genes were included for the analysis with the UKBB permutations and 8 for the analysis with the Swedish permutations. There were no significant gene sets from either analysis. The correlation between the UKBB and Swedish analyses was r = 0.860, *P*< 10^−300^.

##### Visualization

As in previous work (Marouli et al., 2017; Turcot et al., 2018), we have included all trait-associated variants in the heat maps, even if they were excluded from the analysis (e.g. because they were absent in the permutations or did not have a nonsynonymous annotation in the CHARGE annotation file). This is because we assume that if the genes harboring those variants have strong predicted membership in significantly trait-associated gene sets, they are still good candidates for prioritization. In fact, this may be even stronger evidence in favor of these genes because they did not contribute to the enrichment analysis and therefore their prioritization is independently derived (and provides even more support to the implicated biology).

#### Tissue enrichment analysis

We analysed identified genes (all 51 effector transcripts) for tissue enrichment using publicly available expression data from the GTEx project, version 7 and publicly-available islet expression data (van de Bunt et al., 2015). We use transcripts per million (TPM) values for gene level analyses. We have excluded genes from non-coding proteins and only used those with unique HGCN IDs (*n* = 20,160). We ranked all genes by median TPM across all samples, and generated 10,000 permutations of each gene set list (golden, silver, and bronze) by selecting a random gene for each entry in the gene set within ± 150 ranks of the transcript for that gene. For each sample in GTEx tissues, the TPM values were converted into ranks for that gene, and sums of ranks within each tissue were computed for each gene. We calculated enrichment p-values for each tissue by taking the total number of instances when the gene list of interest had a lower sum of ranks than the permuted sum of ranks (divided by the total number of permutations). To check that our results were not driven by sample size differences in each of the 45 analyzed GTEx tissues and islet tissue, we applied a ‘downsampling’ strategy. We performed 3 different downsampling analyses with 100, 150 and 175 samples chosen at random from each of the selected GTEx tissues and compared them to the results obtained with the whole dataset. During each downsampling round, we only used those tissues with at least the target number of samples (100, 150 or 175) because the random selection was performed under a no-replacement condition. Our results were robust to sample size differences and the trends observed were not driven by differences in sample sizes across tissues.

#### RNA-sequencing of human islets

RNA from human islet samples (n=150) was sequenced on Illumina HiSeq2000 as previously described (van de Bunt et al., 2015). Allele-specific expression was assessed using MAMBA (Pirinen et al., 2015). For the isoform effects, all protein-coding and lincRNA transcripts from GRCh37 (Ensembl release 75) were quantified using Salmon v0.8.1 (Patro et al., 2017). Isoform ratios were calculated by dividing each transcript’s expression by the total expression of that gene. For the QTL analysis, all isoforms with expression in < 50% or all samples, with no variance between samples, only 0 or 1 fractions across samples, or those originating from non-autosomal chromosomes were removed. Ratios of the remaining transcripts were rank-transformed to normality. Subsequently, 30 PEER factors to account for potential sources of non-genetic noise were derived from the normalized isoform ratios, and, together with three genotype principal components and a sex covariate, were used in the QTL analysis using FastQTL (Ongen et al., 2016). Finally, the resulting beta-approximated p-values were adjusted for multiple testing across all tested transcripts using the Benjamini-Hochberg procedure.

#### Studies in cellular models

##### Site directed mutagenesis

Human *G6PC* (NM_000151.3) and *G6PC2* cDNA (NM_021176.2) within a pCMV6-Entry vector (with a C-terminal Myc-FLAG-tag) was purchased from OriGene (RC215623 and RC211146 respectively). For the study of PTVs, an N-terminal V5 tag sequence (5’-GGTAAGCCTATCCCTAACCCTCTCCTCGGTCTCGATTCTACG-3’) was cloned into the OriGene vectors. Single nucleotide substitutions were generated in the *G6PC* or *G6PC2* coding sequence using Quikchange II Site-Directed Mutagenesis (Agilent). All mutations were verified by Sanger sequencing and in each case, only the desired nucleotide changes were introduced.

##### Western blot analyses

Western blots were performed on total protein lysates collected from human HEK293 and Huh7 cells and rat INS-1 832/13 cells transfected with each wild type or mutant *G6PC*/*G6PC2* construct using Lipofectamine 2000 (Invitrogen) according to manufacturer’s instructions. All cell lines were tested negative for mycoplasma contamination. For the inhibition of cell proteolysis, cells were treated with 10μM MG-132 (Calchembio) or 100μM chloroquine (Sigma) for 15h. For inhibition of N-linked glycosylation, cells were treated with 1µg/ml tunicamycin (Sigma) for 15h. Cells were collected 36-48h after transfection and homogenized in lysis buffer. Cell lysates were separated by 4–12% SDS-PAGE (Bio-Rad/Invitrogen). The antibodies used for determining recombinant G6PC/G6PC2 expression were: anti-FLAG M2 (Sigma, F1804), anti-V5 (Invitrogen, 46-0705) or anti-myc 4A6 (Millipore, 05-724). A β-tubulin antibody (Santa Cruz, sc-9104) was used as a loading control. Secondary antibodies specific to mouse or rabbit IgG were purchased from Thermo Fisher Scientific. Protein bands were detected using the western enhanced chemiluminescence substrate (BioRad).

##### Immunofluorescence microscopy

Human HEK293, Huh7 and EndoC-βH1 cells were transfected using FuGene 6 transfection reagent (Promega) according to manufacturer’s instructions, in 4-well chamber slides (BD Biosciences). After 48h, cells were fixed with 4% paraformaldehyde in PBS, permeabilized with 0.05% Triton X-100 in PBS and blocked with 10% BSA in PBS-Tween 20. Double immunostaining of cells was carried out using anti-FLAG M2 (Sigma, F1804) or anti-V5 (Invitrogen, 46-0705), together with anti-calreticulin (Thermo, PA3-900) or anti-TGN46 (Sigma, T7576) primary antibodies. The secondary antibodies used were anti-mouse Alexa Fluor 488 and anti-rabbit Alexa Fluor 568, both from Life Technologies. DRAQ5 fluorescent probe (Thermo Fisher Scientific) was applied at 20μM as a far-red nuclear stain. Finally, slides were mounted with ProLong Gold antifade reagent (Life Technologies) and visualized on a BioRad Radiance 2100 confocal microscope with a 60X 1.0 N.A. objective. Images were acquired with different laser settings that were optimized for each sample and therefore fluorescent intensities are not comparable across samples.

##### Glucose-6-phosphatase activity of microsomal samples

For the collection of microsomes, HEK293 cells were transfected with 12.5 µg of wild type or mutant *G6PC* construct in 10 cm dishes using Lipofectamine 2000 (Invitrogen). For the study of *G6PC2* variant activity, site directed mutagenesis was carried out within the conserved sequence regions on the *G6PC* background. Cells were cultured in Dulbecco’s modified Eagle’s medium (DMEM) supplemented with 10% fetal bovine serum, 100 U/ml penicillin, and 100 µg/ml streptomycin. At least two dishes of cells per condition were harvested 48h after transfection and scraped into 0.25M sucrose-5 mM HEPES buffer (SH) followed by several rinses in SH. Cells were mechanically homogenized using a Potter-Elvehjem glass tissue grinder and Teflon pestle, followed by 12 passes through a 27-gauge syringe needle. The homogenate was subjected to centrifugation at 10,000 g for 10 min and the supernatant (post-nuclear fraction) was further centrifuged at 100,000 g for 1h in a TLA 100.4 rotor in an Optima TLX ultracentrifuge (Beckman Coulter). A pellet containing the microsomal fraction was obtained and resuspended in SH. An aliquot of each microsomal sample was lysed in lysis buffer for protein quantification using the Bradford reagent (Bio-Rad) and analysed by western blot (BioRad) to determine the relative levels of recombinant G6PC WT and variant expression. For glucose-6-phosphatase assays, matched amounts of G6PC WT or variant protein (approximately 1-5µg of microsomal protein) were each incubated in a 200 µl reaction mix containing 100 mM MES pH 6.5, and 0 to 20 mM G6P at 37°C for 8 min. The reaction was terminated by addition of 20% trichloroacetic acid and centrifuged at 4,000 rpm for 10 min in a microcentrifuge. The supernatant was mixed in equal parts with a Taussky-Shorr colour reagent (1% ammonium molybdate, 5% iron(II) sulphate heptahydrate in 0.5M H_2_SO_4_) for 7.5 min before measuring absorbance at 660 nm on a spectrophotometer (Molecular Devices Ltd). The amount of phosphates detected was calculated using a KH_2_PO_4_ standard curve. Results were expressed as mean normalised activity (nmol/mg/min) relative to the activity of wild type at 20 mM G6P for every experiment. Finally, Michaelis-Menten enzyme kinetic analysis and paired t tests of determined kinetic constants were carried out on GraphPad Prism 6.0.

##### ER stress response reporter assays

HEK293 cells were co-transfected with *G6PC2* WT or variant constructs and pGL3-Promoter constructs containing ER stress response elements (ERSE-I and ERSE-II) or UPR elements (UPRE-P and UPRE-W) using the FuGene 6 transfection reagent (Promega). A Renilla luciferase gene-containing pRL-CMV was also co-transfected as an internal transfection control. Cells were lysed in passive lysis buffer (Promega) and assayed using the Dual Luciferase Assay System (Promega).

##### Insulin secretion analysis in EndoC-βH1 cells

Gene knockdown was carried out on EndoC-βH1 cells using ON-TARGETplus siRNA (Dharmacon, GE Healthcare) and Lipofectamine RNAiMAX (Life Technologies) at a final concentration of 25 nmol/L siRNA. For static incubation experiments, cells were placed in 2.8 mM glucose DMEM (11966, Gibco by Life Technologies) overnight. Cells were starved in 0 mM glucose medium for 1h the following day, then stimulated in DMEM containing 1 mM glucose, 6 mM glucose, 20 mM glucose, 20 mM glucose with 100 µM tolbutamide (Sigma Aldrich) or 20 mM glucose with 100 µM diazoxide (Sigma Aldrich) at 37°C for 1h. Each condition was carried out in triplicate or quadruplicate wells within each experiment. Viable cell count was measured using the CyQUANT Direct Cell Proliferation Assay kit (C35012, Thermo Scientific). All cell count values were expressed as fluorescent units normalised to mean cell count at 1 mM glucose. Cells were extracted for analysis of insulin content with cold HCl-ethanol (Sigma Aldrich). Insulin levels were measured using the human insulin AlphaLISA detection kit (AL204C, Perkin Elmer).

## QUANTIFICATION AND STATISTICAL ANALYSIS

Western blot bands for protein expression studies were quantified by densitometry analysis using ImageJ and densitometric data between G6PC/G6PC2 WT and each variant from 3-5 independent experiments were compared using two-tailed paired Students’ t tests. For enzymatic assays, mean differences in activity between G6Pase WT protein and each variant protein for the substrate G6P were compared using two-tailed unpaired Students’ t tests of the determined kinetic constants V_max_ and K_m_. For the analysis of ER stress luciferase activity data, a two-way analysis of variance (ANOVA) was applied to compare mean fold difference in reporter activity between G6PC WT and variant. For gene expression analyses, G6PC2 KO and control cells were analysed using two-tailed unpaired Students’ t tests. For the analysis of insulin secretion data, mean differences between *G6PC2* KO cells and control cells for each condition or time point were compared using two-tailed unpaired Students’ t tests. Plotting of graphs and statistical analyses were carried out on GraphPad Prism 6.0 or 7.0. A P value <0.05 was considered significant.

## Acknowledgements

HepG2 cells were the kind gift of the Tomlinson group (OCDEM), INS-1 832/13 cells were the kind gift of Jochen Lang (Université de Bordeaux), and EndoC-βH1 cells were gifted by Raphael Scharfmann (INSERM).

---

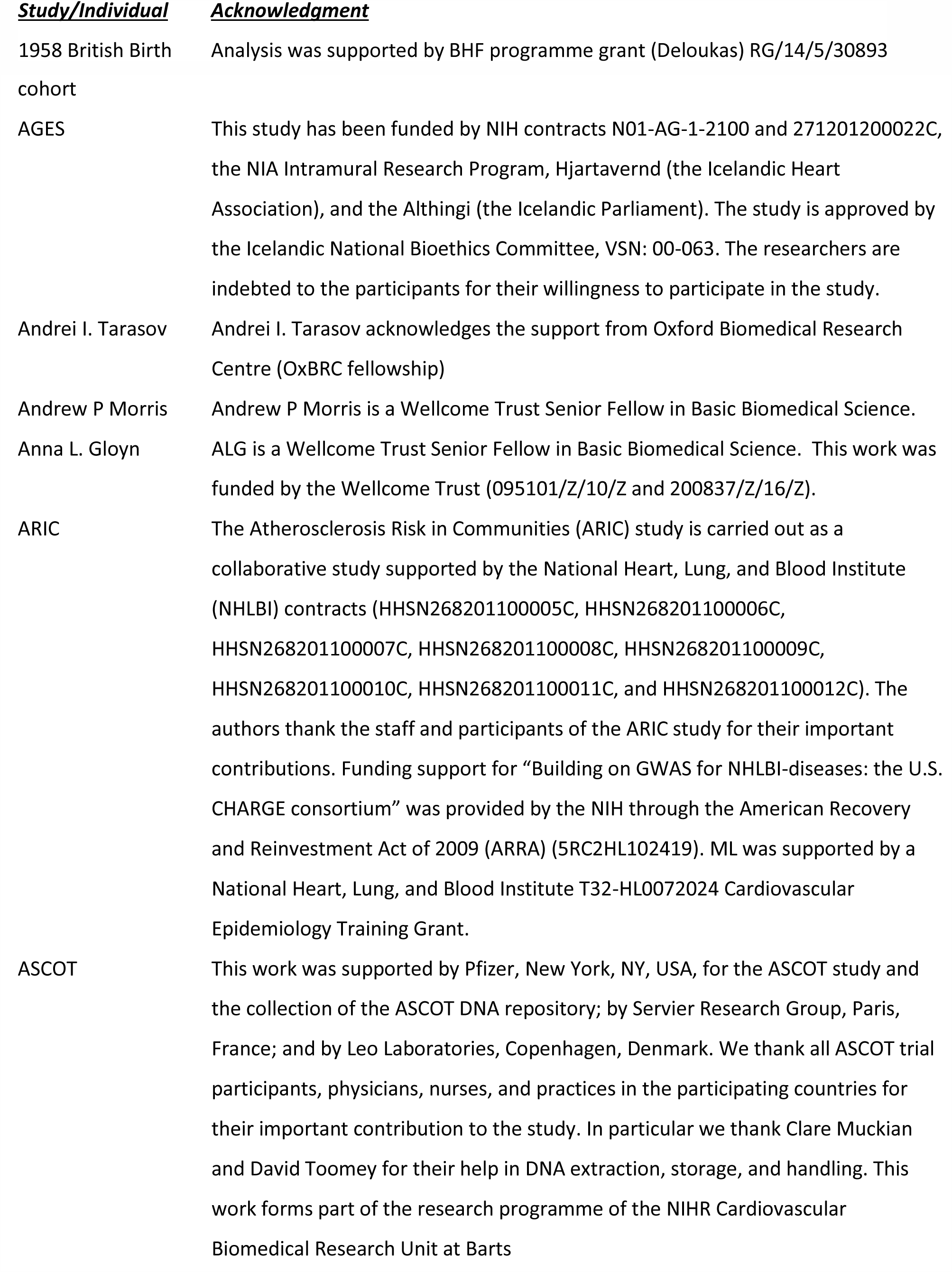
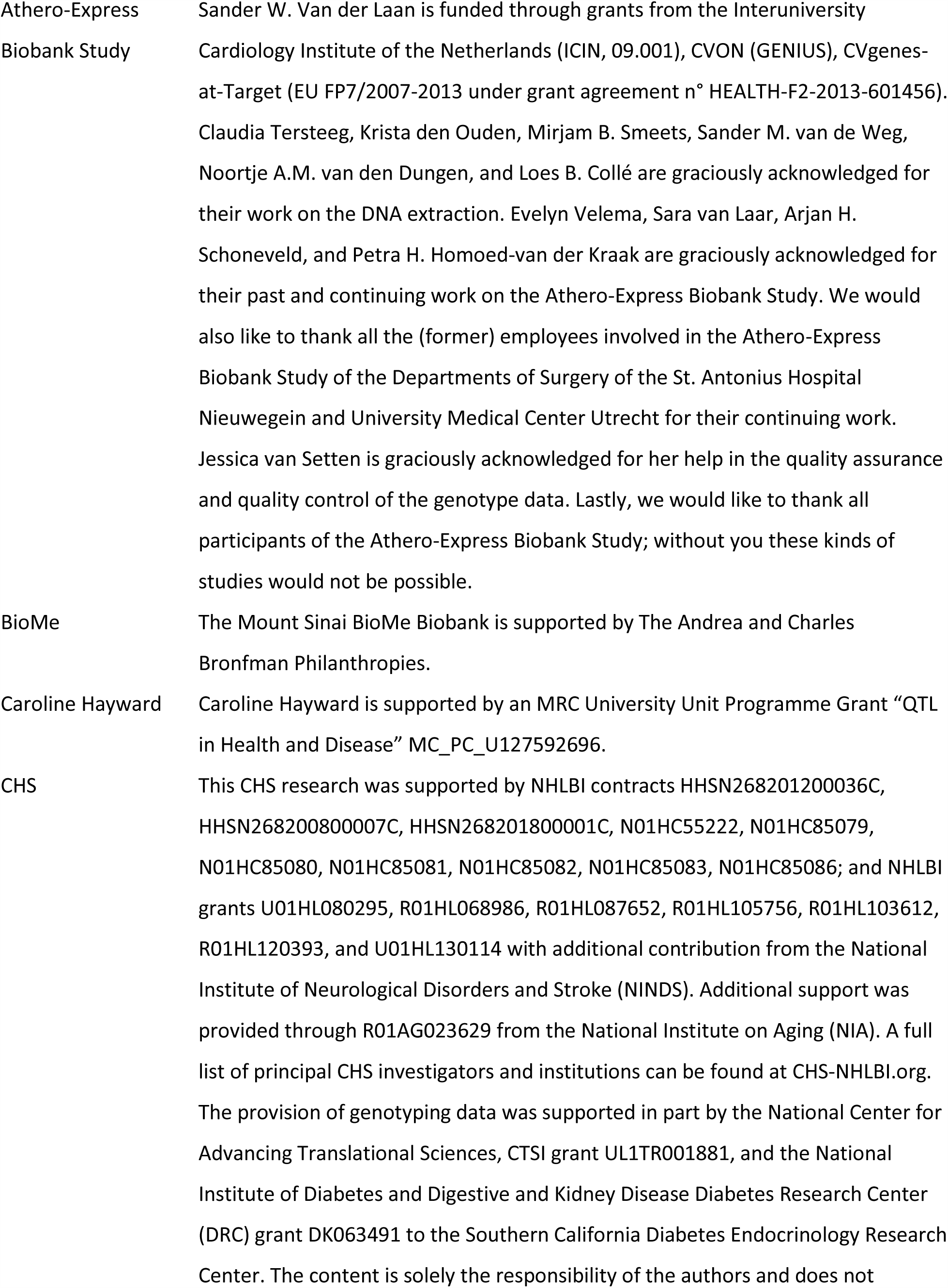
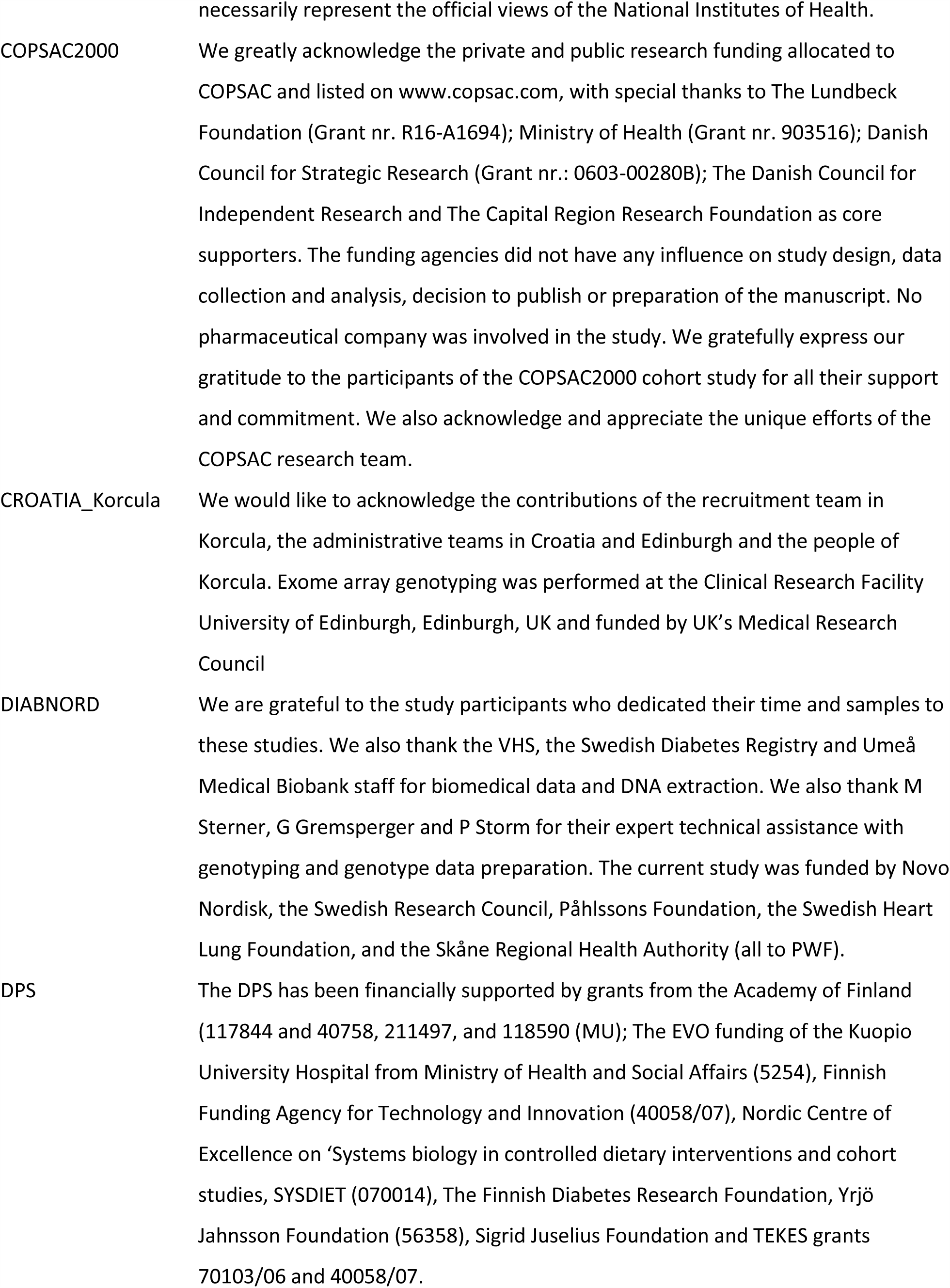
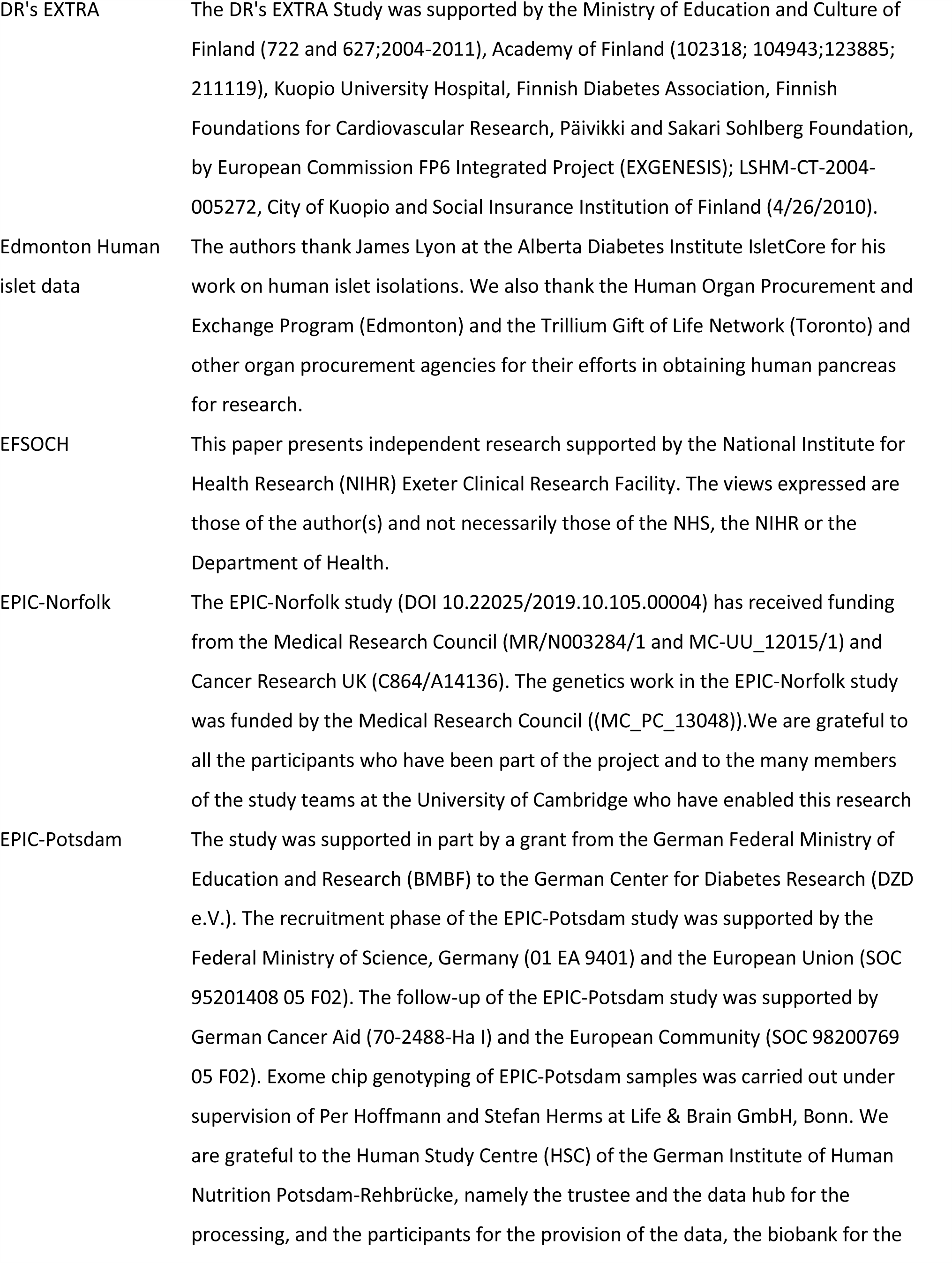
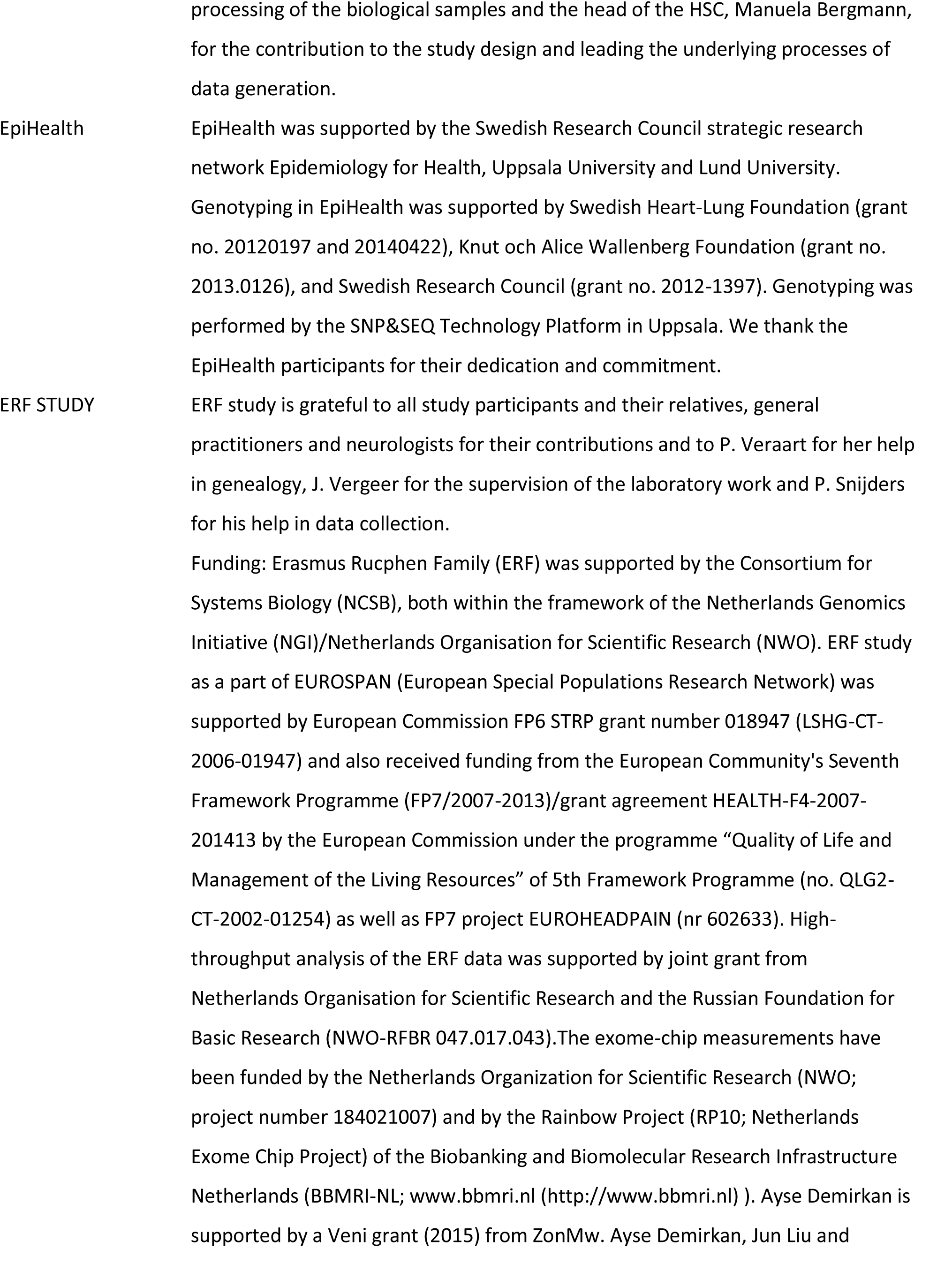
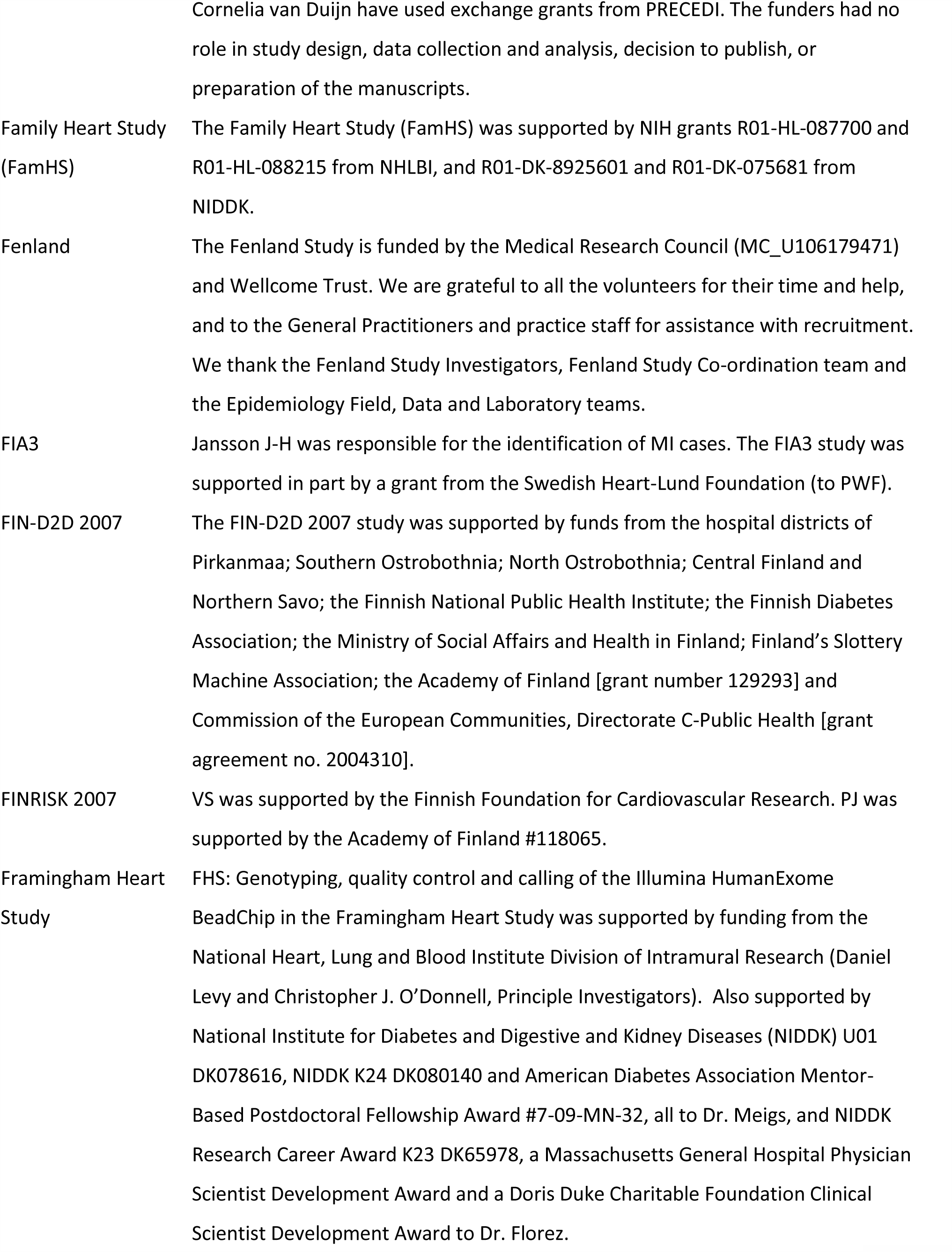
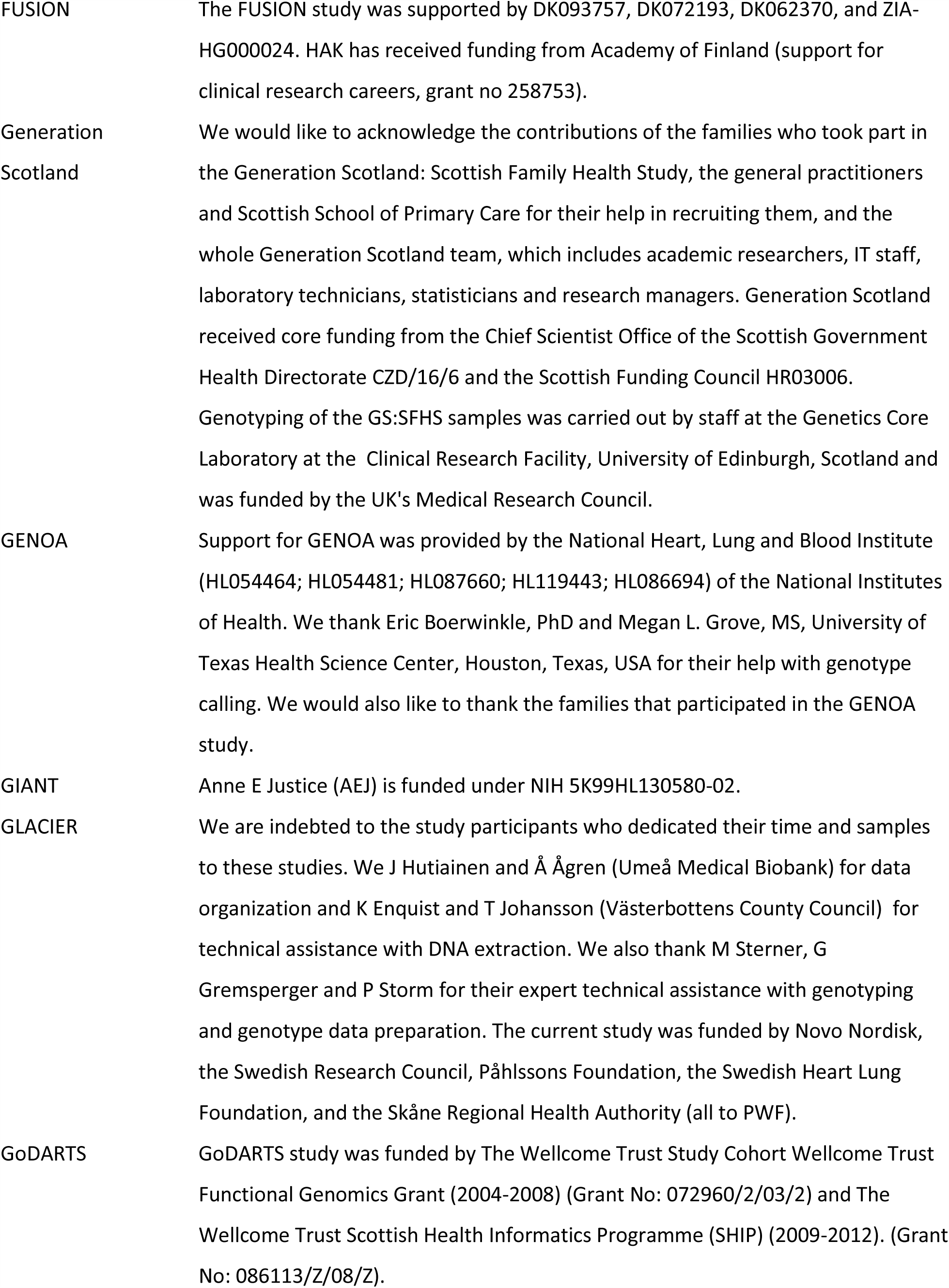
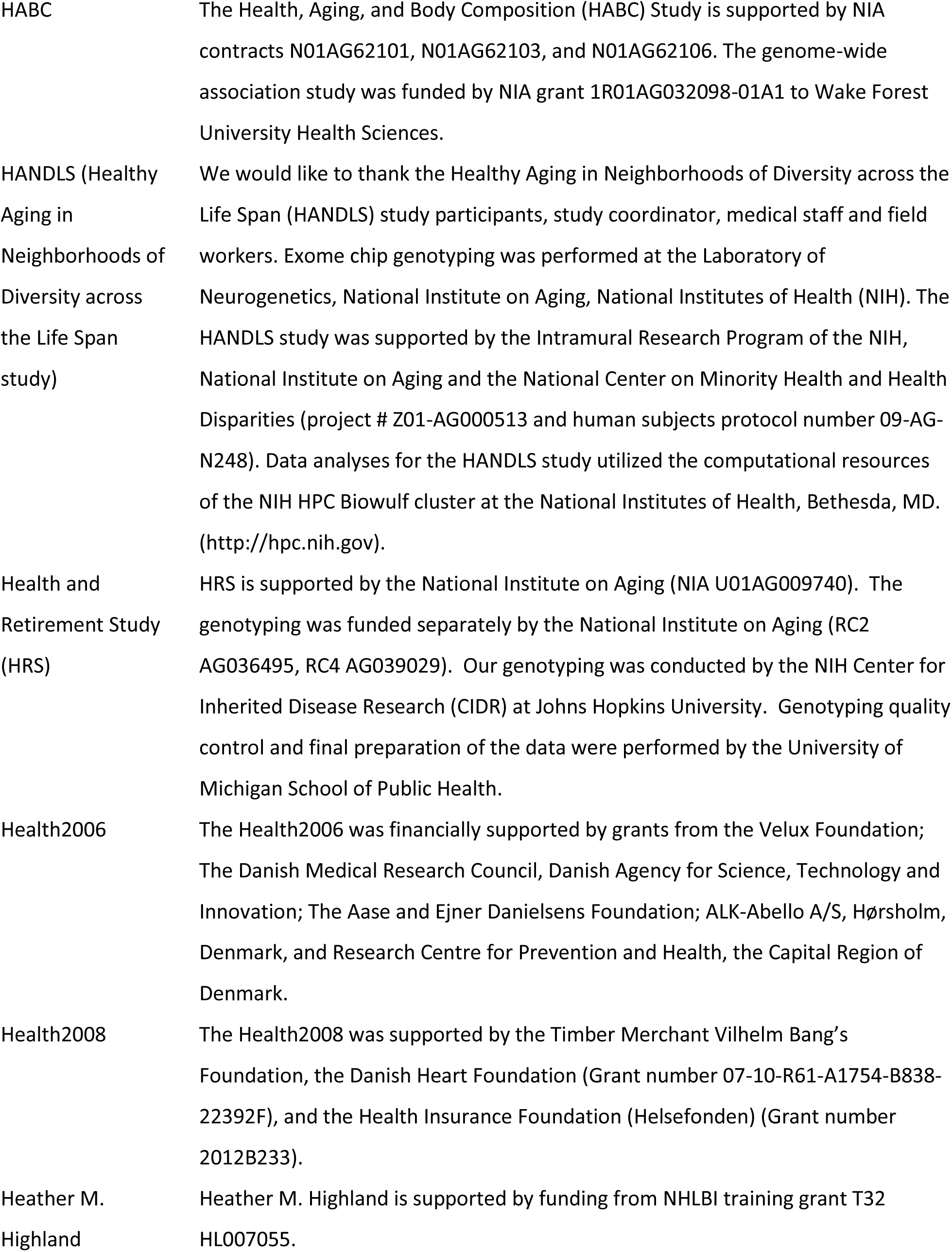
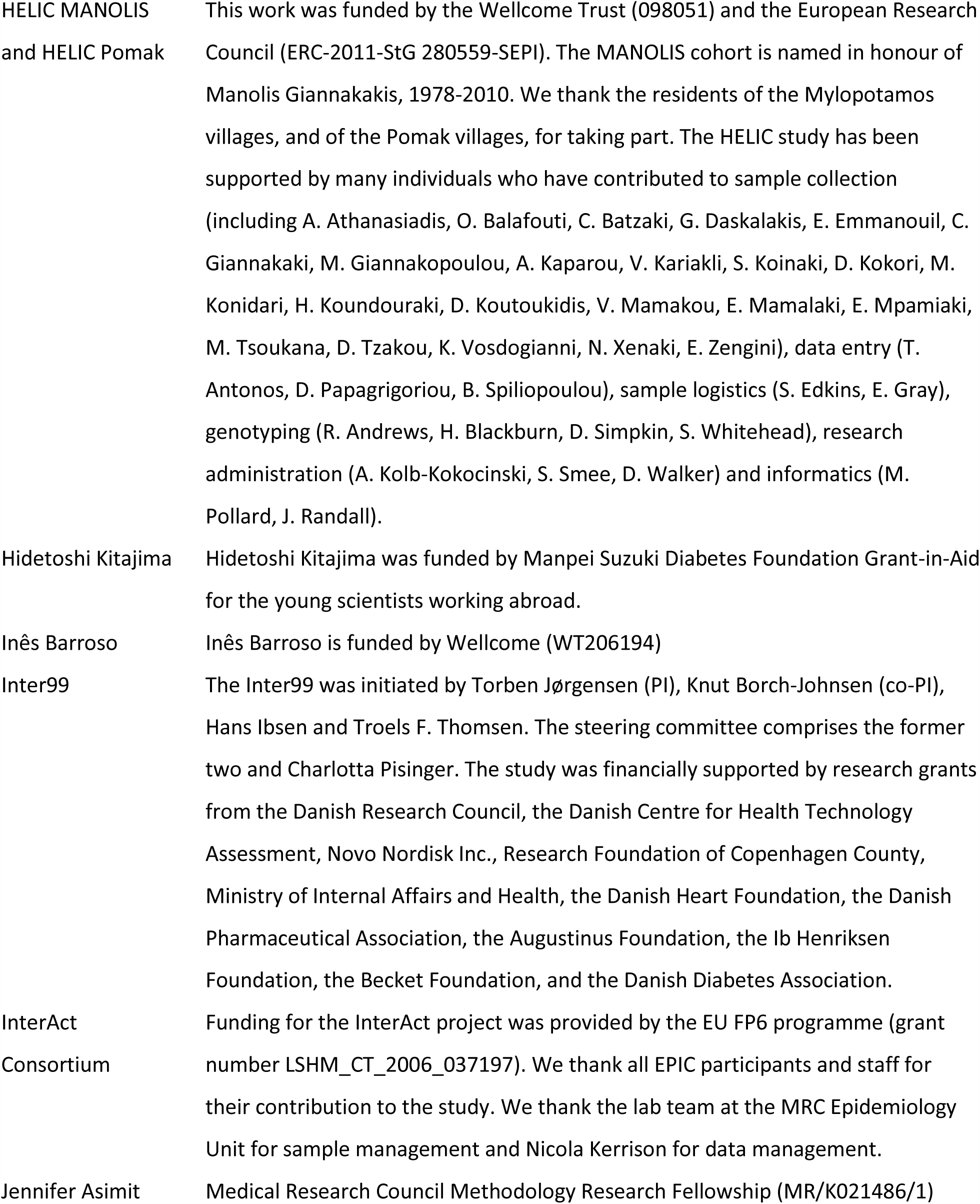
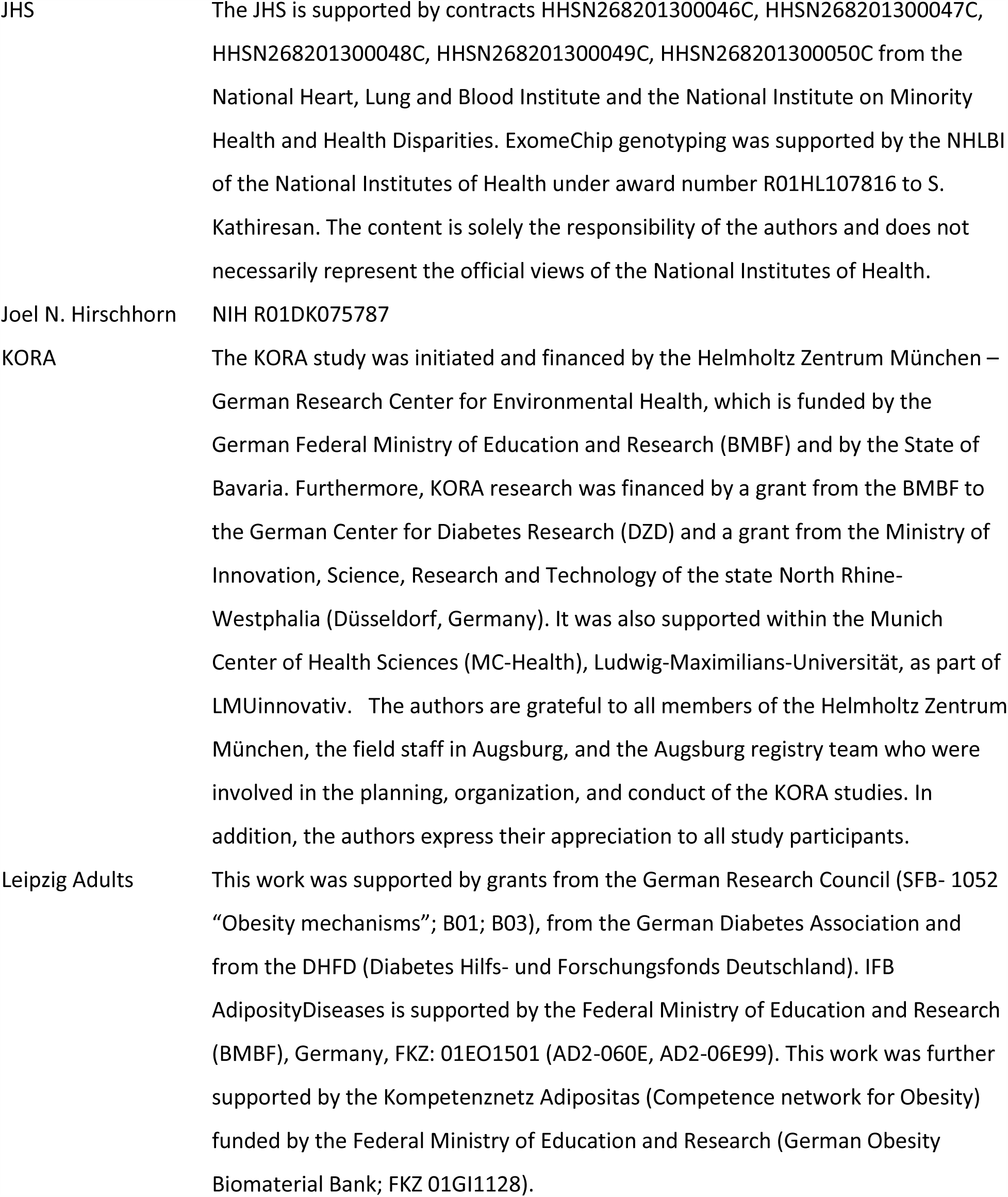
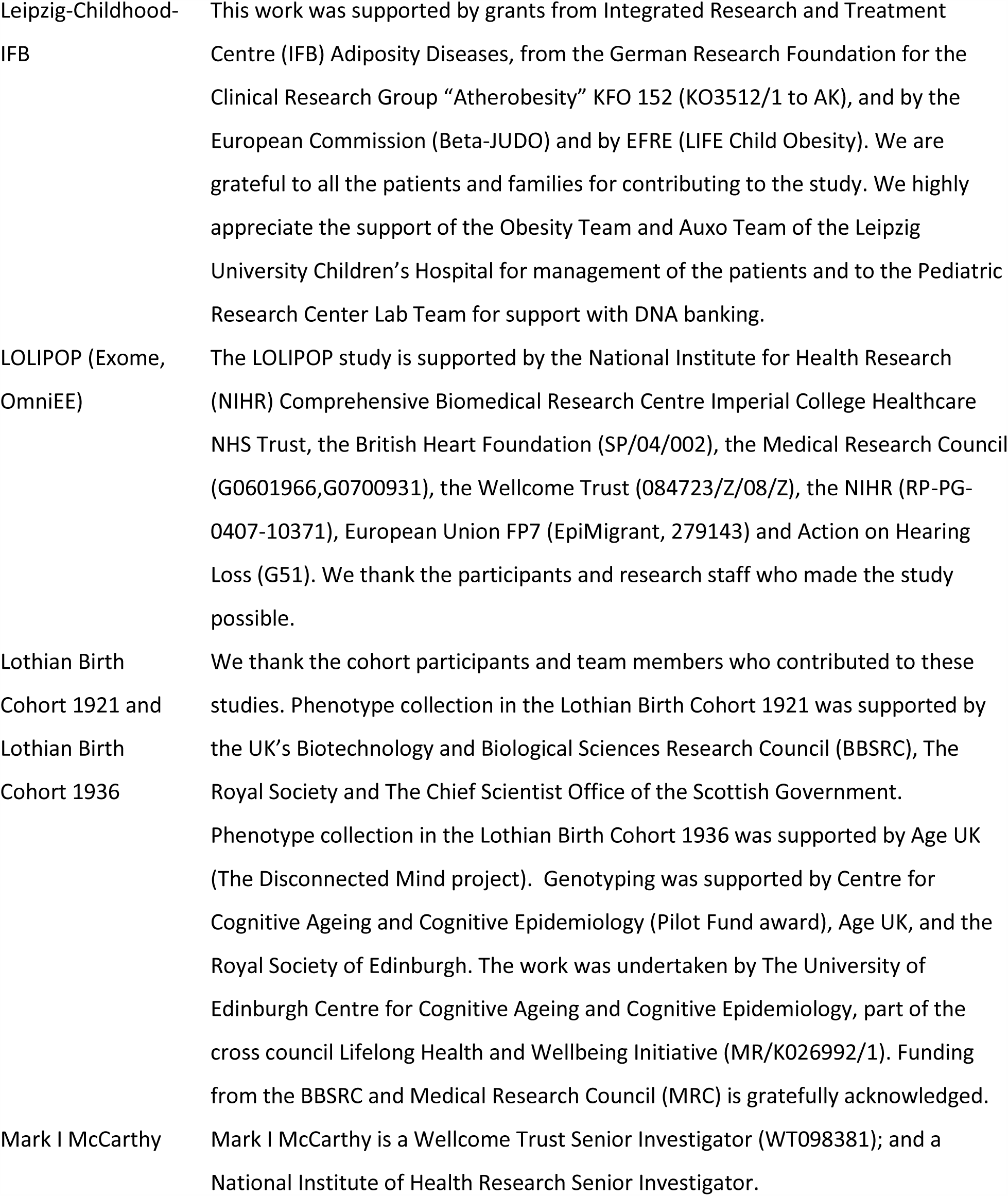
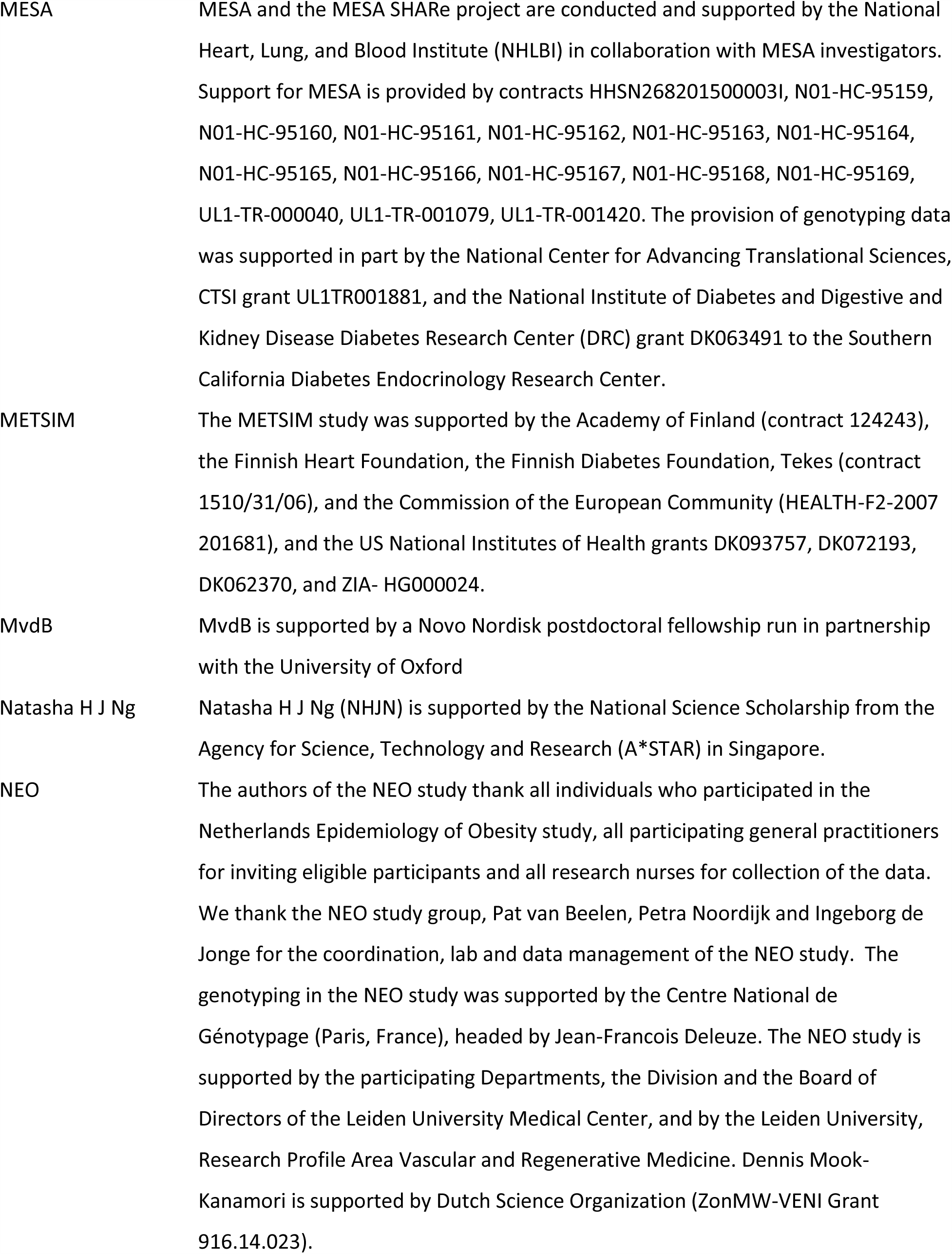
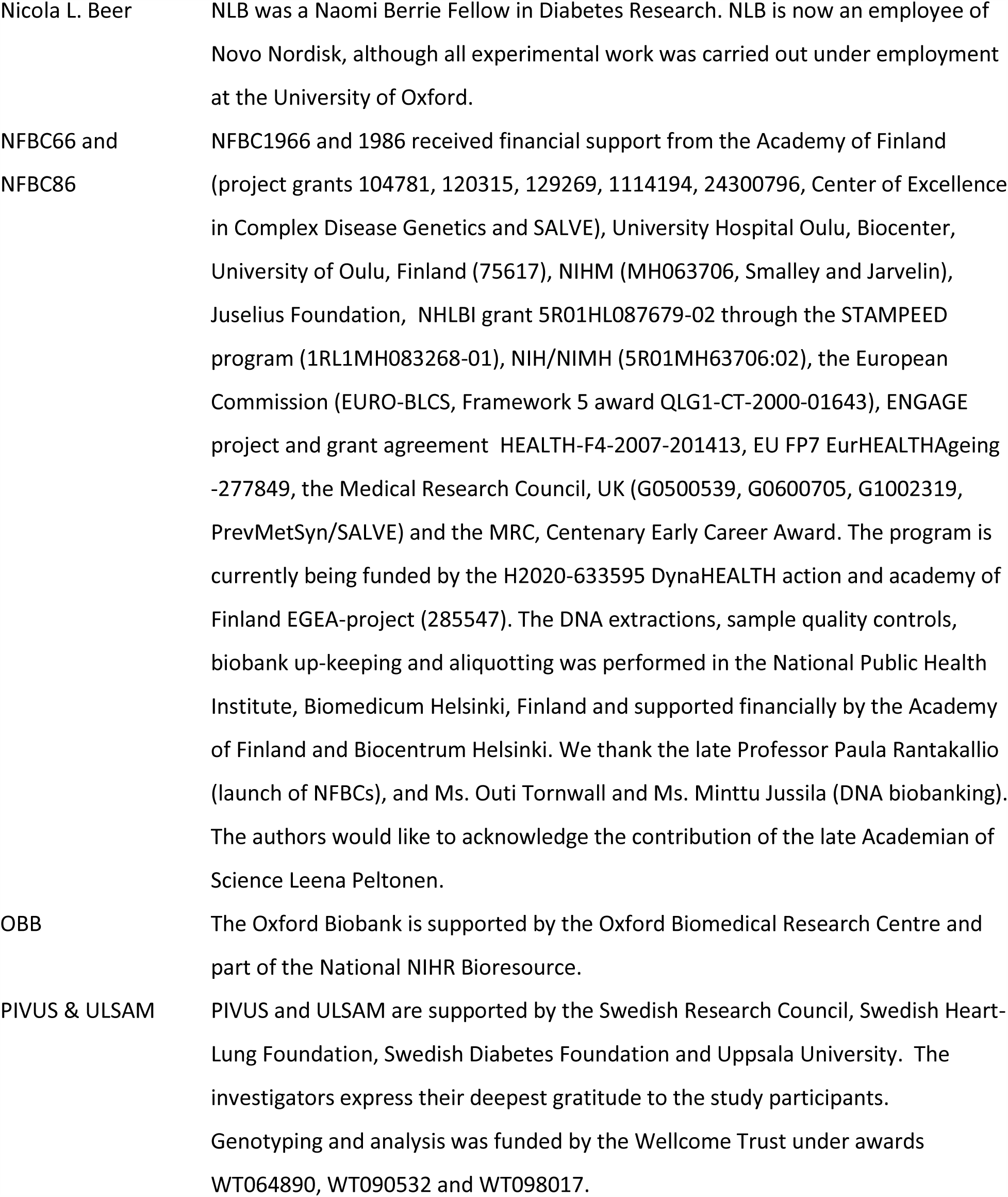
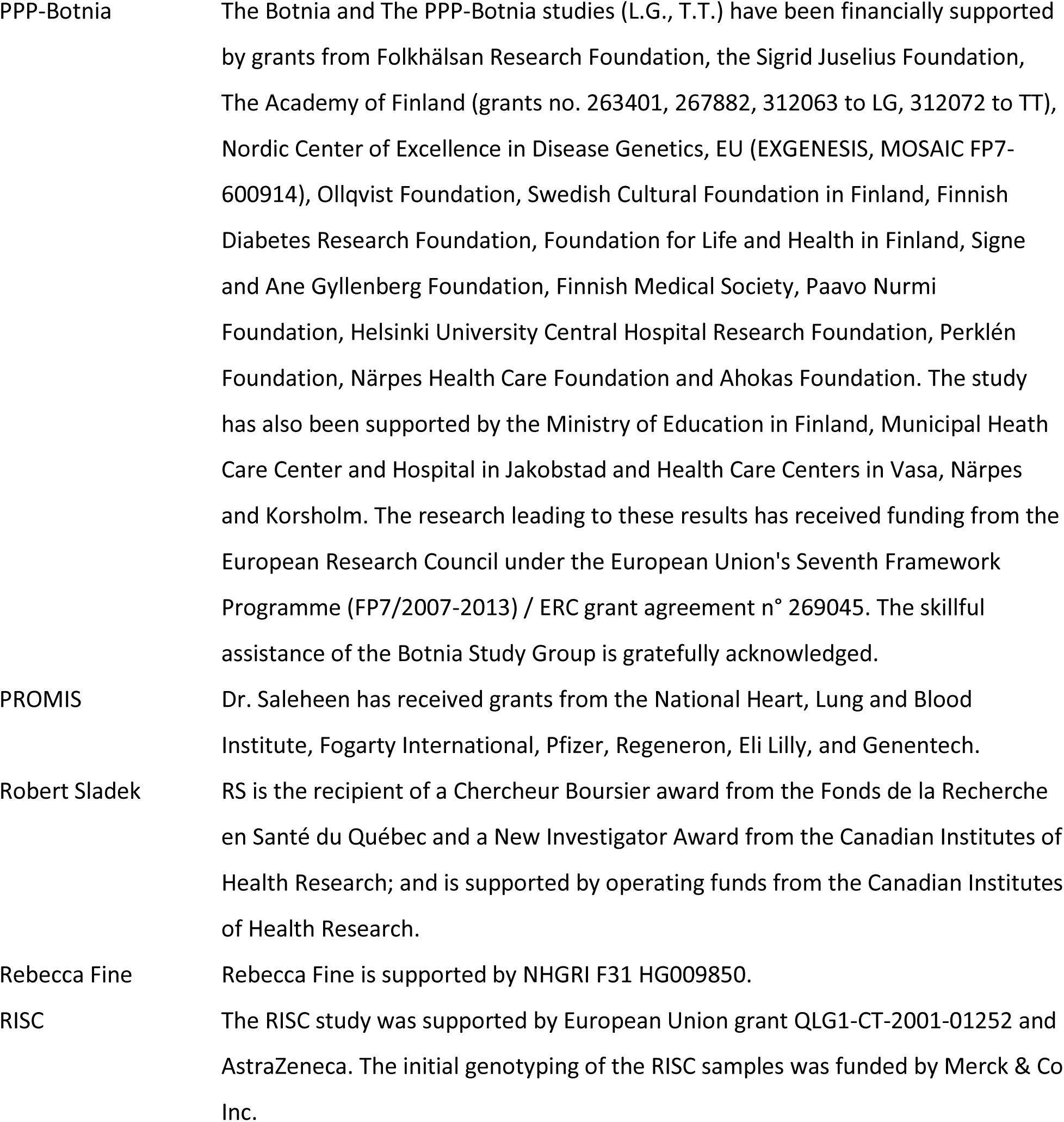
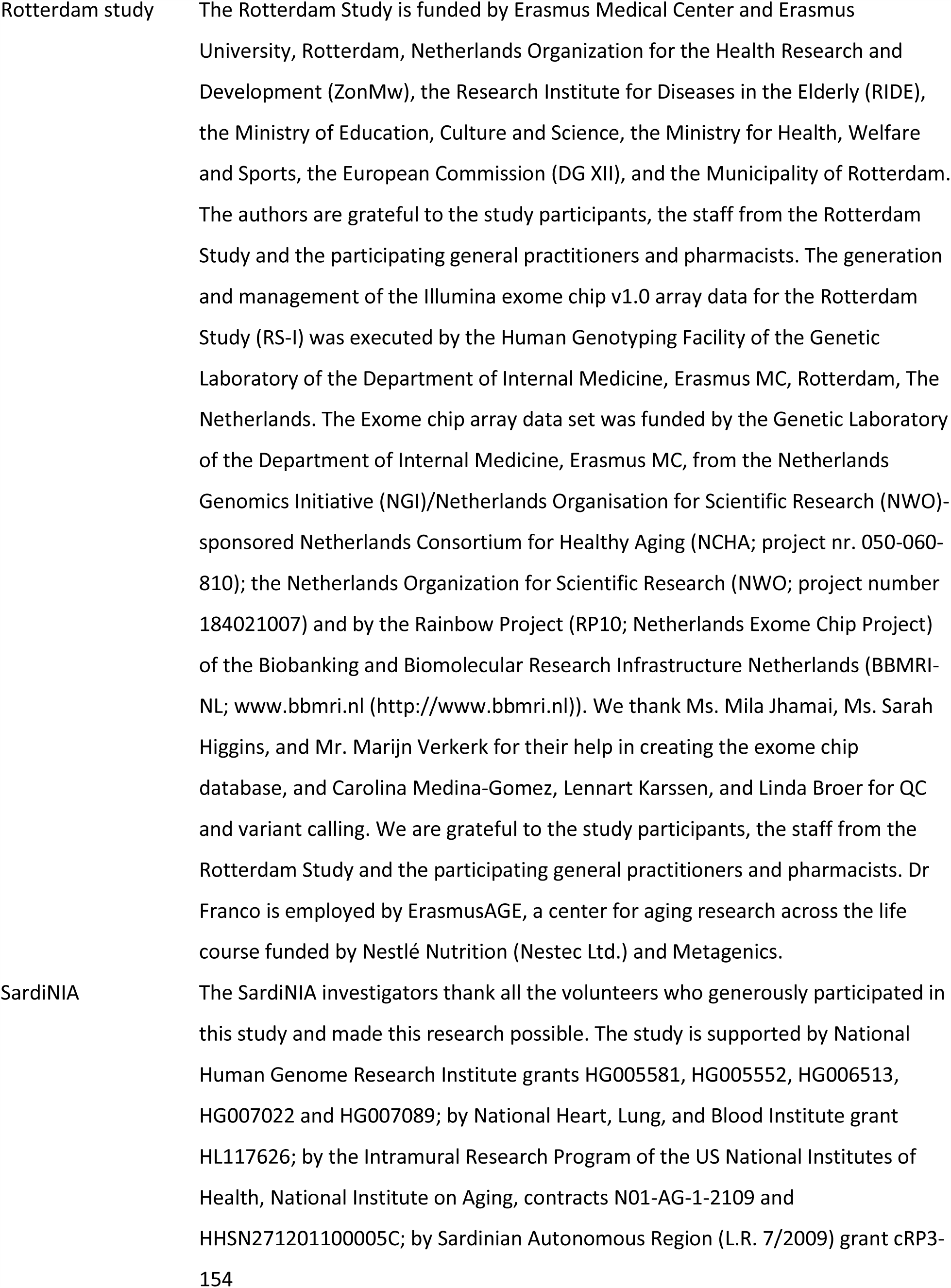
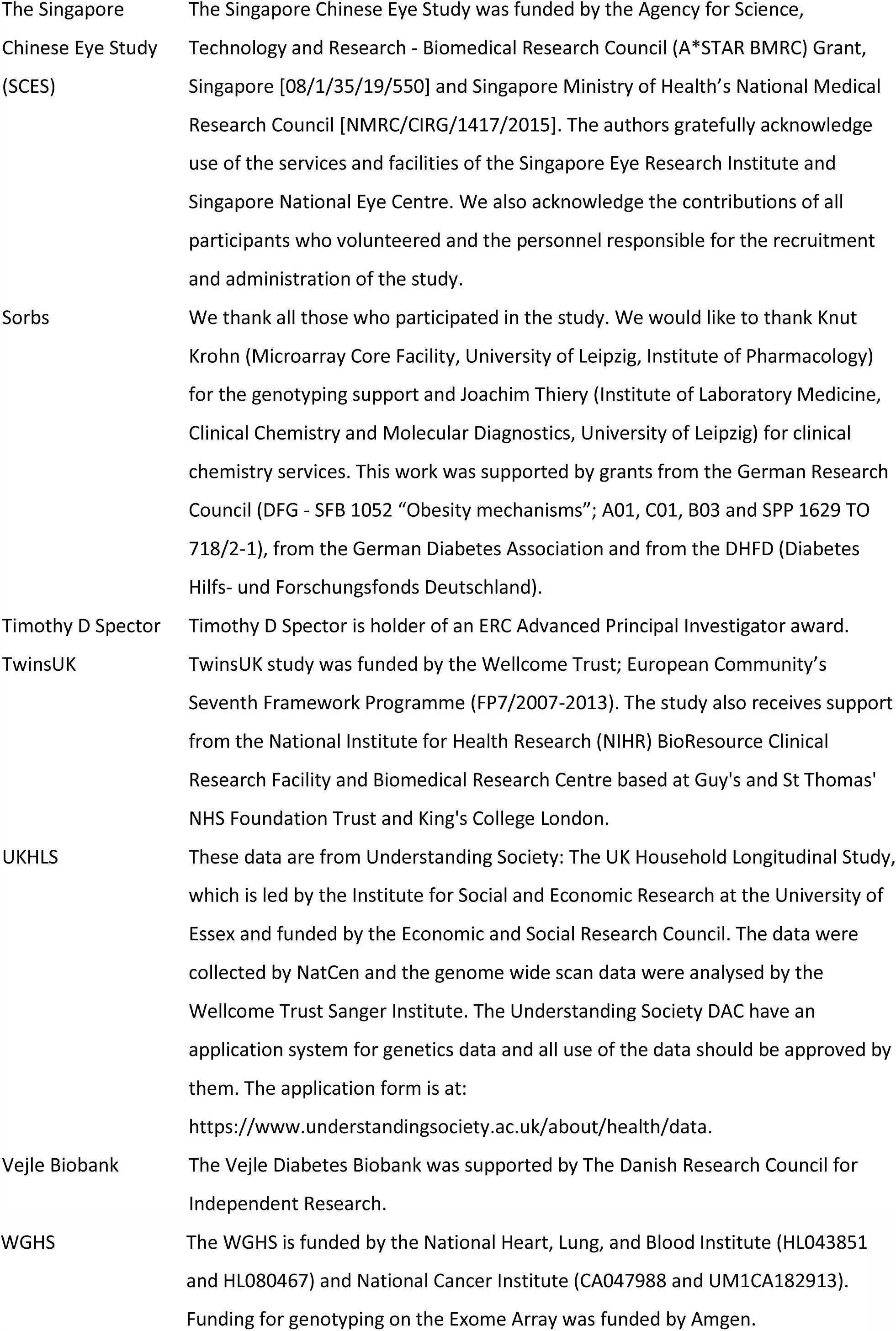

## Disclosures

Soren K. Thomsen: SKT is now an employee of Vertex Pharmaceuticals, although all experimental work was carried out under employment at the University of Oxford.

Martijn van de Bunt: Currently employed by Novo Nordisk. Audrey Y Chu: Currently employed by Merck.

Dennis O. Mook-Kanamori: Dennis Mook-Kanamori is working as a part-time clinical research consultant for Metabolon, Inc.

Paul W. Franks: PWF has been a paid consultant for Eli Lilly and Sanofi Aventis and has received research support from several pharmaceutical companies as part of a European Union Innovative Medicines Initiative (IMI) project.

Mike A. Nalls: Dr. Mike A. Nalls is supported by a consulting contract between Data Tecnica International LLC and the National Institute on Aging (NIA), National Institutes of Health (NIH), Bethesda, MD, USA. Dr. Nalls also consults for Illumina Inc., the Michael J. Fox Foundation, and the University of California Healthcare.

Mark J. Caulfield: MJC is Chief Scientist for Genomics England, a UK government company.

Joel N. Hirschhorn: JHN is on the scientific advisory board of Camp4 Therapeutics.

Erik Ingelsson: Erik Ingelsson is an advisor and consultant for Precision Wellness, Inc., and advisor for Cellink.

Inês Barroso: IB and spouse declare stock ownership in GlaxoSmithkline and Incyte Ltd.

## Supplementary Figure Legends

**Figure S1. GeneMANIA network analysis identifies relevant pathways regulating glycemia.** The networks represent composite networks for (A) FI and (B) 2hGlu, from the GeneMANIA analysis using genes with variant associations at *P*<1 × 10^−5^ for each trait as input. Nodes outlined in red correspond to genes from the input list. Other nodes correspond to related genes based on 50 default databases. Based on the network, GO terms and Reactome pathways that were significantly enriched are depicted. To summarize these results, the most significant term of all calculated terms within the same group (using the Kappa method, see **Methods**) was represented. Each group was assigned a specific color; if a gene is present in more than one term, it will be displayed in more than one color. Barplots with the Bonferroni-adjusted -log10(p-values) of the most significant terms within each group are are shown. Each group was assigned a specific color; if a gene is present in more than one term, it is displayed in more than one color. Details of the networks are summarized in (C). Related to Figure 2 and Table S7.

**Figure S2. Pathway analysis identifies relevant gene sets regulating glycemia.** EC-DEPICT analysis with heatmap visualization (UK Biobank permutations) is shown for a. all traits combined; b. HbA1c only; c. all traits except HbA1c combined; d. FG only; e. 2hGlu only. The columns represent the input genes for the analysis. We used affinity propagation clustering to define a representative “meta-gene set” for groups of highly correlated gene sets (see **Methods**); the rows in the heat map represent significant meta-gene sets (FDR <0.05). The color of each square indicates DEPICT’s z-score for membership of that gene in that gene set, where dark red means “very likely a member” and dark blue means “very unlikely a member”. The gene set annotations indicate whether that meta-gene set was significant at FDR <0.05 or not significant (n.s.) for each of the other EC-DEPICT analyses using the UK Biobank permutations (all traits together, HbA1c only, FG only, 2hGlu only, and all-except-HbA1c). For heatmap intensity and EC-DEPICT *P*-values, the meta-gene set values are taken from the most significantly enriched member gene set. The gene variant annotations are as follows: (1) the European minor allele frequency (MAF) of the input variant, where rare is MAF <1%, low-frequency is MAF 1-5%, and common is MAF > 5%, (2) whether the gene has an Online Mendelian Inheritance in Man (OMIM) annotation as causal for a diabetes/glycemic-relevant syndrome or blood disorder, (3) the effector transcript classification for that variant: gold, silver, bronze, or NA (note that only array-wide significant variants were classified, so suggestively-significant variants are by default classified as “NA”), (4-7) whether each variant was significant (*P*<2 × 10^−7^), suggestively significant (*P*<10^−5^), or not significant in Europeans for each of the four traits, and (8) whether each variant was classified in the analysis (with UK Biobank permutations) or excluded by filters (see **Methods**). AWS: array-wide significant. Related to Figure 2 and Table S8.

**Figure S3. Pathway analysis identifies relevant gene sets regulating glycemia.** EC-DEPICT analysis with heatmap visualization (Swedish permutations) is shown for a. all traits combined; b. HbA1c only; c. all traits except HbA1c combined; d. FG only. (With these permutations, there was no significance for 2hGlu only). We used affinity propagation clustering to define a representative “meta-gene set” for groups of highly-correlated gene sets (see **Methods**); the rows in the heat map represent significant meta-gene sets (FDR <0.05). The color of each square indicates DEPICT’s z-score for membership of that gene in that gene set, where dark red means “very likely a member” and dark blue means “very unlikely a member”. The gene set annotations indicate whether that meta-gene set was significant at FDR <0.05 or not significant (n.s.) for each of the other EC-DEPICT analyses using the Swedish permutations (all traits together, HbA1c only, FG only, and all-except-HbA1c). For heatmap intensity and EC-DEPICT *P*-values, the meta-gene set values are taken from the most significantly enriched member gene set. The gene variant annotations are as follows: (1) the European minor allele frequency (MAF) of the input variant, where rare is MAF <1%, low-frequency is MAF 1-5%, and common is MAF >5%, (2) whether the gene has an Online Mendelian Inheritance in Man (OMIM) annotation as causal for a diabetes/glycemic-relevant syndrome or blood disorder, (3) the effector transcript classification for that variant: gold, silver, bronze, or NA (note that only array-wide significant variants were classified, so suggestively-significant variants are by default classified as “NA”), (4-7) whether each variant was significant (*P*<2 × 10^−7^), suggestively significant (*P*<10^−5^), or not significant in Europeans for each of the four traits, and (8) whether each variant was included in the analysis (with Swedish permutations) or excluded by filters (see **Methods**). AWS: array-wide significant. Related to Figure 2 and Table S8.

**Figure S4. Functional characterisation of G6PC variants.** Related to Figure 4.

(A) Cellular localisation of Q347X was assessed in HEK293 cells and overlaid with a marker for the ER, calreticulin, (left) or the trans-golgi network, TGN46 (right). White arrows point to positions of the golgi apparatus. Scale bar indicates 10μm. (B) Glucose-6-phosphatase activity of G6PC-R83C (n=3), with representative western blot of microsomal protein isolated from HEK293 shown. (C) Glucose-6-phosphatase activity of G6PC-Q347X (n=2), with representative western blot of microsomal protein isolated from HEK293 shown. (D) Protein expression levels of G6PC-A204S in microsomal protein extracted from HEK293 cells was found to be downregulated by 41% compared to WT based on densitometric analysis (n=4), with representative western blot shown. Data presented as mean ± SEM and analysed using paired Students’ t test. * p=0.01. Unt: Untransfected; WT: Wild type.

**Figure S5. Functional characterisation of G6PC2 variants and the effect of G6PC2 knockdown on insulin content and secretion in EndoC-βH1 cells.** Related to Figure 5.

(A) Variants prioritised for functional study in the context of the predicted G6PC2 protein structure (RefSeq NP_066999.1) in the ER membrane. Amino acid residues are coloured as described in the legend. Variants selected for functional study, in green, are labelled. The N-terminal V5 and C-terminal Myc-FLAG tags present in the expression constructs are indicated. (B) Quantification of total G6PC2 variant protein expression (both upper and lower bands of representative western blot in Figure 5) in INS-1 832/13 cells based on western blot densitometric analysis of Myc-tagged G6PC2 constructs relative to tubulin control (n=5). (C) Expression levels of G6PC2 variant proteins in HEK293 by western blot densitometric analysis of FLAG-tagged G6PC2 constructs or V5-tagged G6PC2-R283X relative to tubulin control (n=4). Representative blots are shown for untreated cells and cells treated with proteasomal inhibitor MG-132 or lysosomal inhibitor chloroquine. (D) Glucose-6-phosphatase activity of the R281X variant in G6PC (proxy for R283X in G6PC2) in HEK293 (n=2), with representative western blot of microsomal protein shown. (E) Total insulin secretion and insulin content were assessed at basal and high glucose conditions (with and without drug treatment) following 96-120h *G6PC2* knockdown in EndoC-βH1. Unpaired two-tailed Students’ t tests were used to compare *G6PC2* knockdown to control for each condition, from n=16 across 4 independent experiments. Tol: tolbutamide; Diaz: diazoxide. All data presented as mean ± SEM. * p=0.01-0.05; ** p=0.001-0.01; *** p<0.001.

**Figure S6. *G6PC2* expression in RNA-Seq data from 150 human islet donor samples.** (A) Allelic balance was observed for *G6PC2* rs146779637 (p.R283X) in two heterozygote human islet samples. (B) The glucose-raising rs560887-G allele associates significantly (*q*-value<0.01) with increased expression of the long *G6PC2* isoform (purple) and reduced expression of the short *G6PC2* isoform lacking exon 4 (brown).

## Supplementary Table Legends

**Table S1. Cohort characteristics, genotyping and quality control (QC), glucose, insulin, 2hGlu and HbA1c analyses and covariates.**

**Table S2. Association of identified lead coding variants with T2D and anthropometric traits (height, BMI and WHR) from publicly-available association results.** Alleles E/O: effect allele/other allele; EAF: effect allele frequency; Neff: Number of samples in the analysis; BETA: effect size; SE: standard error. Related to Table 1 and Table S3.

**Table S3. Coding variant associations in known glycemic trait loci with conditional results on established signals where available.** Related to Table 1.

**Table S4. Full gene-based results including all variants included in the masks, for both novel and previously-established genes.** Related to Table S9.

**Table S5. HbA1c-associated loci lookup results for blood cell traits.** Related to Table 1.

**Table S6. Annotation and classification of effector transcripts into “gold”, “silver” and “bronze” categories.** Related to Tables 1 and 2 and Figure 1.

**Table S7. Gene Set Enrichment Analysis by GeneMANIA network analysis showing enriched GO terms and Reactome pathways in the network for (A) HbA1c; (B) FG; (C) FI; (D) 2hGlu.** GOID: Gene Ontology ID; GOTerm: Gene Ontology Term. Gene Set Enrichment (GSE) of networks was performed with ClueGO using GO terms and REACTOME gene sets. The enrichment results were considered significant when Bonferroni-adjusted p-value < 0.05 and at least 3% of the genes contained in the tested gene set is included in the network. Gene sets were also grouped using kappa score into functional groups to improve visualization of enriched pathways. Related to Figures 2 and S1.

**Table S8. (A-E) EC-DEPICT results (UK Biobank permutations) for (A) all traits combined; (B) all traits except HbA1c combined; (C) HbA1c only; (D) FG only and (E) 2hGlu only. (F-I) EC-DEPICT results (Swedish permutations) for (F) all traits combined; (G) all traits except HbA1c combined; (H) HbA1c only and (I) FG only.** Related to Figures 2, S2 and S3.

**Table S9. Full *G6PC2* gene-based results and conditional analyses for FG and HbA1c.** Related to Tables 2 and S4.

