## Supplemental Information for "Tissue-Specific Alteration of Metabolic Pathways Influences Glycemic Regulation"

COLLABORATORS

Michaela Benzeval[1], Jonathan Burton[1], Annette Jäckle[1], Meena Kumari[1], Heather Laurie[1], Peter Lynn[1], Stephen Pudney[1], Birgitta Rabe[1], Dieter Wolke[2], Nicholas Buck[1]

1. Institute for Social and Economic Research
2. University of Warwick

Figure S1

A

**FI**

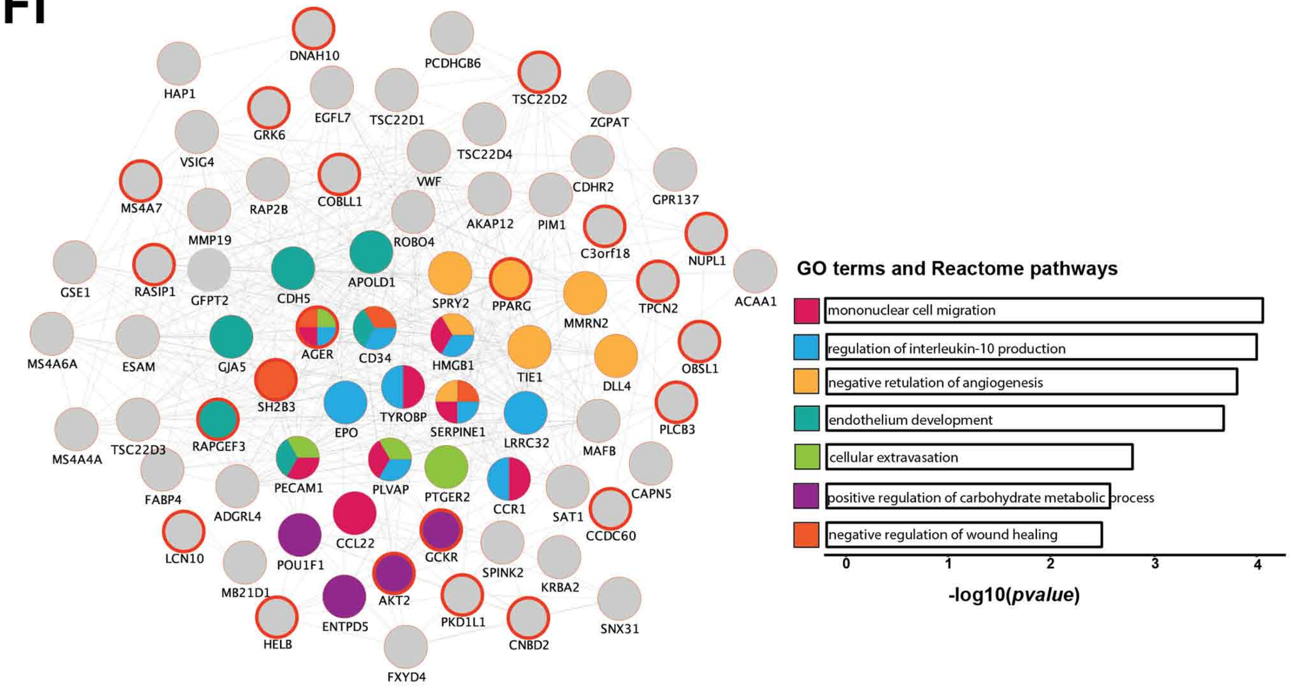


C

| Trait | # Genes in query list | # Nodes | # Edges | Clustering coefficient |
|---|---|---|---|---|
| HbA1c | 57 | 105 | 1284 | 0.52 |
| FG | 28 | 76 | 173 | 0.26 |
| FI | 24 | 74 | 424 | 0.33 |
| 2hGlu | 12 | 60 | 78 | 0.12 |

B

**2hGlu**

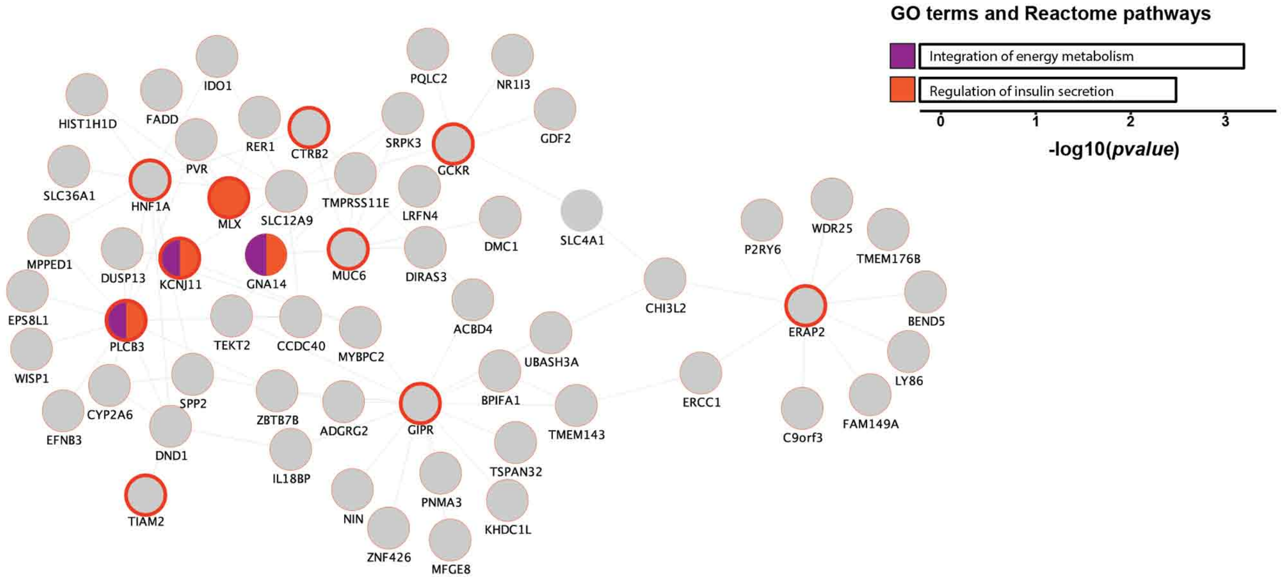


**Figure S1. GeneMANIA network analysis identifies relevant pathways regulating glycemia.** The networks represent composite networks for (A) FI and (B) 2hGlu, from the GeneMANIA analysis using genes with variant associations at P<1 × 10-5 for each trait as input. Nodes outlined in red correspond to genes from the input list. Other nodes correspond to related genes based on 50 default databases. Based on the network, GO terms and Reactome pathways that were significantly enriched are depicted. To summarize these results, the most significant term of all calculated terms within the same group (using the Kappa method, see Methods) was represented. Each group was assigned a specific color; if a gene is present in more than one term, it will be displayed in more than one color. Barplots with the Bonferroni-adjusted -log10(p-values) of the most significant terms within each group are are shown. Each group was assigned a specific color; if a gene is present in more than one term, it is displayed in more than one color. Details of the networks are summarized in (C). Related to Figure 2 and Table S7.

Figure S2
a. All traits combined

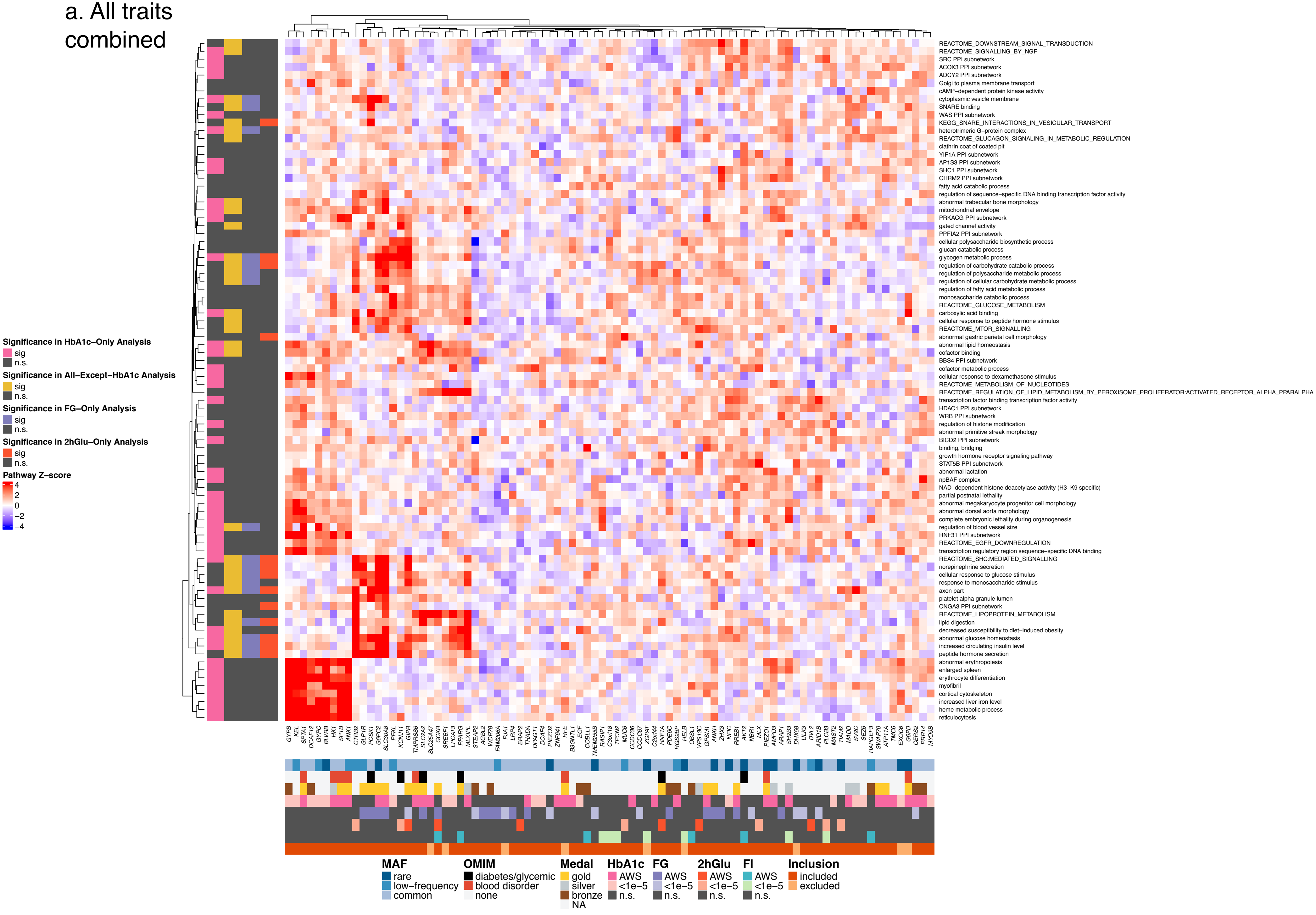

Figure S2
b. HbA1c only

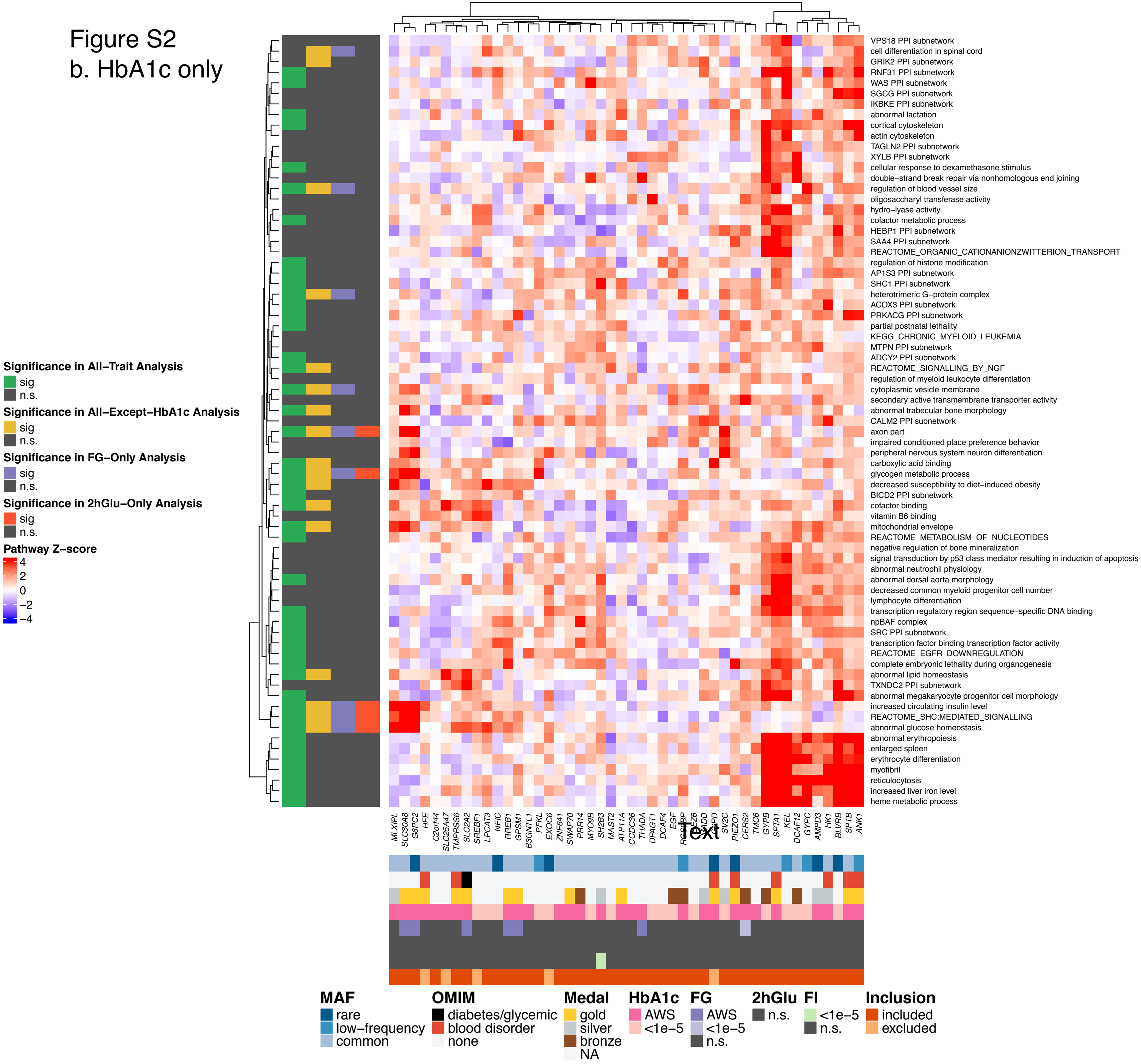

### Figure S2
#### c. All traits except HbA1c

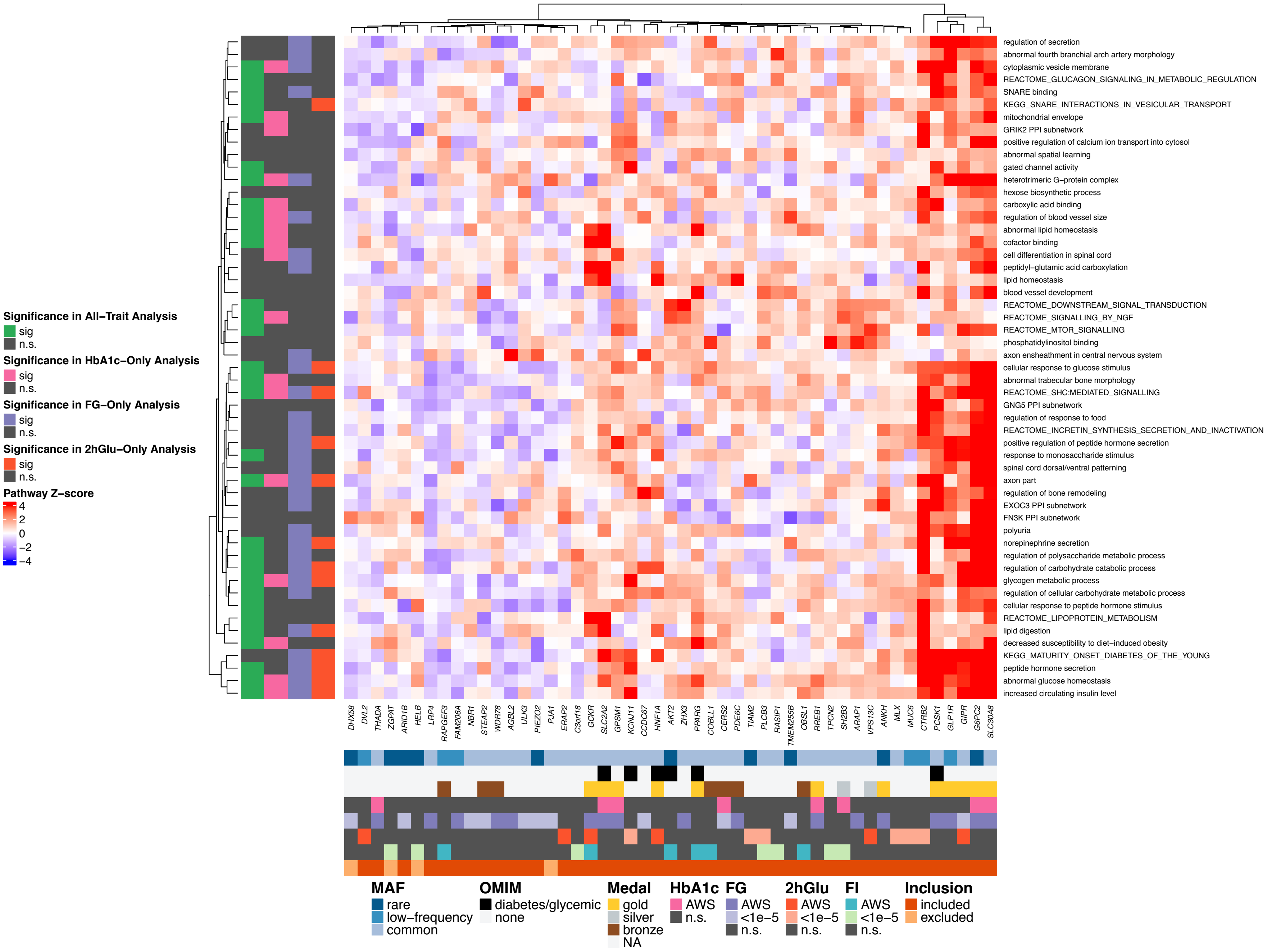

Figure S2
d. FG only

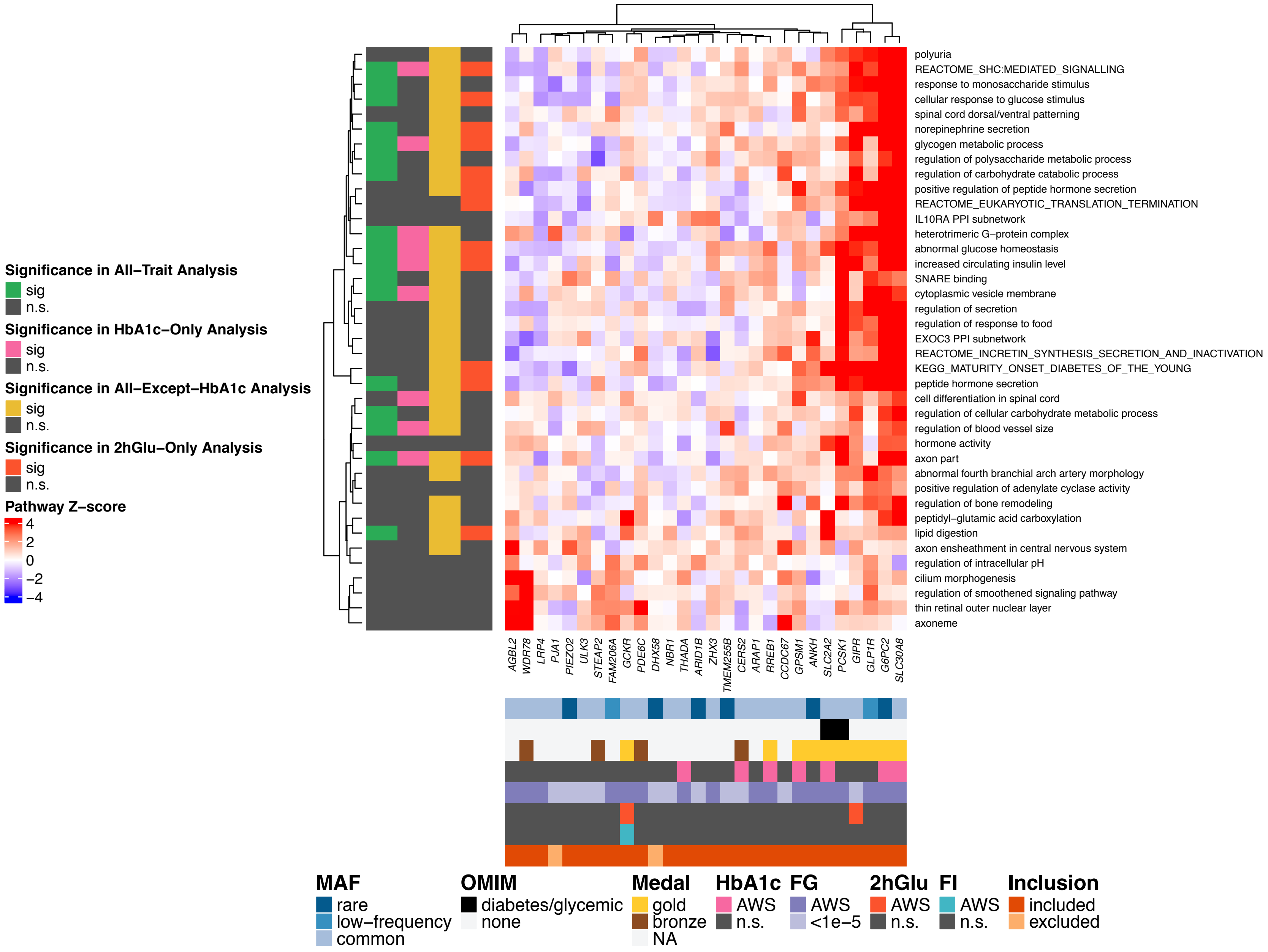

Figure S2
e. 2hGlu only

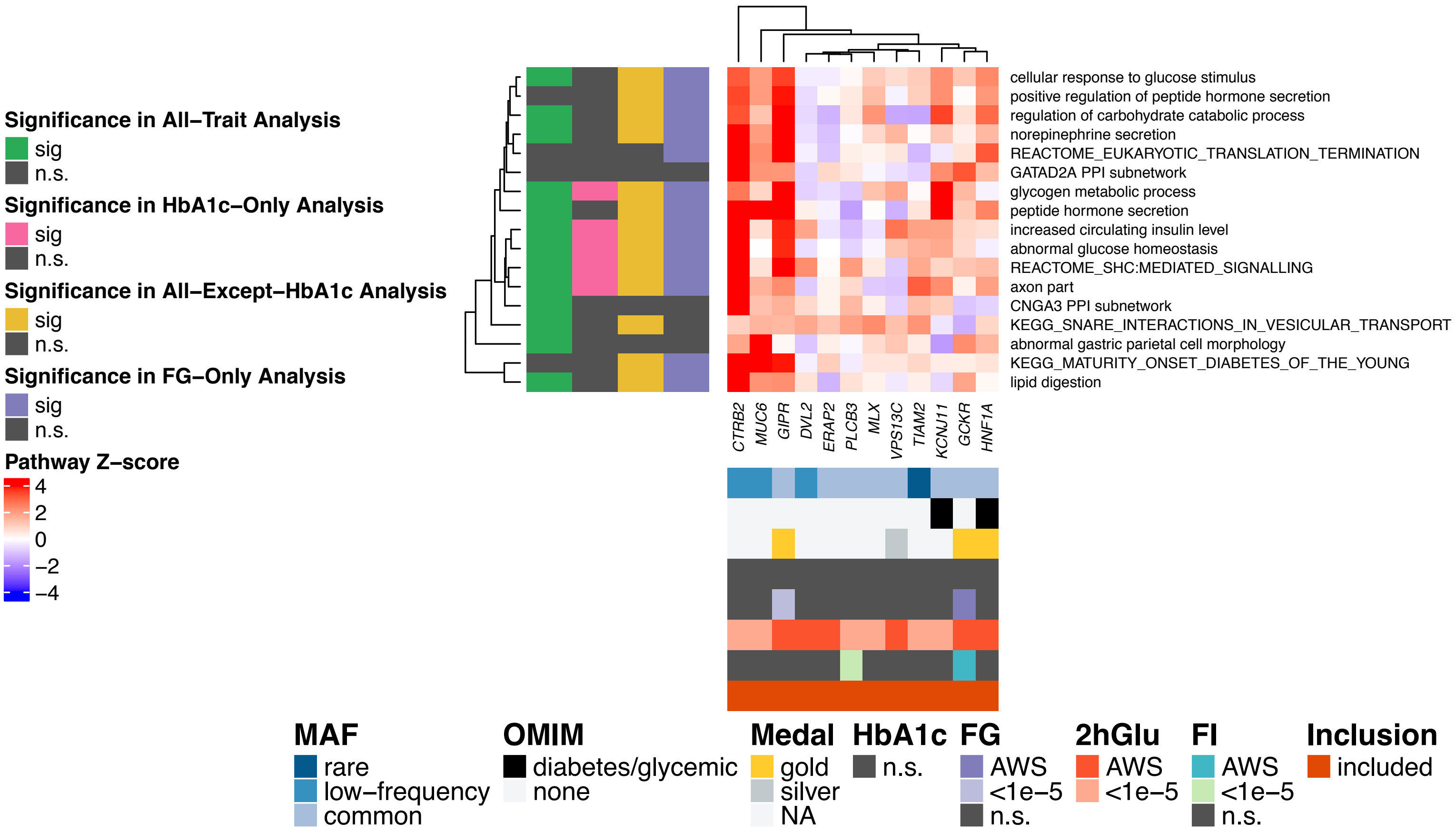

### Figure S3

#### a. All traits combined

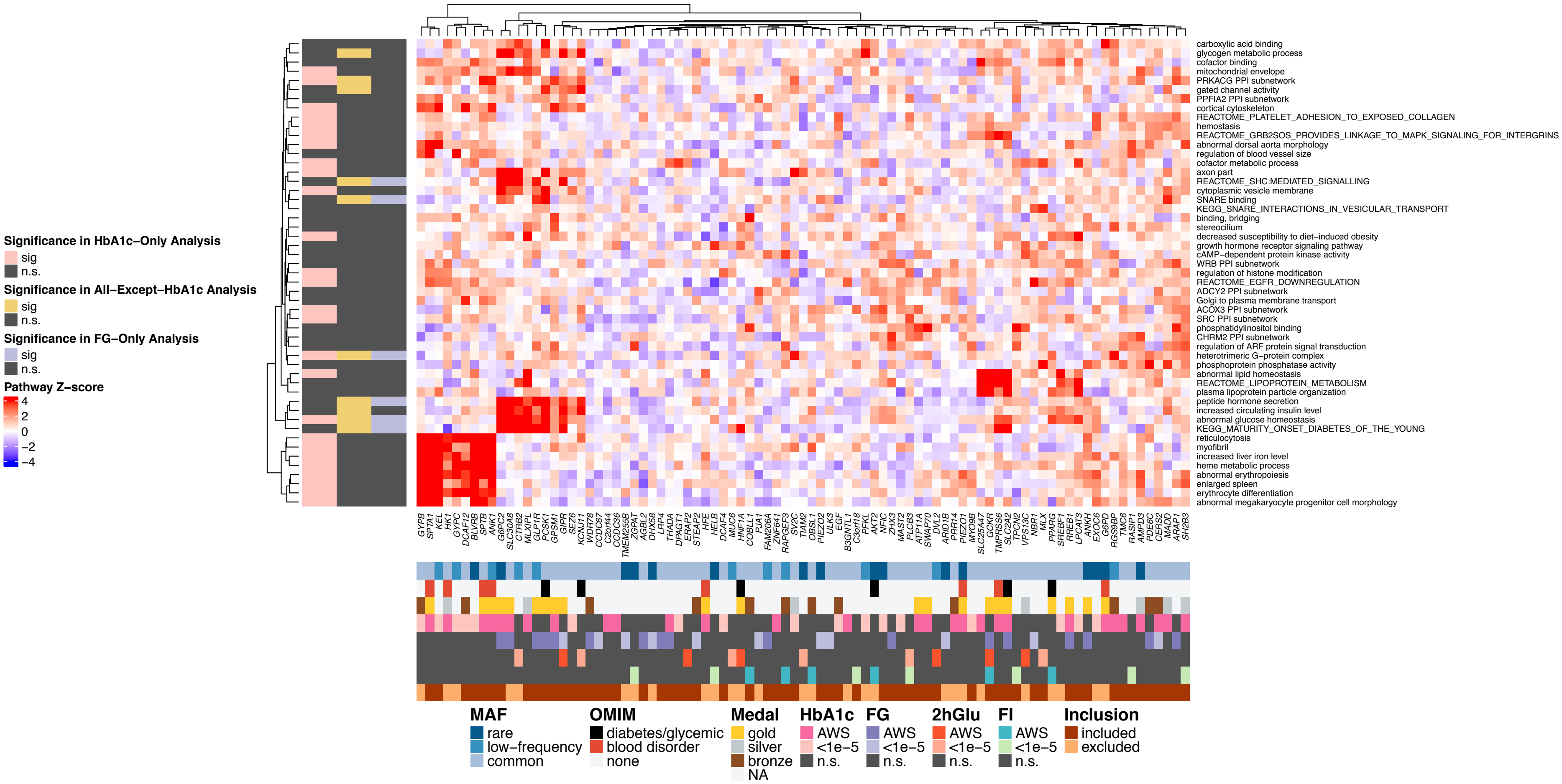

### Figure S3
#### b. HbA1c only

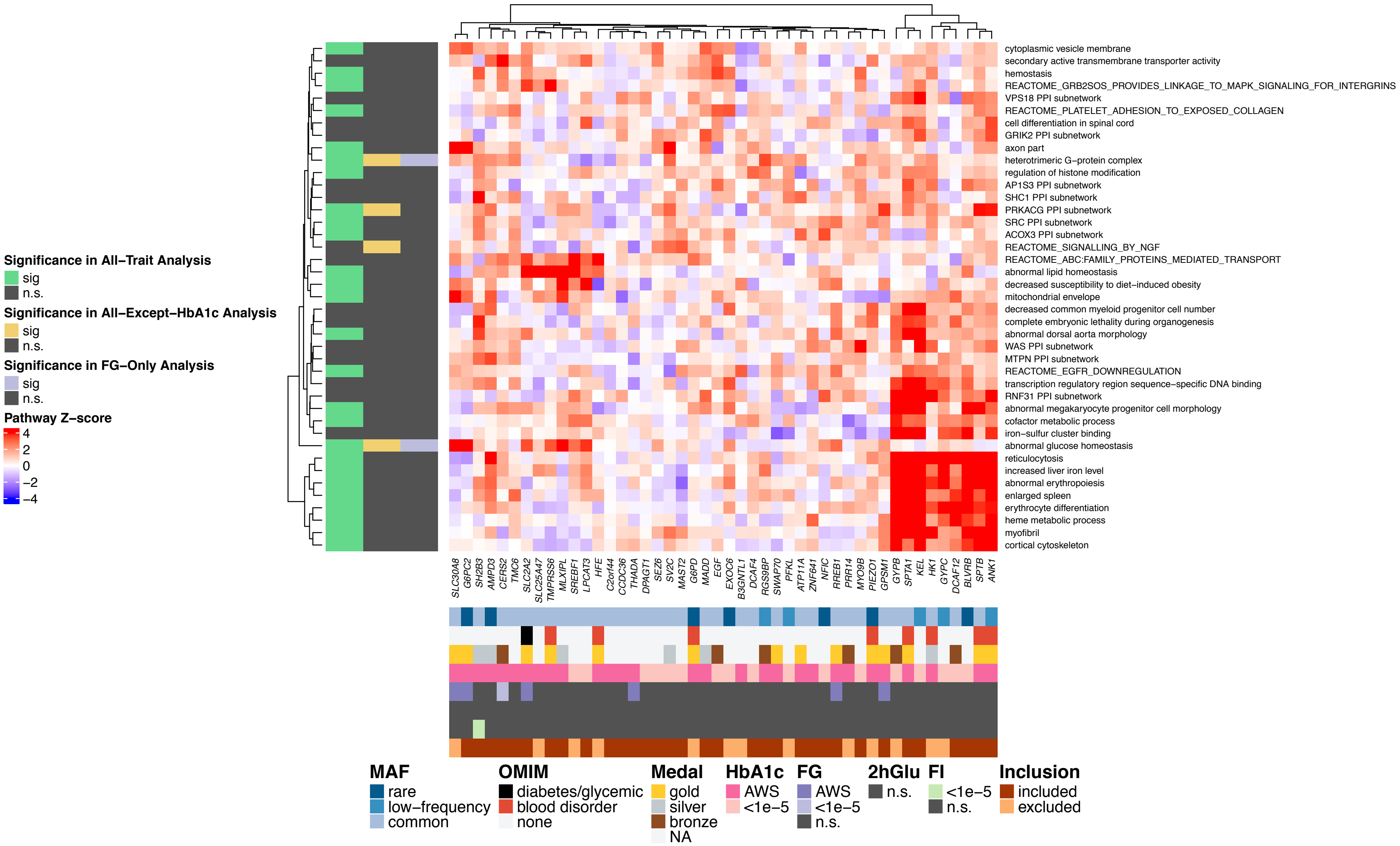

Figure S3

c. All traits except HbA1c

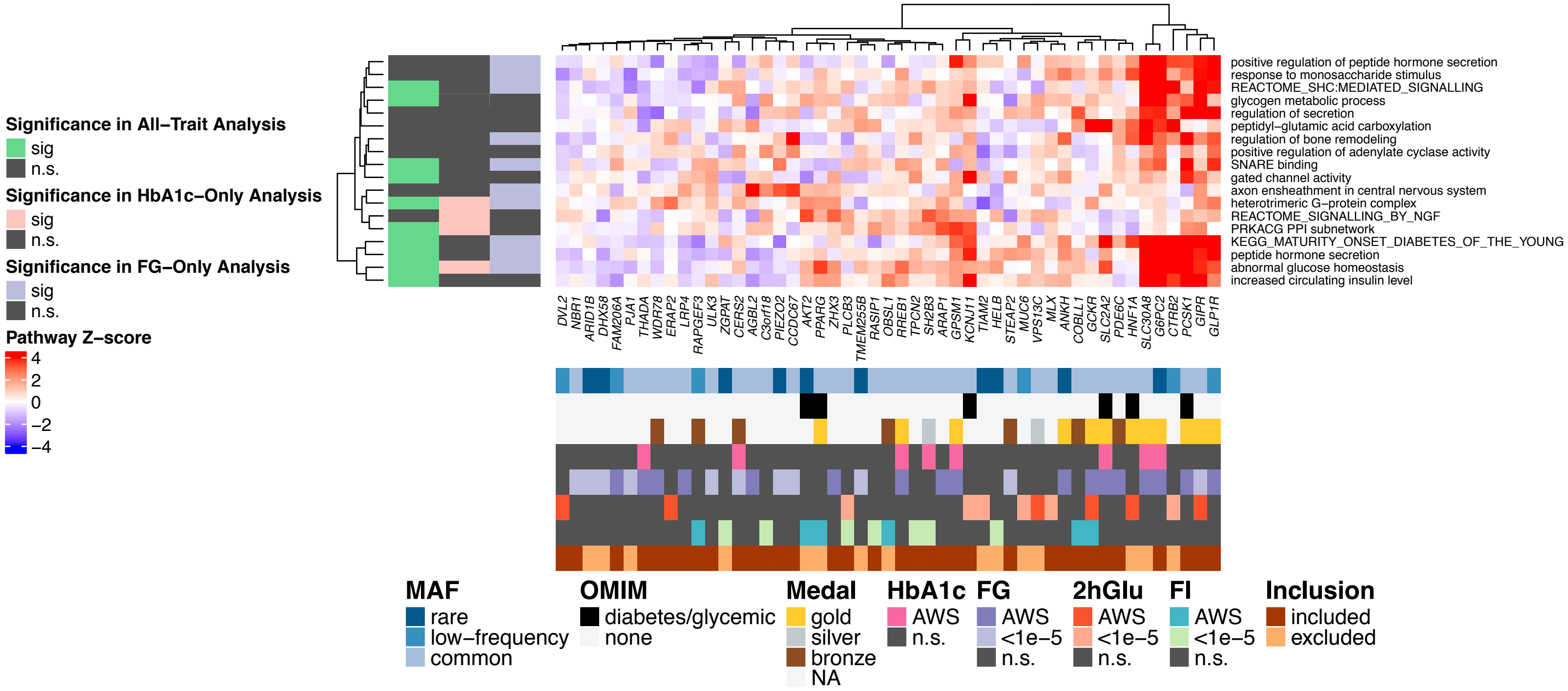

Figure S3
d. FG only

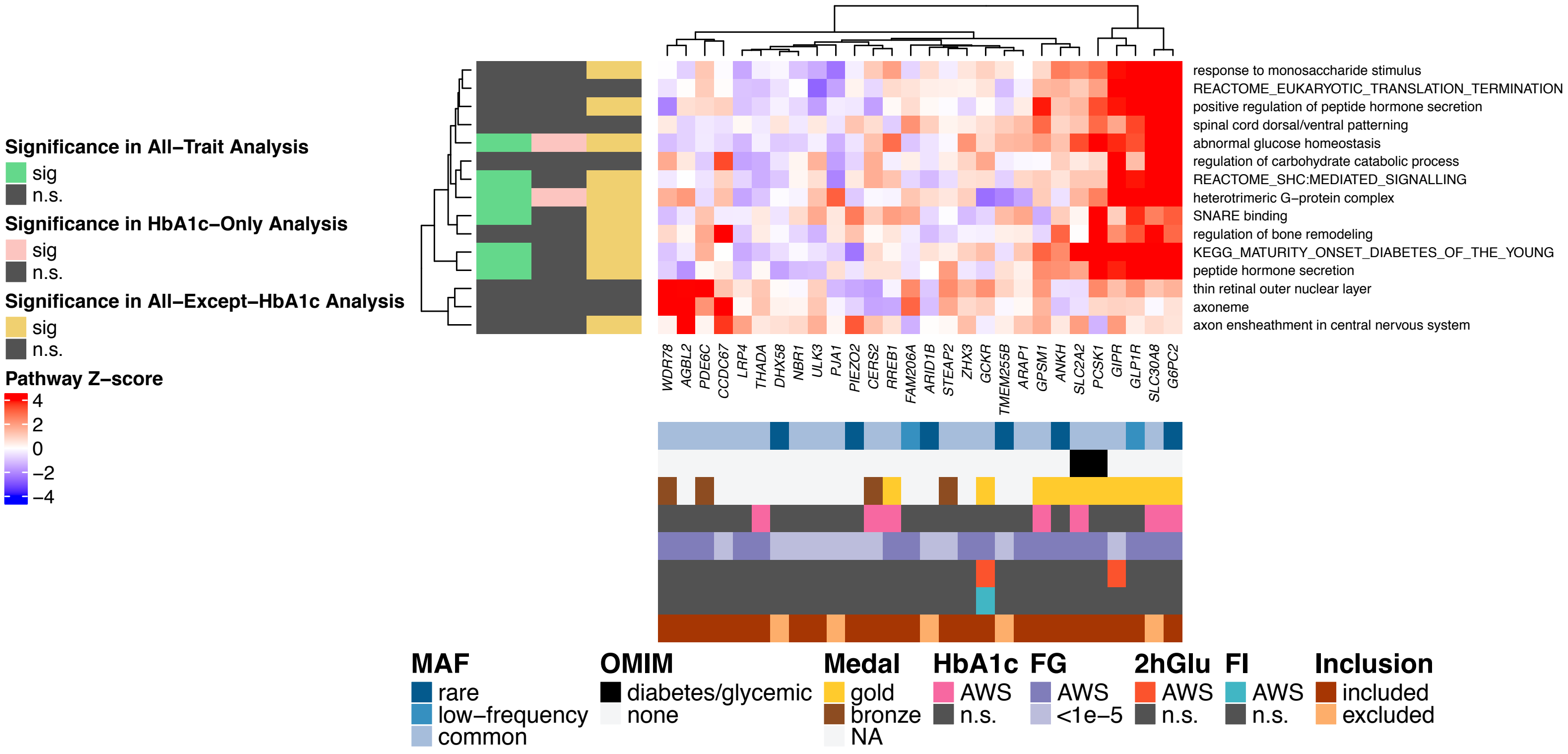

### Figure S4

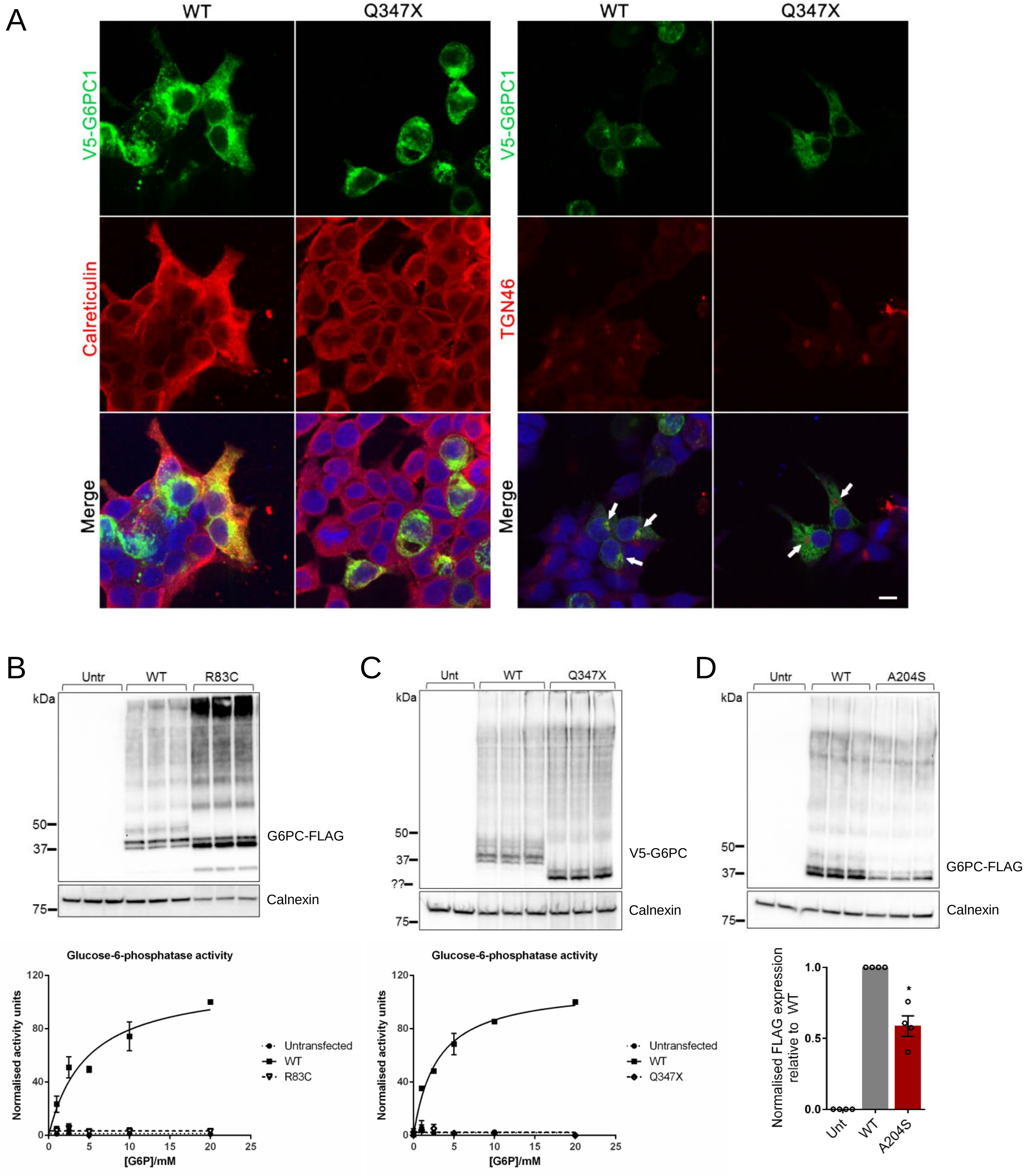

A
WT
Q347X
WT
Q347X
V5-G6PC1
Calreticulin
Merge
V5-G6PC1
TGN46
Merge
B
Untr
WT
R83C
kDa
50
37
75
G6PC-FLAG
Calnexin
C
Unt
WT
Q347X
kDa
50
37
??
75
V5-G6PC
Calnexin
D
Untr
WT
A204S
kDa
50
37
75
G6PC-FLAG
Calnexin
Glucose-6-phosphatase activity
Normalised activity units
[G6P]/mM
Untransfected
WT
R83C
Glucose-6-phosphatase activity
Normalised activity units
[G6P]/mM
Untransfected
WT
Q347X
Normalised FLAG expression relative to WT
Unt
WT
A204S

**Figure S4. Functional characterisation of G6PC variants.** Related to Figure 4.
(A) Cellular localisation of Q347X was assessed in HEK293 cells and overlaid with a marker for the ER, calreticulin, (left) or the trans-golgi network, TGN46 (right). White arrows point to positions of the golgi apparatus. Scale bar indicates 10μm.
(B) Glucose-6-phosphatase activity of G6PC-R83C (n=3), with representative western blot of microsomal protein isolated from HEK293 shown.
(C) Glucose-6-phosphatase activity of G6PC-Q347X (n=2), with representative western blot of microsomal protein isolated from HEK293 shown.
(D) Protein expression levels of G6PC-A204S in microsomal protein extracted from HEK293 cells was found to be downregulated by 41% compared to WT based on densitometric analysis (n=4), with representative western blot shown. Data presented as mean ± SEM and analysed using paired Students' t test. * p=0.01. Unt: Untransfected; WT: Wild type.

### Figure S5

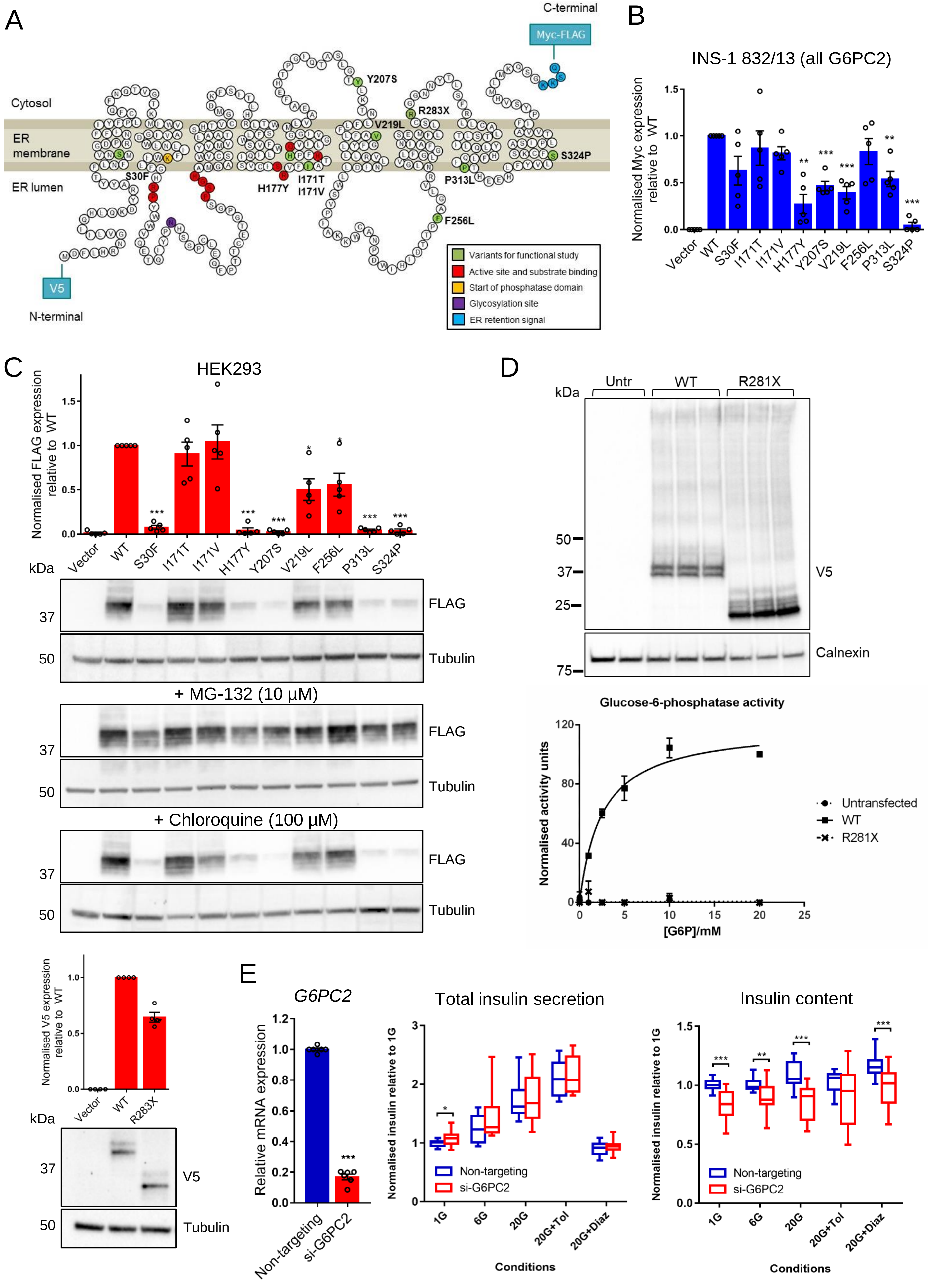

A
C-terminal
Myc-FLAG
Cytosol
ER membrane
ER lumen
V5
N-terminal
S30F
H177Y
I171T
I171V
Y207S
V219L
R283X
P313L
F256L
S324P
Variants for functional study
Active site and substrate binding
Start of phosphatase domain
Glycosylation site
ER retention signal
B
INS-1 832/13 (all G6PC2)
Normalised Myc expression relative to WT
Vector
WT
S30F
I171T
I171V
H177Y
Y207S
V219L
F256L
P313L
S324P
C
HEK293
Normalised FLAG expression relative to WT
kDa
FLAG
Tubulin
+ MG-132 (10 μM)
+ Chloroquine (100 μM)
Normalised V5 expression relative to WT
R283X
V5
D
Untr
WT
R281X
Calnexin
Glucose-6-phosphatase activity
Normalised activity units
[G6P]/mM
Untransfected
E
G6PC2
Relative mRNA expression
Non-targeting
si-G6PC2
Total insulin secretion
Normalised insulin relative to 1G
Conditions
1G
6G
20G
20G+Tol
20G+Diaz
Insulin content

**Figure S5. Functional characterisation of G6PC2 variants and the effect of *G6PC2* knockdown on insulin content and secretion in EndoC-βH1 cells.** Related to Figure 5.

(A) Variants prioritised for functional study in the context of the predicted G6PC2 protein structure (RefSeq NP_066999.1) in the ER membrane. Amino acid residues are coloured as described in the legend. Variants selected for functional study, in green, are labelled. The N-terminal V5 and C-terminal Myc-FLAG tags present in the expression constructs are indicated.

(B) Quantification of total G6PC2 variant protein expression (both upper and lower bands of representative western blot in Figure 5) in INS-1 832/13 cells based on western blot densitometric analysis of Myc-tagged G6PC2 constructs relative to tubulin control (n=5).

(C) Expression levels of G6PC2 variant proteins in HEK293 by western blot densitometric analysis of FLAG-tagged G6PC2 constructs or V5-tagged G6PC2-R283X relative to tubulin control (n=4). Representative blots are shown for untreated cells and cells treated with proteasomal inhibitor MG-132 or lysosomal inhibitor chloroquine.

(D) Glucose-6-phosphatase activity of the R281X variant in G6PC (proxy for R283X in G6PC2) in HEK293 (n=2), with representative western blot of microsomal protein shown.

(E) Total insulin secretion and insulin content were assessed at basal and high glucose conditions (with and without drug treatment) following 96-120h *G6PC2* knockdown in EndoC-βH1. Unpaired two-tailed Students' t tests were used to compare *G6PC2* knockdown to control for each condition, from n=16 across 4 independent experiments. Tol: tolbutamide; Diaz: diazoxide. All data presented as mean ± SEM. * $p=0.01-0.05$; ** $p=0.001-0.01$; *** $p<0.001$.

### Figure S6

A

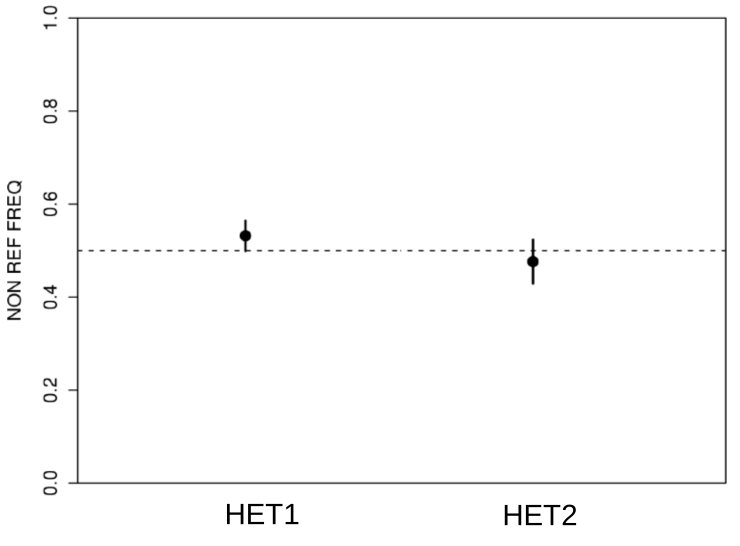


B

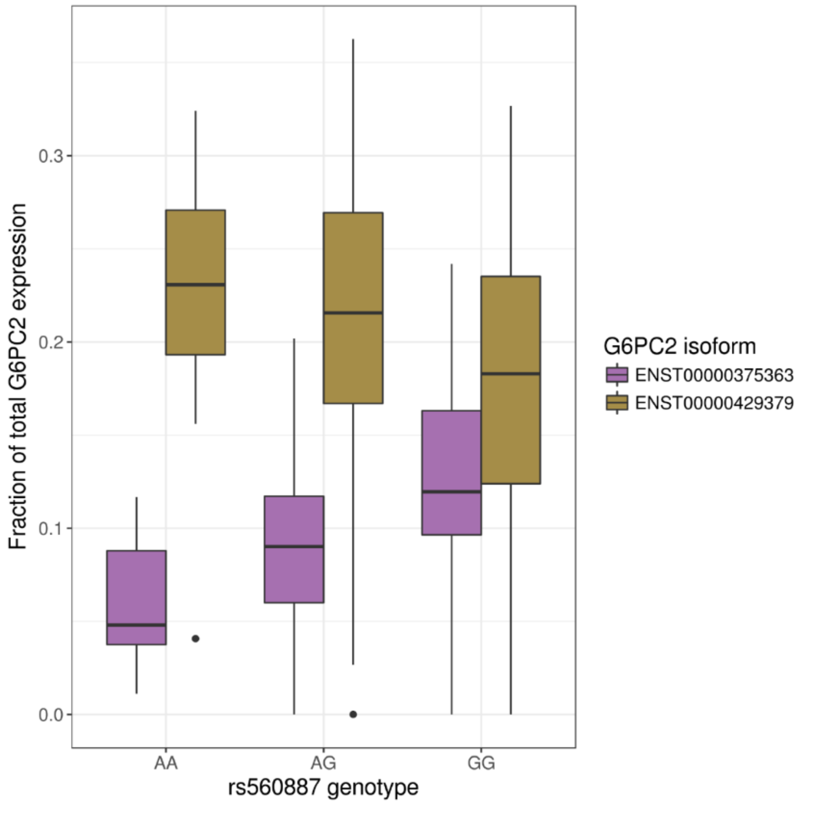


**Figure S6. *G6PC2* expression in RNA-Seq data from 150 human islet donor samples.** (A) Allelic balance was observed for *G6PC2* rs146779637 (p.R283X) in two heterozygote human islet samples.

(B) The glucose-raising rs560887-G allele associates significantly (*q*-value<0.01) with increased expression of the long *G6PC2* isoform (purple) and reduced expression of the short *G6PC2* isoform lacking exon 4 (brown).
